# Native Metabolomics Identifies the Rivulariapeptolide Family of Protease Inhibitors

**DOI:** 10.1101/2021.09.03.458897

**Authors:** Raphael Reher, Allegra T Aron, Pavla Fajtová, Paolo Stincone, Chenxi Liu, Ido Y Ben Shalom, Wout Bittremieux, Mingxun Wang, Marie L Matos-Hernandez, Kelsey L Alexander, Eduardo J Caro-Diaz, C Benjamin Naman, Chambers C. Hughes, Pieter C Dorrestein, Anthony J O’Donoghue, William H Gerwick, Daniel Petras

**Affiliations:** Scripps Institution of Oceanography, University of California San Diego, USA; Skaggs School of Pharmacy and Pharmaceutical Science, University of California San Diego, USA; Institute of Pharmacy, Martin-Luther-University Halle-Wittenberg, Halle (Saale), Germany; Department of Pharmaceutical Sciences, School of Pharmacy, University of Puerto Rico - Medical Sciences Campus, San Juan, Puerto Rico; Li Dak Sum Yip Yio Chin Kenneth Li Marine Biopharmaceutical Research Center, Department of Marine Pharmacy, College of Food and Pharmaceutical Sciences, Ningbo University 315832, Ningbo, China; Department of Chemistry and Biochemistry, University of California San Diego, USA; CMFI Cluster of Excellence, Interfaculty Institute of Microbiology and Infection Medicine, University of Tuebingen, Germany; Department of Microbial Bioactive Compounds, Interfaculty Institute for Microbiology and Infection Medicine, University of Tuebingen, Germany; German Center for Infection Research, Partner Site Tuebingen, Germany

**Author notes:** **Corresponding authors**, Correspondence should be addressed to Daniel Petras for questions regarding native metabolomics and MS-based structure elucidation, and to William Gerwick for question regarding cyanobacteria sampling and NMR-based structure elucidation.

## Abstract

The identity and biological activity of most metabolites still remain unknown. A key bottleneck in the full exploration of this tremendous source of new structures and pharmaceutical activities is the compound purification needed for bioactivity assignments of individual compounds and downstream structure elucidation. To enable bioactivity-focused compound identification from complex mixtures, we developed a scalable native metabolomics approach that integrates non-targeted liquid chromatography tandem mass spectrometry, and simultaneous detection of protein binding via native mass spectrometry. While screening for new protease inhibitors from an environmental cyanobacteria community, native metabolomics revealed 30 cyclodepsipeptides as chymotrypsin binders. Mass spectrometry-guided purification then allowed for the full structure elucidation of four new specialized metabolites via tandem mass spectrometry, chemical derivatization, and nuclear magnetic resonance spectroscopy. Together with the evaluation of biological activities, our results identified the rivulariapeptolides as a family of serine protease inhibitors with nanomolar potency, highlighting native metabolomics as promising approach for drug discovery, chemical ecology, and chemical biology studies.

## Main

Specialized metabolites, often referred to as natural products, are a tremendous pool of chemically diverse and pharmaceutically active organic compounds. By some estimates more than 50% of all current pharmaceuticals are based on or inspired by natural products^1^. Nevertheless, the vast majority of biological activities and pharmaceutical potential of specialized metabolites, as well as their ecological functions, still remain to be discovered^2^.

While mining genome and meta-genome data has begun to provide an overview of the biosynthetic potential of nature^3,4^, most specialized metabolites remain inaccessible, as the living organism that produces these metabolites cannot be cultured or their gene clusters remain silent under laboratory culturing conditions. Natural product discovery and chemical ecology studies in environmental samples are hence becoming more and more attractive^5^. Along with next-generation sequencing technologies, recent instrument and computational advances in nuclear magnetic resonance (NMR) spectroscopy and non-targeted liquid chromatography tandem mass spectrometry (LC-MS/MS) offer tremendous assistance to explore the uncharted metabolic space of nature^6^. These tools enable large-scale compound dereplication, rapid identification of chemical analogs, and *de novo* annotation of molecular formulas, substructures, chemical classes and structures^7–12^. However, the assignment of bioactivity of newly identified metabolites typically requires assays using pure compounds. Therefore, the isolation of specialized metabolites is typically guided by repetitive bioactivity assays. Together with full structure elucidation, this process usually takes months, and is therefore a major bottleneck for the systematic exploration of Nature for novel pharmaceutically active compounds and comprehensive chemical ecology studies.

A logical next step is the development of *activity metabolomics*^13^ or *functional metabolomics*^14^ approaches that aim to add functional information to the metabolites detected in a given system. Native electrospray ionization (ESI) and affinity mass spectrometry (MS) such as pulsed ultrafiltration (UF) MS, size exclusion (SEC) or affinity bead-based pull-down assays are increasingly being used to analyze non-covalent binding of biomolecules^15–23^. An important difference between native MS and affinity MS is that native MS detects ligands directly bound to a protein, whereas affinity MS approaches typically measure ligand binding indirectly as free compounds. For affinity MS, the target protein is captured by UF, SEC, centrifugation or magnetic removal and the released ligand is subsequently identified by small molecule MS analysis^24–27^. Both native and affinity MS approaches have been applied with single compounds as well as substrate pools, which allows for the simultaneous screening of thousands of compounds. An important inherent limitation in the use of ligand pools is that multiple ligands compete for binding of the target at the same site, and therefore compounds with highest affinity or concentration are more easily discovered. Additionally, the annotation of bound ligands remains a challenging task, especially if the ligand pool contains multiple isobaric compounds. While direct infusion native MS workflows have been developed that can identify metabolites bound to proteins by MS/MS^28^, combining the separation power of ultra-high-performance liquid chromatography (UHPLC) and the selectivity of native MS and MS/MS would offer a promising improvement to decipher protein-metabolite interactions out of complex biological mixtures, such as environmental samples. However, typical UHPLC mobile phase conditions disfavor non-covalent protein binding due to an acidic pH and high organic solvent content. To perform native MS coupled to UHPLC, we developed an experimental setup that increases pH and water content of the mobile phase post-column and infuses a protein binding partner before entering the ESI interface (**Figure 1**). As the protein is constantly infused post-column, one can monitor the intact protein mass over the LC-MS run and observe mass shifts when eluting metabolites bind to the protein at a defined retention time. Using collision induced dissociation (e.g., *Higher Energy C-Trap Dissociation* (HCD) in the setup), the complex can be dissociated again in the mass spectrometer and a “binding threshold” can be applied to distinguish between specific and non-specific binding. In combination with parallel non-targeted MS/MS analyses, the mass and compound ID or compound class can be assigned (level 2 or level 3 annotation^29,30^).

**Figure 1:**
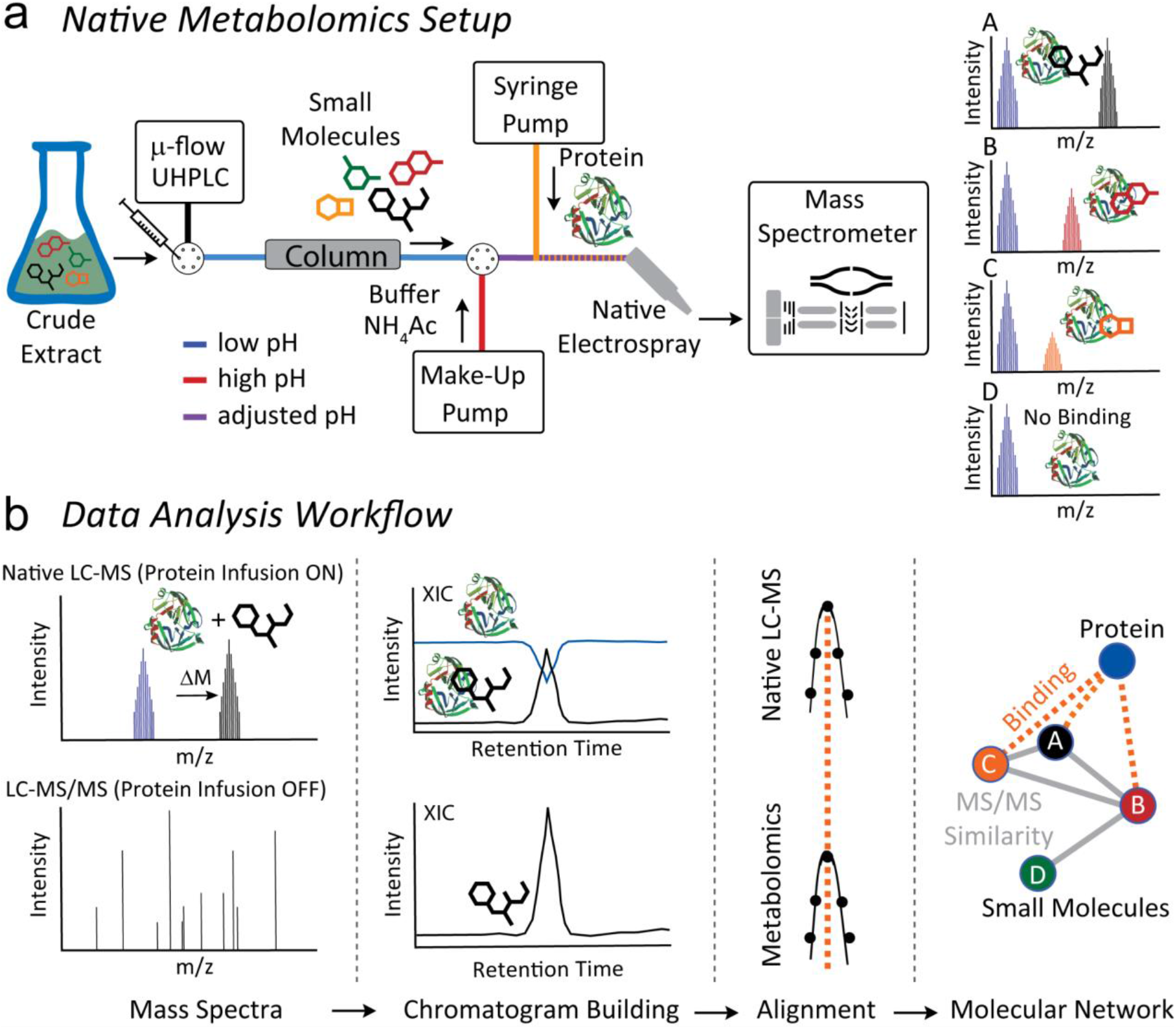
(a) Native metabolomics setup. A crude extract is separated by μ-flow UHPLC. The pH is adjusted after chromatography with ammonium acetate to “native-like” conditions via the make-up pump. Orthogonally, protein of interest is infused, and the resulting protein-binder complexes are measured by FT-MS. The procedure is repeated as a metabolomics run (high resolution UHPLC-MS/MS acquisition without protein infusion). For the data analysis (b) the □ *m/z* and retention time of the native MS run are correlated with *m/z* values and retention time of the metabolomics (LC-MS/MS) run and subsequently visualized using molecular networking and retention time pairing that links the observed mass differences of the protein in the native state vs bound states with the parent mass of MS/MS spectra of the small molecule.

After the successful proof-of-concept study, we screened for new protease inhibitors from an environmental cyanobacteria biofilm as a first application. In general, cyanobacteria have been a rich source of highly bioactive natural products^31–34^, and in particular protease inhibitors from numerous chemical classes^35–40^. Protease inhibitors are key compounds used for treatment of viral infections (SARS-CoV-2, HIV, and Hepatitis C)^41,42^, cancer^43^, diabetes^44^, hypertension^45^, and as general anticoagulants^46^. Several of the approved protease inhibitors are analogs of natural products such as aliskiren, captopril, and carfilzomib that target renin, angiotensin-converting enzyme, and proteasome, respectively^47^. In this study, we used chymotrypsin as the protease target to identify inhibitors from a marine cyanobacteria community. Using the native metabolomics approach, we identified 30 chymotrypsin binders in the methanolic crude extract with a single LC-MS run. The masses and MS/MS spectra of the binders were queried against structural and spectral databases, revealing that most of them were unknown. This led to the targeted isolation and structure elucidation of a family of new, and highly potent protease inhibitors, which we termed “rivulariapeptolides”.

## Results

### Development of the native metabolomics approach

In a crude extract, native metabolomics provides binding information about each compound towards a protein of interest. In the experimental setup, we utilized a single 10-minute LC-MS run to discover compounds that bind to the serine protease, chymotrypsin. The workflow is as follows: a crude extract is analyzed using native ESI while the protein of interest is infused post-column throughout the entire LC gradient. Binding of a small molecule to the protein of interest results in a peak with a mass corresponding to the protein bound to the compound. The *m/z* difference between the protein-ligand complex and the unbound protein reveals the molecular weight of the ligand while the ratio of the intensity of the protein-ligand peaks relative to the unbound protein peaks hints towards the relative binding affinity under the given conditions.

We first optimized the pH for native mass spectrometric acquisition of chymotrypsin and confirmed that the enzyme remains active under the native metabolomics buffer conditions. As a positive control for binding, we used molassamide^48^, a known non-covalent serine protease inhibitor of the 3-amino-6-hydroxy-2-piperidone (Ahp)-cyclodepsipeptide family. We found that an ammonium acetate buffer of pH 4.5 showed the highest peak intensity (**Figure S1a**). Next, we injected a serial dilution of molassamide into the native metabolomics LC-MS setup where it was mixed post-column with a constant concentration of chymotrypsin. The protein-ligand complex was detected at a deconvoluted mass of 26195.1 Da and the unbound apoprotein at 25232.6 Da (**Figure S1b**). The observed Δ *mass* of 962.5 Da matches the mass of molassamide (962.4749 Da). After deconvolution and integrating the peak area of the protein-ligand complex and plotting it against the molassamide concentration in the peak window, we obtained a binding curve. The resulting curve depicts a concentration-dependent increase of proteinligand to unbound protein ratio with increasing ligand concentration (**Figure S1c**). The limit of detection for the molassamide-chymotrypsin interaction was between 0.1 and 1 μg/mL (1 - 10 ng on column) (**Figure S1d**). To further test the biological relevance of the native metabolomics conditions, chymotrypsin was assayed with crude extract from the cyanobacterium *Rivularia* sp. using a fluorescence substrate competition assay. The bioassay conditions were designed to mimic the pH and solvent composition expected in the native mass spectrometry setup. Although chymotrypsin is optimally active in near-neutral pH, it retains good activity in 10 mM ammonium acetate at pH 4.5. Under these conditions, chymotrypsin was completely inhibited by 10 μg/mL of extract with 50% inhibition at 0.84 μg/mL (**Figure S1e**). Chymotrypsin was then assayed in increasing concentrations of acetonitrile (ACN) to determine if enzyme activity was retained in this solvent. Activity was reduced by 9% to 34% in ACN concentrations up to 33.3% v/v. In the presence of 41.7% v/v ACN, which corresponds to the end of the UHPLC gradient after the make-up addition, activity was decreased by 70% (**Figure S1f**). These results confirmed that chymotrypsin can be used as a target protease for native metabolomics as it retains activity at pH 4.5 in ACN concentrations up to 42% and binds to compounds from an inhibitory crude extract. To test the variability of the changing ACN concentration during the LC separation, we performed a series of flow injections (without column) over the full gradient (**Figure S2a**). The XIC of the molassamide bound chymotrypsin reveals similar signal responses throughout the gradient (5-99% ACN on column). While injecting a pool of control compounds at concentrations of 10 μg/mL, we observed compound specific signal responses of the chymotrypsin-ligand complexes (**Figure S2b**). To further assess the specificity of the protein-small molecule interactions, we evaluated binding of the linear oligopeptidic cysteine protease inhibitor gallinamide A, the cyclic depsipeptide FR900359, the isoflavone genistein, the phenol phloroglucinol, and the anthraquinone quinalizarin^33,49,50^. We did not observe binding of these negative controls to chymotrypsin under native MS conditions **(Figure S2c).**

### Native metabolomics reveals chymotrypsin binders

Following the successful proof-of-concept experiments, we next screened for potential chymotrypsin binders from a crude extract of a biofilm from the marine cyanobacterium *Rivularia* sp, collected from coral sediments at Carlos Rosario Beach in Culebra, Puerto Rico, U.S. The methanol extract was separated by reversed-phase UHPLC and ammonium acetate buffer and chymotrypsin were infused post-column, prior to native ESI and acquisition of mass spectrometry data in the high *m/z* range (2500-5000 *m/z*). The crude extract was subsequently re-injected, without infusion of chymotrypsin to obtain high-resolution LC-MS/MS data of compounds in the extract in the low *m/z* range (300-2000 *m/z*).

As a first step of data analysis, we plotted the total ion current (TIC) of the crude extract metabolomics data (**Figure 2a,** left) and extracted ion chromatogram (XIC) of the apo-chymotrypsin from charge state deconvoluted native mass spectrometry data (**Figure 2a,** right) which shows several negative peaks in the range of 4.5 - 5.5 minutes. The decrease in signal of the apo-protein in that retention time range is due to the emergence of larger masses that correspond to protein-ligand complexes. After feature finding of the deconvoluted masses and matching of the parallel metabolomics LC-MS/MS data of the crude extracts by retention time and exact mass matching, we could identify more than 30 potential small molecule-protein complexes. To display the family of small molecules that form protein ligand complexes and to show their structural relations, we visualized them in a correlation molecular network (**Figure 2b**) that is based on their MS/MS similarity (grey line), retention time, and mass matching between protein and small molecules through the red dashed lines.

**Figure 2:**
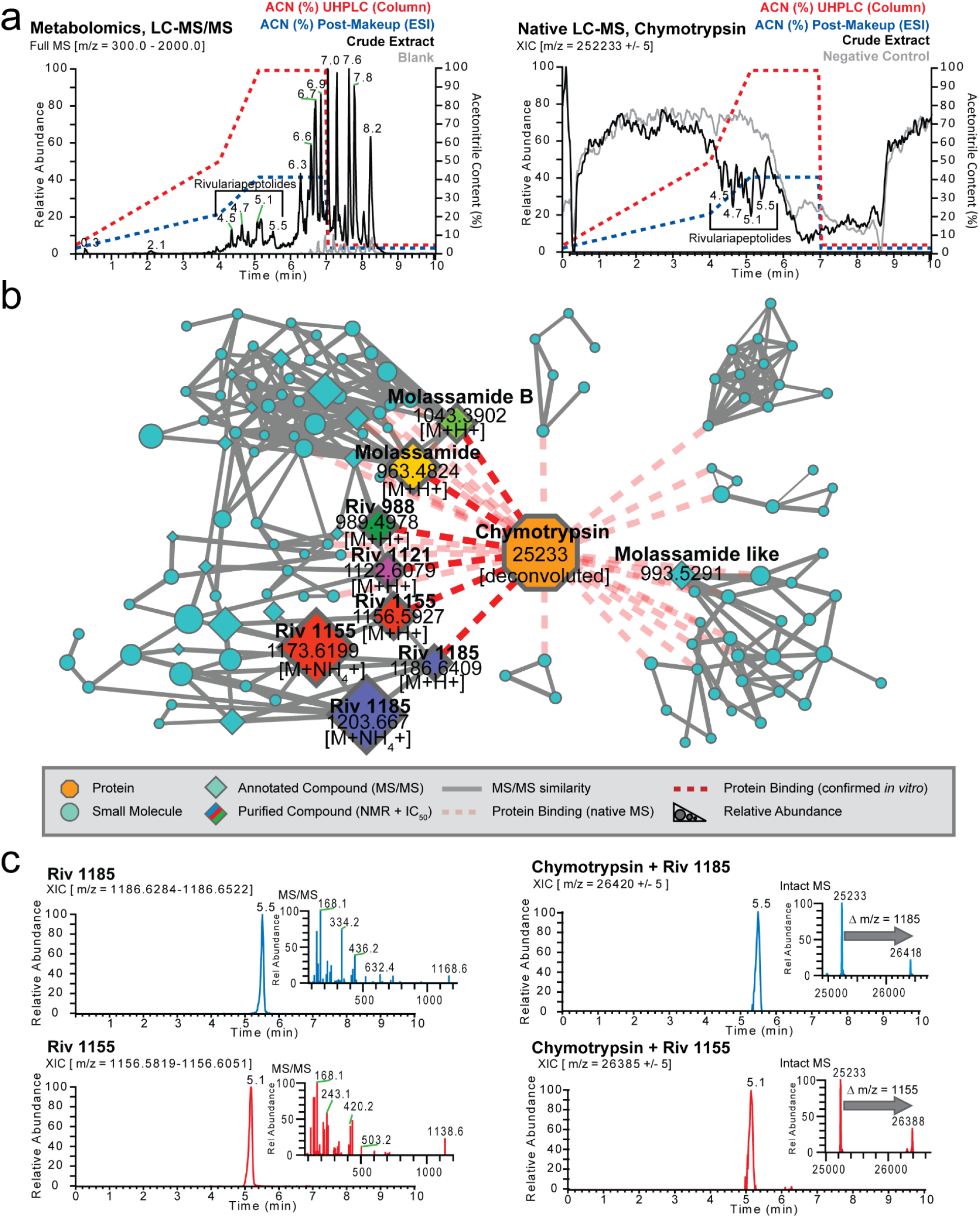
(a) Left panel: Full mass spectrum (*m/z* 300 - 2,000) of cyanobacterial crude extract and blank. The retention times (RT = 4.5 - 5.5 min) of major Ahp-cyclodepsipeptides are highlighted. Right panel: Deconvoluted extracted ion chromatogram (*m/z* 25,233 ± 5) of alpha-chymotrypsin screened against the cyanobacterial crude extract under native MS conditions and negative control. Acetonitrile (ACN) concentration on column and post-column (including make-up) are shown as dashed lines. The ACN concentrations are given at pump for a given time and a ~ 2 min delay time between pump and column has to be taken into account. (b) Correlation molecular network of deconvoluted chymotrypsin and putatively new small molecule inhibitors binders by native MS (A larger version of the network with detailed precursor mass labels of all nodes is available in the supporting information Figure S3). (c) Upper left panel: Extracted ion chromatogram (*m/z* 1,186.6284 - 1,186.6522) and MS^2^ spectrum of putative new chymotrypsin-binder rivulariapeptolide 1185 at RT = 5.5 min. Upper right panel: Extracted ion chromatogram (*m/z* 26,420 ± 5) of a putative chymotrypsin-binder complex. Mass difference between the putative chymotrypsin-binder complex (*m/z* 26,420 ± 5) and apo-chymotrypsin (25,233 ± 5) suggests a molecular weight of 1,187 ± 5 Da for the putative chymotrypsin binder rivulariapeptolide 1185. Lower left panel: Extracted ion chromatogram (*m/z* 1,156.5819 - 1,156.6051) and MS^2^ spectrum of putative chymotrypsin-binder rivulariapeptolide 1185 at RT = 5.1 min. Lower right panel: Extracted ion chromatogram (*m/z* 26,385 ± 5) of a putative chymotrypsin-binder complex. Mass difference between the putative chymotrypsin-binder complex (*m/z* 26,385 ± 5) and apo-chymotrypsin (25,233 ± 5) suggests a molecular weight of 1,152 ± 5 Da for the putative chymotrypsin binder rivulariapeptolide 1155.

Two of the most abundant being *m/z* 1186.6400 and *m/z* 1156.5923 that also show perfect overlap of the chromatographic profiles between intact protein and metabolomics LC-MS/MS data (**Figure 2c**). Based on their high relative abundance we targeted them for further purification, structure determination by NMR and orthogonal protease inhibition assays.

### Rivulariapeptolides, a family of new Ahp-cyclodepsipeptides

The potential chymotrypsin binder with the *m/z* 1186.6400, identified by native metabolomics, was next targeted for isolation and structure elucidation, using state-of-the-art high-resolution MS/MS and NMR approaches^51^. We first separated the *Rivularia* crude extract into four fractions of decreasing polarity via solid phase extraction. SMART NMR^11^ analysis was applied to the most hydrophilic fraction and all but one structure of the top 10 SMART results were predicted as cyclic depsipeptides (**Figures 3a/b, S4**), including 6 of the top 10 as Ahp-cyclodepsipeptides (three from marine filamentous cyanobacteria: somamide B, molassamide, lyngbyastatin 6)^48,52–54^. Complementarily, MS/MS-based molecular networking analysis of the *Rivularia* crude fractions assisted with the annotation of the known Ahp-peptides molassamide, kurahamide, and loggerpeptin A along with several putatively new ones (**Figure 3a/b, S5a**)^48,53,55^. Next, the SIRIUS and ZODIAC tools^56,57^ were applied to determine the molecular formula of the chymotrypsin-binding feature, with exact mass *m/z* 1186.6400 [M+H]^+^, as C_61_H_87_N_9_O_15_ (0.5 ppm). Subsequently, we classified the MS/MS spectrum indicative for a ‘cyclic depsipeptide’ based on the classification with CANOPUS^9^. Further substructures of the molecule were predicted as benzene, hydroxy-benzene, and proline/*N*-acyl-pyrrolidine derivatives, as well as piperidinone/delta-lactam for the Ahp-family defining moiety (**Figures 3c, S5b**).

**Figure 3:**
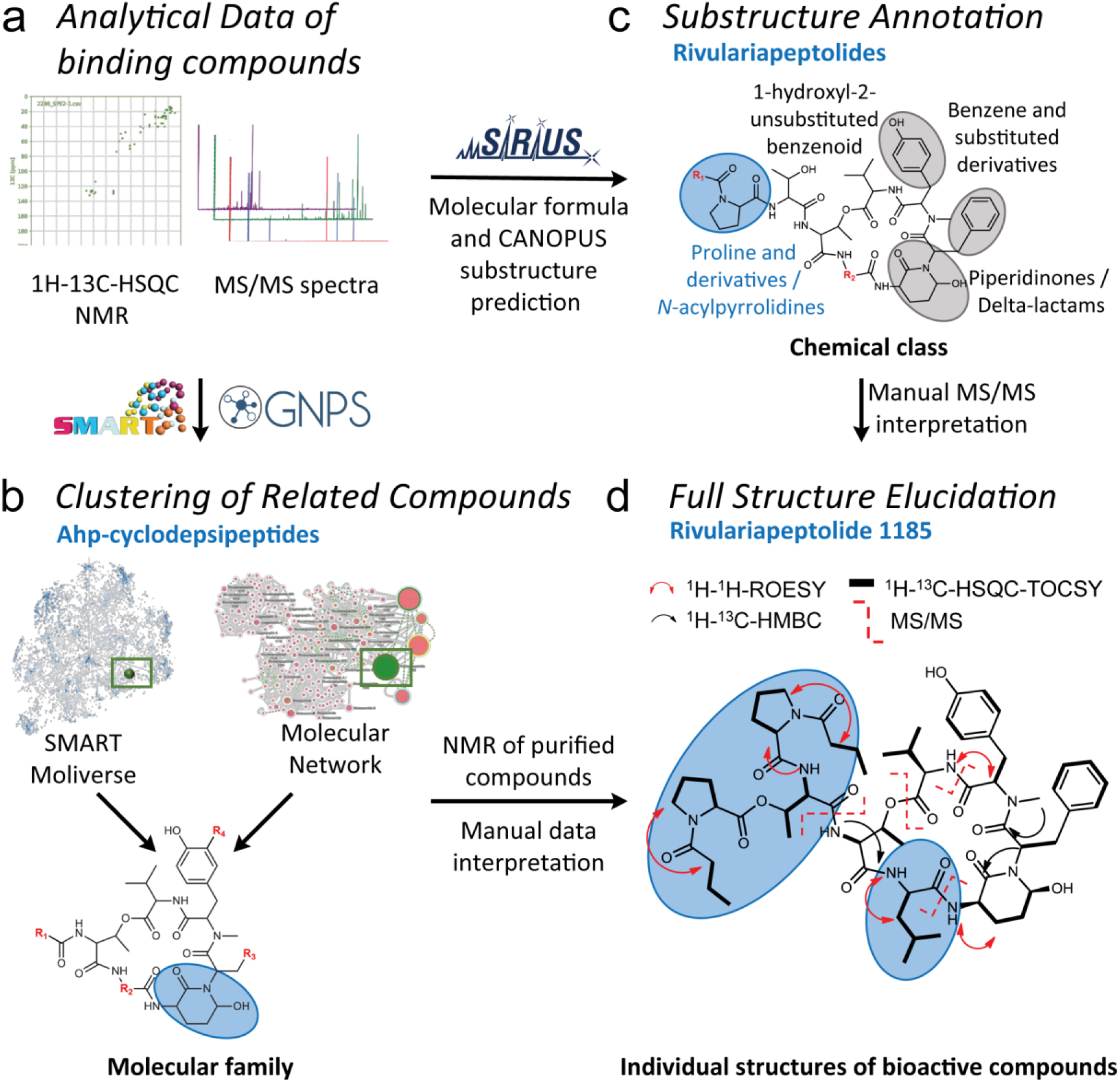
Structure elucidation workflow based on NMR and MS/MS data (a). The workflow combined automated *in-silico* MS/MS and NMR annotation tools for fast compound class identification and dereplication of known natural products exemplified by the new natural product rivulariapeptolide 1185. (b) Molecular networking and SMART analysis suggested the presence of a Ahp-cyclodepsipeptide molecular family (c) In depth MS and MS/MS analysis of the new natural products with SIRIUS helped to establish the molecular formula and substructural information about the characteristic *N*-acylated proline residues. (d) Unambiguous structure elucidation by various 1D/2D NMR and MS/MS experiments led to the planar structure of rivulariapeptolide 1185. Selected 2D NMR-derived correlations and MS^2^ fragmentations are depicted. The distinctive structural moieties are highlighted in grey and blue bubbles.

To unambiguously determine the structure, we isolated *m/z* 1186.6400, named rivulariapeptolide 1185 (**1**), and performed 1D/2D NMR experiments and manual MS/MS interpretation (**Figure 3d, Figures S6-S11, S41, Table S1**). Subsequently, we targeted the isolation of further rivulariapeptolides by preparative HPLC, based on their protein-ligand complex ratios from the native metabolomics experiments as well as their relative abundance. In that way, we isolated and elucidated the planar structures of the rivulariapeptolides 1185, 1155, 1121, and 989 (**1, 2, 3, 4**) with the exact masses 1186.6400 [M+H]^+^ (C_61_H_88_N_9_O_15_, 0.5 ppm), 1156.5923 [M+H]^+^ (C_59_H_82_N_9_O_15_, - 0.2 ppm), 1122.6080 [M+H]^+^ (C_56_H_84_N_9_O_15_, - 0.1 ppm) and 989.4978 [M+H]^+^ (C_50_H_69_N_8_O_13_, - 0.1 ppm). In addition to rivulariapeptolides, we identified the already known molassamide (**5**) as well as new derivative of the latter that we termed “molassamide B” (**6**) with *m/z* 1041.3924 [M+H]^+^ (C_48_H_66_BrN_8_O_13_, −0.3 ppm), which is ortho-brominated (**Figures S12-S41, Tables S2-S3**). The absolute configurations of the amino acids were determined by UHPLC-MS analysis of the acid hydrolysates of **2** and its pyridinium dichromate oxidation product and subsequent advanced Marfey’s analysis (**Figure S42a**). The analyses revealed L-configurations for all amino acids as is the case for other cyanobacterial Ahp-cyclodepsipeptides. The relative configuration of the stereocenters of the (3S, 6R)-Ahp unit as well as the geometry of the double bond (Z-configuration) of the 2-amino-2-butenoic acid (Abu) moiety was determined by NOESY and HMBC NMR experiments (**Figure S42b**). Assuming that the other rivulariapeptolides (**1, 3, 4**) and molassamides (**5**+**6**) described here originate from the same biosynthetic peptide synthetase, they most probably also share the same backbone configuration. The highly comparable MS/MS and NMR data sets of compounds **1**–**6** (**Figures S6-S41**, **Tables S1-S3**) provide additional evidence that the discovered Ahp-cyclodepsipeptides from this study share the same configuration.

Finally, the chymotrypsin inhibitory activities of the purified Ahp-cyclodepsipeptides **1-6** were assessed by specific biochemical assays and confirmed the results of the native metabolomic protein infusion MS experiments (**Figure 4**). All six compounds were found to be nanomolar chymotrypsin inhibitors with compound **1** being the most potent (**Figure 4**, IC_50_ = 13.17 ± SD nM). The newly described family of rivulariapeptolides is characterized by a rare (duplicated) *N*-butyrylated proline moiety in the side chain. All for the first time described peptides **1-4** and **6** reside among the six most potent chymotrypsin inhibitors, so far reported from the compound class of Ahp-cyclodepsipeptides (**Table S4**).

**Figure 4:**
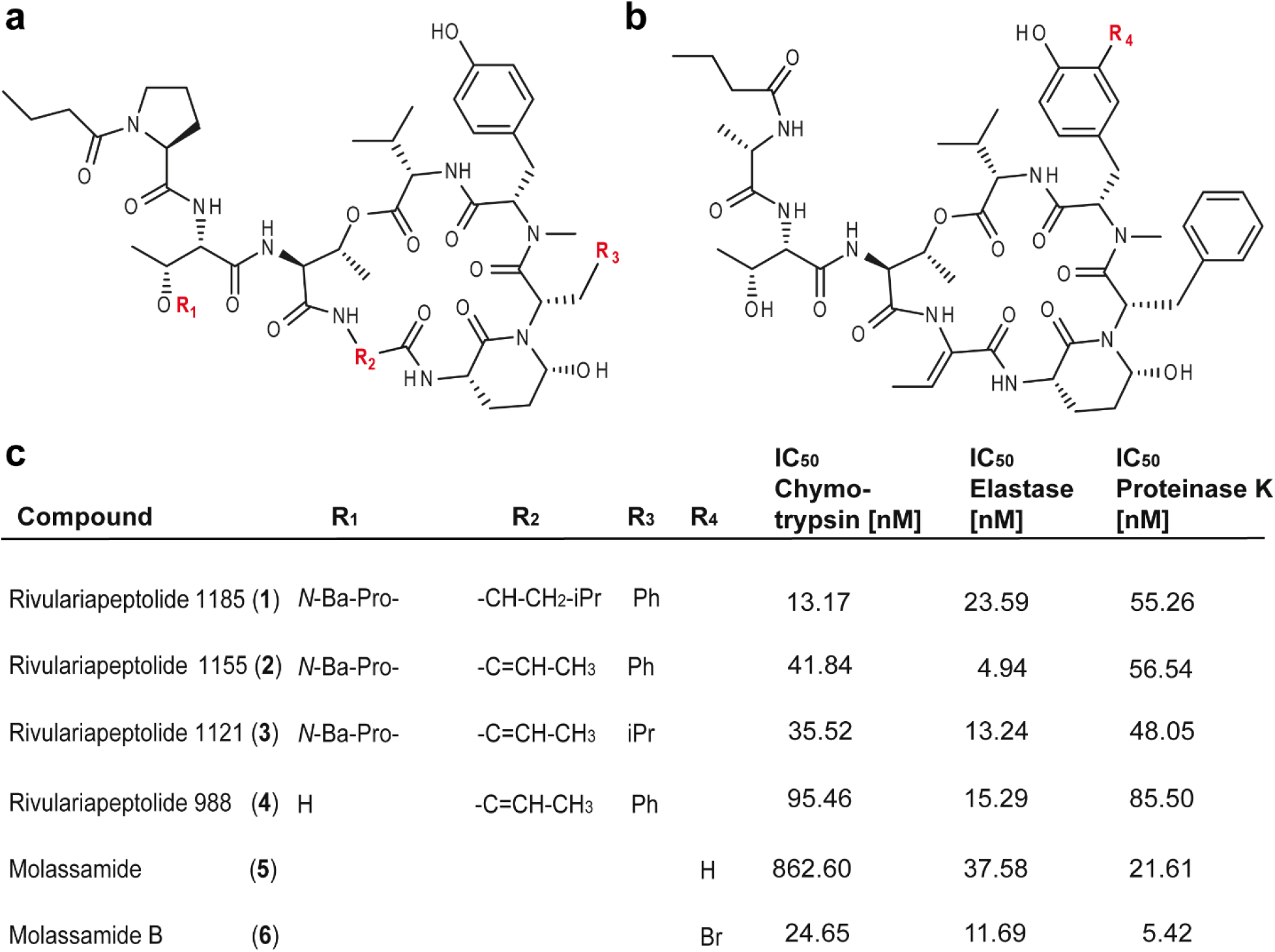
(a) Structures of the isolated rivulariapeptolides 1185 (**1**), 1155 (**2**), 1121 (**3**), 988 (**4**). (b) Structures of the isolated known molassamide (**5**), and the new molassamide B (**6**). (b) Potency of isolated compounds for selected serine proteases following 40 min pre-incubations. Data are presented as the mean ± SD, n = 3. Abbreviations: Ba = butyric acid, Pro = proline.

Intriguingly, a single ortho-bromination in the *N*-methyltyrosine moiety led to a 35-fold increase in potency for the new compound **6** (IC_50_ = 24.65 ± SD nM), when compared to the known compound **5** (IC_50_ = 862.60 ± SD nM). These promising results led us to test these six Ahp-cyclodepsipeptides against two other serine proteases, elastase and proteinase K. Elastase is produced in either the pancreas, for digestion of food, or by neutrophils for degradation of foreign proteins. Neutrophil elastase is a well-established drug target for treatment of acute lung injury and acute respiratory distress syndrome^58^. Proteinase K is a fungal serine endopeptidase that is commonly used in molecular biology procedures^59^. This enzyme family are play important roles in fungal infection of insects^60^ and mammals^61^. While **2** was found to be a potent elastase inhibitor (IC_50_ = 4.94 ± SD nM), **6** was discovered to be the most potent proteinase K inhibitor known to date (IC_50_ = 5.42 ± SD nM). The isolated compounds were docked by induced-fit, inside the binding pocket of alpha-chymotrypsin (PDBID 4Q2K) and all were found to have a similar binding mode (**Figure S43**) that was revealed by crystal structures of Ahp-cyclodepsipeptides in complex with serine proteases^62,63^.

## Discussion

Here we describe the use of native metabolomics protein infusion MS to simultaneously detect protein-metabolite binding and annotate their molecular structures. This approach can be used for rapid screening of small molecule modulators for proteins of interest, directly from crude extracts. In our case study, we identified 30 chymotrypsin binding natural products in a 10 min LC-MS run from a few μg of crude extract (not including downstream isolation and *de novo* structure elucidation). In comparison to flow injection experiments (injection of crude extract in the native MS setup without column) we observed strong signal decrease, most likely through collective ion suppression effects (**Figure S2d,** left panel). When injecting preincubated chymotrypsin-crude extract mixture into our system (**Figure S2d,** right panel) we observed more acceptable signal response and several protein-metabolite complexes, which we attributed to molassamide B, a molassamide derivative and rivulariapeptolide 1185. However, in comparison to native metabolomics, the number of putative binders observed was significantly lower, indicating that chromatographic separation is an important factor for the sensitivity of the approach. Most importantly, the chromatographic dimension is also essential for the unambiguous linking of binding information to MS/MS features which facilitates dereplication and downstream structure elucidation.

Native metabolomics is generalizable to other protein targets that are accessible via native ESI^64–66^ and which are available in large enough quantities. The amounts of protein needed for native metabolomics high-throughput screening are a few mg for a 24 h screen of 96 samples with a 10 min LC-MS method, which is achievable for many commercially available proteins or in-house heterologous protein expression.

Besides the ad-hoc experimental determination of binding information, we systematically organized this information in the public GNPS spectral library through spectrum tags. Hence, native metabolomics derived properties are accessible in future experiments and can provide biological context to complex metabolomes.

While we primarily used the method for an initial screening approach and assigned binary binding information (binder/non-binder), titration experiments using native mass spectrometry can be used to determine relative dissociation constants (K_d_) by fitting the intensity ratio of bound and unbound protein as a function of the added ligand. This method first assumes that no dissociation takes place during the transmission through the mass spectrometer, and second, that the observed gas phase intensity ratio correlates with the solution ratio. These assumptions imply that ESI titration measurements can deliver a relative “snapshot” of the solution concentrations to reflect solution-phase binding affinities. Nevertheless, as the experimental environment is inherently different (gas phase vs. solution) absolute binding affinities might differ from solution based orthogonal assays^67^.

From a natural product discovery perspective, it is very interesting that 30 putative bioactive molecules for a target protein were discovered from a single extract. This indicates that the chemical space for certain bioactive molecular families can often be underestimated when compared to traditional bioactivity-guided approaches, as they are typically biased towards the most abundant or most active compounds. At least for Ahp-cyclodepsipeptides, recent biosynthetic studies suggest that the high structural diversity of these compounds is mainly driven by the hypervariability of amino acids in the positions proceeding and following the Ahp-unit (see **Figure 4a** for the definitions of residues R_3_ and R_2_, respectively)^68^. The events impacting R_2_ can be explained by high-frequency point mutations. This is sought to provide an evolutionary platform to iteratively test combinations while maintaining the central activity. However, the amino acid substitutions at R_3_ most likely occur via recombination events, thereby allowing for evolutionary shortcuts^68^. These biosynthetic hypotheses are supported by the compounds isolated in this study and add to a better understanding of structure-activity relationships for Ahp-cyclodepsipeptides. Comparing rivulariapeptolide 1185 (**1**) to rivulariapeptolide 1121 (**3**), a leucine residue is swapped for an Abu unit at R_2_, and phenylalanine is replaced by leucine at R_3_. The most surprising structure-activity relationship gained from this study, however, was that a single substitution of bromine (molassamide B, **6**) for hydrogen (molassamide, **5**) led to a thirtyfive-, three-, and four-fold increase in potency towards chymotrypsin, elastase, and proteinase K, respectively.

The protease inhibition of the compounds discovered with the native metabolomics workflow was confirmed with an orthogonal fluorescence assay against three proteases. At nanomolar concentration their IC_50_ show high potency and exhibit distinct selectivity (**Figure 4, Table S4**). For example, molassamide (**5**) is the second most potent inhibitor screened against proteinase K but is the least potent inhibitor for chymotrypsin and elastase. Rivulariapeptolide 1155 (**2**), on the other hand, is the most potent elastase inhibitor but shows much lower inhibition against both chymotrypsin and proteinase K.

Together, these findings highlight the utility of native metabolomics approach presented herein. Beyond the discovery of novel protease inhibitors, we anticipate that native metabolomics will be applied for the screening of a broad variety of interactions of biomolecules from complex mixtures at scale. We anticipate that native metabolomics will be used as a central tool for activity/ functional metabolomics workflows. This will not only benefit drug discovery and chemical ecology studies but could also be leveraged for the generation of large-scale training data for machine learning approaches to predict protein-ligand interaction.

## Methods

### Cyanobacterial collection and taxonomy

Marine cyanobacteria biofilm samples were collected in an intertidal zone growing on rock/reef substrate near Las Palmas Beach, Manatí, Puerto Rico, U.S. (GPS coordinates: 18°28’32.0”N 66°30’00.5”W) on May 14th, 2019, and at 0.5 - 2.0 m of water at Carlos Rosario Beach in Culebra, Puerto Rico, U.S (GPS coordinates: 18°19’30.0”N 65°19’48.0”W) on April 6th, 2019. Biomass for both samples was hand collected (DRNA Permit O-VS-PVS15-SJ-01165-15102020). Microscopic examination indicated that this collection was morphologically consistent with the genus *Rivularia*. 16S rDNA analysis confirmed the identity as *Rivularia* spp. PCC 7116. Voucher specimen available from E.C.D. as collection no. MAP14MAY19-1, and from W. H. G. as collection no. CUR6APR19-1.

### Micro-flow LC-MS/MS data acquisition

For micro-flow UHPLC-MS/MS analysis 2 μL were injected into vanquish UHPLC system coupled to a Q-Exactive (setup A) or a Q-Exactive HF (setup B) quadrupole orbitrap mass spectrometer (Thermo Fisher Scientific, Bremen, Germany) with an Agilent 1260 quaternary HPLC pump (Agilent, Santa Clara, USA) or in setup B a fully integrated vanquish quaternary UHPLC pump (Thermo Fisher Scientific, Bremen, Germany) as make-up pumps. For reversed-phase chromatographic, a C18 core-shell microflow column (Kinetex C18, 150 x 1 mm, 1.8 um particle size, 100 A pore size, Phenomenex, Torrance, USA) was used. The mobile phase consisted of solvent A (H_2_O + 0.1 % formic acid (FA)) and solvent B (acetonitrile (ACN) + 0.1 % FA). The flow rate was set to 150 μL/min (setup A) or 100 μL/min (setup B). In setup A, a linear gradient from 5-50 % B between 0-4 min and 50-99 % B between 4 and 5 min, followed by a 2 min washout phase at 99% B and a 3 min re-equilibration phase at 5 % B. In setup B, a linear gradient from 5-50 % B between 0-8 min and 50-99 % B between 8 and 10 min, followed by a 3 min washout phase at 99% B and a 5 min re-equilibration phase at 5 % B. Data-dependent acquisition (DDA) of MS/MS spectra was performed in positive mode. Electrospray ionization (ESI) parameters were set to 40 arbitrary units (AU) sheath gas flow, auxiliary gas flow was set to 10 AU and sweep gas flow was set to 0 AU. Auxiliary gas temperature was set to 400 °C. The spray voltage was set to 3.5 kV and the inlet capillary was heated to 320 °C. S-lens level was set to 70 V applied. MS scan range was set to 200-2000 *m/z* with a resolution at *m/z* 200 (R_*m/z* 200_) of 70,000 with one micro-scan. The maximum ion injection time was set to 100 ms with automatic gain control (AGC) target of 5E5. Up to two MS/MS spectra per duty cycle were acquired at *R*_*m/z* 200_ 17,000 with one micro-scan. The maximum ion injection time for MS/MS scans was set to 100 ms with an AGC target of 5.0E5 ions and a minimum 5% AGC. The MS/MS precursor isolation window was set to *m/z* 1. The normalized collision energy was stepped from 20 to 30 to 40% with z = 1 as the default charge state. MS/MS scans were triggered at the apex of chromatographic peaks within 2 to 15 s from their first occurrence. Dynamic precursor exclusion was set to 5 s. Ions with unassigned charge states were excluded from MS/MS acquisition as well as isotope peaks. For native metabolomics experiments, the same chromatographic parameters were used and in addition 220 μL/min (setup A) or 150 μL/min (setup B) 10 mM ammonium acetate buffer was infused post-column through a make-up pump and a PEEKT-splitter and enzyme solution was infused with 2 μL/min flow rate via the integrated syringe pump. ESI settings were set to 40 arbitrary units (AU) sheath gas flow, auxiliary gas flow was set to 10 AU and sweep gas flow was set to 0 AU. Auxiliary gas temperature was set to 400 °C. The spray voltage was set to 3.5 kV and the inlet capillary was heated to 300 °C. S-lens level was set to 80 V applied. MS scan range was set to 2000-4000 *m/z* with a resolution *R*_*m/z* 200_ 140,000 with 2 microscans. MS acquisition was performed in all ion fragmentation (AIF) mode with *R*_*m/z* 200_ with 20% HCD collision energy and an isolation window of 2000 - 4000 *m/z* (setup A) or 2500 - 4000 *m/z* (setup B).

### Native metabolomics data analysis

For native LC-MS data, multiple charged spectra were deconvoluted using the xtract algorithm in Qualbrowser, part of the Xcalibur software (Thermo Scientific). Both deconvoluted native LC-MS and metabolomics LC-MS/MS .raw files were converted to centroid .mzML file format using MSconvert of the proteowizard software package. Feature finding of both file types was performed using a modified version of MZmine2.37 (corr.17.7). Feature tables from both intact protein mass and metabolomics data were matched by their retention time (RT) and an *m/z* offset corresponding to the mass of chymotrypsin (25234 Da) with an RT tolerance of 0.2 min and a mass tolerance of 4 Da. Feature tables (.csv), MS/MS spectra files (.mgf), and ion identity networking results (.csv) were exported and uploaded to the MassIVE repository. LC-MS/MS data was submitted to GNPS for feature-based molecular networking analysis. Downstream combined Molecular-Networks and chymotrypsin small molecule binding were visualized as networks in cytoscape (3.8.2).

### pH dependency of native metabolomics

pH values over the entire LC gradient and peak intensities were both assessed using three different make-up solvents. Make-up solvent A was water, make-up solvent B was 10 mM ammonium acetate buffer, and make-up solvent C was 10 mM ammonium acetate buffer + 0.2% ammonium hydroxide, v/v. Molassamide was prepared as 100 μM solutions from a 10 mM stock solution in DMSO by preparing a 1:100 dilution into solvent mixture 1 (water + 10% acetonitrile + 0.1% formic acid). Samples were analyzed as described in micro-flow LC-MS/MS data acquisition (setup A); 2 μL of each solution were injected into the mass spectrometer, while chymotrypsin (Sigma), dissolved in water to a final concentration of 2 mg/mL, was injected through the syringe pump at a flow rate of 2 μL/min. pH values were assessed by disconnecting the flow to the source and collecting ~30 μL of solvent every minute then testing this value on pH paper.

### Titration of ligands and concentration dependency

Molassamide, Riv 1155, and Riv 1185 were prepared as 100 μM solutions from a 10 mM stock solution in DMSO by preparing a 1:100 dilution into solvent mixture 1 (water + 10% acetonitrile + 0.1% formic acid). From this 100 μM solution, dilutions were prepared at 10 μM, 1 μM, and 0.1 μM into solvent mixture 1.2 μL of each solution were injected into the mass spectrometer, then 5 μL and 10 μL of the 100 μM solution were injected to yield final concentrations of 250 μM and 500 μM, respectively. Samples were analyzed as described in Micro-Flow LC-MS/MS data acquisition (setup A), while chymotrypsin (Sigma Aldrich) was dissolved in water to a final concentration of 2 mg/mL and injected through the syringe pump at a flow rate of 2 μL/min. The ratio of bound to unbound protein was plotted against the ligand concentration in a given HPLC peak window. Data points were fitted using the solver function.

### Determination of limit of detection, selectivity, and flow-injection experiments

Molassamide was dissolved in 50% MeOH and a serial dilution with a dilution factor of 2 and 10 where performed yielding final concentrations of 100, 50, 10, 1, and 0.1 μg/mL. 2 μL of each dilution where injected into the native metabolomics microflow LC-MS system (setup B) and peak areas from chymotrypsin bound molassamide where extracted and plotted against their concentration. For testing the selectivity of the method, a series of standards (gallinamide A, FR900359, quinalizarin, genistein, phloroglucinol, cymodepsipeptide A, lingaoamide, molassamide B, rivulariapeptolide 1121, rivulariapeptolide 1185, tutuilamide A) where dissolved to 100 μg/mL and pooled to final concentrations of 10 μg/mL in 50% MeOH. For flow injection analysis the 10 μg/mL molassamide standard was used and the UHPLC column was bypassed with a stainless steel union. Continuous injections during the microflow LC gradient were performed manually through the direct control function in the Xcalibur software (Thermo Scientific) with ~ 1 min spacing.

### Chymotrypsin activity assays in native mass spectrometry buffer

Cyanobacteria extract (1 mg/ml, methanol) was diluted in 10 mM ammonium acetate pH 4.5 to 30 ug/mL and then sequentially diluted 1.5-fold to 0.52 μg/mL in the same buffer. Bovine chymotrypsin (Sigma Aldrich) and Suc-Ala-Ala-Pro-Phe-AMC (Calbiochem, 230914) were diluted to 300 nM and 150 μM, respectively in 10 mM ammonium acetate pH 4.5. In a 384-well black microplate, 10 μL of enzyme, substrate and cyanobacteria extract were combined (30 μL final volume) such that the concentrations in the reaction were 100 nM chymotrypsin, 50 μM of Suc-Ala-Ala-Pro-Phe-AMC and 10 μM to 0.17 nM of cyanobacteria extract. For solvent compatibility assays, chymotrypsin (100 nM) was assayed with 10 μg/mL and 1 μg/mL extract in 10 mM ammonium acetate, pH 4.5 containing 8.3 to 41.7% acetonitrile. A control assay lacked acetonitrile and cyanobacteria extract. All assays were performed in triplicate wells at 25°C in a Synergy HTX Multi-Mode Microplate Reader (BioTek, Winooski, VT) with excitation and emission wavelengths of 360 and 460 nm, respectively. The initial reaction velocity in each well was recorded and the dose response curve was generated using GraphPad Prism 9 software.

### Calculation of IC_50_ values

Chymotrypsin (1 nM), proteinase K (10 nM), and elastase (20 nM) was preincubated with 0 to 3 μM of each compound for 40 min in Dulbecco’s phosphate buffered saline, pH 7.4 containing 0.01% Tween-20. The reaction was initiated by addition of 25 μM of Suc-Ala-Ala-Pro-Phe-AMC (Calbiochem, 230914) for proteinase K and chymotrypsin, 25 μM of MeOSuc-Ala-Ala-Pro-Val-AMC (Cayman, 14907) for elastase in a final volume of 30 μL, respectively. The release of the AMC fluorophore was recorded in a Synergy HTX multi-mode reader (BioTek Instruments, Winooski, VT) with excitation and emission wavelengths at 340 nm and 460 nm, respectively. The maximum velocity was calculated in RFU/sec over 10 sequential points on the linear part of the progress curve. The IC50 values were determined by nonlinear regression in GraphPad Prism 9.

### NMR spectroscopy

Deuterated NMR solvents were purchased from Cambridge Isotope Laboratories. 1H NMR and 2D NMR spectra were collected on a Bruker Avance III DRX-600 NMR with a 1.7 mm dual tune TCI cryoprobe (600 and 150 MHz for 1H and 13C NMR, respectively) and a JEOL ECZ 500 NMR spectrometer equipped with a 3 mm inverse detection probe. NMR spectra were referenced to residual solvent DMSO signals (δH 2.50 and δC 39.5 as internal standards). The NMR spectra were processed using MestReNova (Mnova 12.0, Mestrelab Research) or TopSpin 3.0 (Bruker Biospin) software.

### Extraction and isolation

The preserved cyanobacterial biomass from collection no. CUR6APR19-1 was filtered through cheesecloth, and then (98.7 g dry wt) was extracted repeatedly by soaking in 500 mL of 2:1 CH_2_Cl_2_ /MeOH with warming (<30 °C) for 30 min to afford 1.44 g of dried extract. A portion of the extract was fractionated by reverse-phase solid phase extraction (C_18_-SPE) using a stepwise gradient solvent system of decreasing polarity (Fr. 1-1 35% ACN/H_2_O, 124.4 mg; Fr. 1-2 70% ACN/H_2_O, 76.1 mg; Fr. 1-3 100% ACN, 77 mg; Fr. 1-4 100% MeOH, 254.9 mg. Fr. 1-2 was dissolved in 70% ACN/H2O and purified by preparative HPLC using a Kinetex 5 μm RP 100 Å column (21.00 × 150mm) and isocratic elution using 50% ACN/H2O for 8 minutes then ramping up to 100% in 14 minutes at the flow rate of 20 mL/min, yielding 56 subfractions. Rivulariapeptolide 1185 (compound **1**) and 1155 (compound **2**) were isolated from subfractions 1-2-20 to 1-2-23 that were combined (4.5 mg) and further purified by semi-preparative HPLC using a Synergi 4 μm Hydro-RP 80 Å column (10.00 × 250 mm) and isocratic elution gradient elution using 35% ACN / 65% H2O isocratic at the flow rate of 3.5 mL/min for 3 minutes the ramping up to 55% ACN in 22 minutes, then ramping up to 100% ACN in one minute and holding the gradient at 100% ACN for another 5 minutes yielding **1** (1.6 mg, RT = 23.3 min) and **2** (1.3 mg, RT =21.8 min) as a colorless, amorphous solid. The same HPLC conditions were used to isolate compounds **3** (rivulariapeptolide 1121, from subfraction 1-2-10 and 1-2-11, 1.1 mg, RT= 18.5min) and **4** (rivulariapeptolide 988, from subfraction 1-2-7, 1.3 mg, RT = 12.6 min). Fr. 1-1 was dissolved in 30% ACN/H2O and purified by preparative HPLC using a Kinetex 5 μm RP 100 Å column (21.00 × 150mm) and isocratic elution using 30% ACN/H2O for 10 minutes then ramping up to 50% in 10 minutes and then to 95% in 2 min at the flow rate of 20 mL/min, yielding 29 subfractions. Molassamide (compound **5**) was isolated from subfractions 1-1-6 and 2-1-10 were combined (3.3 mg) and further purified by semi-preparative HPLC using a Synergi 4 μm Hydro-RP 80 Å column (10.00 × 250 mm) and isocratic elution gradient elution using 35% ACN / 65% H2O isocratic at the flow rate of 3.5 mL/min for 3 minutes the ramping up to 55% ACN in 22 minutes, then ramping up to 100% ACN in one minute and holding the gradient at 100% ACN for another 5 minutes yielding **5** as a colorless, amorphous solid (1.8 mg) at RT = 10.8 min. The same HPLC conditions were used to isolate compound **6** (Molassamide B, from subfraction 1-1-4 and 1-1-5, 1.8 mg, RT = 13.7 min).

## Data Availability

All raw (.raw), deconvoluted (xtract.raw) and centroided (.mzXML or .mzML) mass spectrometry data as well as processed data feature table (.csv) and MS/MS spectra (.mgf) are available through the MassIVE repository (massive.ucsd.edu) with the identifier MSV000087964, MSV000088586 and MSV000088578. The MS/MS spectra of the new discovered derivatives, including tags as protease inhibitors, have been added to the GNPS library (gnps.ucsd.edu) with the following IDs: rivulariapeptolide 1185 (**1**): CCMSLIB00005723387; rivulariapeptolide 1155 (**2**): CCMSLIB00005723986, CCMSLIB00005720236; rivulariapeptolide 1121 (**3**): CCMSLIB00005723398; rivulariapeptolide 988 (**4**): CCMSLIB00005723393; molassamide (**5**): CCMSLIB00005723404; molassamide B (**6**): CCMSLIB00006710020. Raw NMR data for compounds **1**-**6** has been deposited to Zenodo (zenodo.org) and can be accessed under the following link: https://sandbox.zenodo.org/record/905199.

## Code Availability

The modified version of MZmine2.37 (corr.17.7) used in this study is available at https://github.com/robinschmid/mzmine2/releases. The code for the mass-offset matching for native metabolomics data analysis is available under https://github.com/Functional-Metabolomics-Lab/Native-Metabolomics.

## Acknowledgment

We thank the Deutsche Forschungsgemeinschaft for the support of D.P. through a postdoctoral research fellowship (PE 2600/1-1) and of D.P. and C.C.H. through the CMFI Cluster of Excellence (EXC 2124). R.R., P.C.D., and W.H.G. were supported by the Gordon and Betty Moore Foundation (GBMF7622) and by the US National Institutes of Health (R01 GM107550, P41 GM103484 and R03 CA211211). WB was supported in part by the Research Foundation – Flanders (12W0418N).

## Contributions

R.R., W.H.G. and D.P. conceived the study. R.R., K.L.A., C.B.N., E.J.C., and W.H.G. collected and extracted environmental samples. A.T.A., and D.P. developed the native metabolomics approach. W.B. wrote software code. M.W. aided in integration with GNPS tags. R.R., A.T.A., P.S., and D.P. performed MS experiments. R.R. and M.L.M. performed compound isolation. R.R. carried out NMR experiments. R.R., P.S., C.C.H., and D.P. performed total hydrolysis and derivatization experiments. P.F., C.L., and A.J.O. performed activity assays. I.B.S. performed docking studies. R.R., and D.P. wrote the manuscript. All authors edited and approved the final manuscript.

## Ethics declarations

### Competing interests

P.C.D. and W.H.G. are scientific advisors of Sirenas. P.C.D. is a scientific advisor of Galileo, Cybele, and scientific advisor and co-founder of Ometa Labs LLC and Enveda with approval by the UC San Diego. M.W. is a founder of Ometa Labs LLC.

## Supplementary information

**Supplementary Results, Methods, Figures, and NMR tables.**

## Notes

https://massive.ucsd.edu/ProteoSAFe/dataset.jsp?task=7ab346a09ca64aa1bd19bdc035801c15

https://sandbox.zenodo.org/record/905199

### Summary of Updates

Revised Manuscript and SI

