## Supplemental Information for "Native Metabolomics Identifies the Rivulariapeptolide Family of Protease Inhibitors"

### TABLE OF CONTENTS


| Content | Page |
|---|---|
| **Supplementary Methods** | 3 |
| **Supplementary figures** | |
| Figure S1: Optimization of native MS conditions | 8 |
| Figure S2: Selectivity tests and comparison of native metabolomics with direct infusion | 9 |
| Figure S3: Correlation Networking of metabolite-protein binding | 10 |
| Figure S4: SMART 2.1 reproducibility test and mixture analysis | 11 |
| Figure S5: Ion identity network and CANOPUS analysis of *Rivularia* sp fractions | 12 |
| Figure S6: $^{1}$H NMR of compound **1** in DMSO-*d*6 | 13 |
| Figure S7: $^{1}$H-$^{1}$H COSY-CLIP of compound **1** in DMSO-*d*6 | 13 |
| Figure S8: $^{1}$H-$^{13}$H HSQC of compound **1** in DMSO-*d*6 | 14 |
| Figure S9: $^{1}$H-$^{13}$C HMBC of compound **1** in DMSO-*d*6 | 14 |
| Figure S10: $^{1}$H-$^{13}$C HSQC-TOCSY of compound **1** in DMSO-*d*6 | 15 |
| Figure S11: $^{1}$H-$^{1}$H NOESY of compound **1** in DMSO-*d*6 | 15 |
| Figure S12: $^{1}$H NMR of compound **2** in DMSO-*d*6 | 16 |
| Figure S13: $^{13}$C NMR of compound **2** in DMSO-*d*6 | 16 |
| Figure S14: $^{1}$H-$^{1}$H COSY of compound **2** in DMSO-*d*6 | 17 |
| Figure S15: $^{1}$H-$^{13}$C HSQC of compound **2** in DMSO-*d*6 | 17 |
| Figure S16: $^{1}$H-$^{13}$C HMBC of compound **2** in DMSO-*d*6 | 18 |
| Figure S17: $^{1}$H-$^{13}$C TOCSY of compound **2** in DMSO-*d*6 | 18 |
| Figure S18: $^{1}$H-$^{1}$H NOESY of compound **2** in DMSO-*d*6 | 19 |
| Figure S19: $^{1}$H NMR of compound **3** in DMSO-*d*6 | 19 |
| Figure S20: $^{1}$H-$^{1}$H COSY of compound **3** in DMSO-*d*6 | 20 |
| Figure S21: $^{1}$H-$^{13}$C HSQC of compound **3** in DMSO-*d*6 | 20 |
| Figure S22: $^{1}$H-$^{13}$C HMBC of compound **3** in DMSO-*d*6 | 21 |
| Figure S23: $^{1}$H-$^{13}$C TOCSY of compound **3** in DMSO-*d*6 | 21 |
| Figure S24: $^{1}$H-$^{1}$H NOESY of compound **3** in DMSO-*d*6 | 22 |
| Figure S25: $^{1}$H NMR of compound **4** in DMSO-*d*6 | 22 |
| Figure S26: $^{1}$H-$^{13}$C HSQC of compound **4** in DMSO-*d*6 | 23 |
| Figure S27: $^{1}$H-$^{13}$C HMBC of compound **4** in DMSO-*d*6 | 23 |
| Figure S28:$^{1}$H-$^{13}$C HSQC-TOCSY of compound **4** in DMSO-*d*6 | 24 |
| Figure S29: $^{1}$H-$^{1}$H ROESY of compound **4** in DMSO-*d*6 | 24 |
| Figure S30: $^{1}$H NMR spectrum of compound **5** in DMSO-*d*6 | 25 |
| Figure S31: $^{1}$H-$^{1}$H COSY of compound **5** in DMSO-*d*6 | 25 |
| Figure S32: $^{1}$H-$^{13}$C HSQC of compound **5** in DMSO-*d*6 | 26 |
| Figure S33: $^{1}$H-$^{13}$C HMBC of compound **5** in DMSO-*d*6 | 26 |
| Figure S34: $^{1}$H NMR spectrum of compound **6** in DMSO-*d*6 | 27 |
| Figure S35: $^{1}$H-$^{1}$H COSY of compound **6** in DMSO-*d*6 | 27 |
| Figure S36: $^{1}$H-$^{13}$C HSQC of compound **6** in DMSO-*d*6 | 28 |
| Figure S37: $^{1}$H-$^{13}$C HMBC of compound **6** in DMSO-*d*6 | 28 |
| Figure S38: $^{1}$H-$^{13}$C HSQC-TOCSY of compound **6** in DMSO-*d*6 | 29 |

Figure S39: Mirror MS² plot of molassamide from this study (black) and GNPS library spectrum (green) 29
Figure S40: Comparison of 1H spectra of molassamide from this study and molassamide isolated by others 30
Figure S41: MS/MS annotations for compounds **1**, **2**, **3** 31
Figure S42: Absolute and relative stereochemistry determination for compound **2** 32
Figure S43: Docking and structure activity relationship studies of compound **1**-**6** 33

**Supplementary Tables**

Table S1. NMR table for rivulariapeptolide 1185 (**1**) 34
Table S2: NMR table for rivulariapeptolides 1155 (**2**), 1121 (**3**), and 988 (**4**) 35
Table S3: NMR table for molassamide (**5**) and molassamide B (***6***) 37
Table S4: Top 50 chymotrypsin inhibitors among Ahp-cyclodepsipeptides 39

**Supplementary References** 41

#### Supplementary Methods

##### Reproducibility of SMART results

Mixture analysis of AHP-cyclodepsipeptide-containing SPE-fraction by acquiring data on two different NMR spectrometers and subsequent SMART 2.1 analysis[1]. The same crude extract was splitted into two equal halves and processed in the same way, but on different days via solid phase extraction into 4 fractions (1-1, 1-2, 1-3, 1-4 & 2-1, 2-2, 2-3, 2-4, respectively). HSQC NMR data for Fraction 1-1 (1 mg) was acquired in a 1.7 mm TCI MicroCryoProbe (599.10 MHz) to obtain the correlations of the major components of that fraction in about 13 minutes by applying non uniform sampling[2] and Acceleration by Sharing Adjacent Polarization[3] protocols. HSQC NMR data for Fraction 2-1 (1 mg) was acquired on a JEOL ECZ 500 NMR spectrometer equipped with a 3 mm inverse detection probe to obtain the correlations of the major components of that fraction in about 14 hours. The raw data was processed (phase correction, baseline correction, removal of t1 noise) with MestreNova Version 12.0 and the major correlations were manually peak picked. The resulting table was exported and analyzed by SMART 2.1 (https://smart.ucsd.edu/classic).

#### In silico MS/MS annotation

Comparison of the SMART results revealed that eight of ten predicted structures were identical and five or respectively six out of ten compounds were AHP-cyclodepsipeptides. In depth analysis of the top ten predicted structures with NPClassifier[4] (https://npclassifier.ucsd.edu/) showed that all, but one structure were classified as cyclic depsipeptides and all of the structures were further classified as polyketides, suggesting a NRPS/PKS hybridic pathway for the biosynthesis of the main compound in fraction 1-1 (and 2-1, respectively).

We annotated the Ahp-cyclodepsipeptide candidates as members of the subfamily rivulariapeptolides followed by their molecular weight if CANOPUS[5] detected the substructure element 'delta-lactam' or 'piperidone' with a posterior probability of > 99% in combination with the detection of substructural elements such as 'pyrrolidinecarboxamide', '*N*-acylpyrrolidines', and 'proline and derivatives' with a posterior probability of > 90%, indicative of the characteristic *N*-butyrylated proline residue. All MS/MS of isolated compounds were deposited as public spectra within GNPS[6] along with confirmed activity of the respective compounds as protease inhibitors utilizing GNPS tags.

#### Relative and absolute configurations of the rivulariapeptolides

The absolute configurations of the amino acids were determined by UHPLC-MS analysis of the acid hydrolysates of 2 (rivulariapeptolide 1155) and its PDC oxidation product (**Figure S41a**). Oxidation followed by acid hydrolysis liberated glutamic acid, allowing the assignment of the C-3 position of the Ahp (3-amino-6-hydroxy piperidone) unit. The analysis revealed L-configuration for C-3 as well as for the C-alphas of every further amino acid in the rivulariapeptolides (as it is the case for all previously reported cyanobacterial Ahp-cyclodepsipeptides). The relative stereochemistry of the Ahp moiety was determined to be (3S*, 6R*)-Ahp based on NMR spectroscopic data (**Figure S41b**). In the Ahp ring system, NOESY correlations were observed between the diaxially oriented H-3 (δ 3.78 ppm), H-5a (δ 1.71 ppm), and the equatorially oriented H-6 (δ 5.07 ppm, br s

($J < 1$ Hz))[7]. Thus, the hydroxyl group of C-6 had to be axially oriented and this axial orientation of the 6-OH group is responsible for the downfield shift of H-4a (δ 2.42 ppm). Furthermore, the NOESY correlations between H-3 (δ 3.78 ppm) and the equatorial H-4b (δ 1.58 ppm) supported the assigned relative stereochemistry. Together with the results from the PDC oxidation and Marfey's analysis we determined the absolute configuration to be (3S, 6R)-Ahp in line with the stereochemical assignments in other Ahp-cyclodepsipeptides such as tutuilamide A and molassamide, respectively[8,9]. The geometry of the Abu olefinic bond was determined as "Z" based on HMBC and NOE correlations in DMSO-*d6*. A four-bond HMBC correlation between $H_3$-4 and C-1 of Abu indicates a "w" configuration for bonds between these atoms and, therefore, the Z-geometry for the double bond[10]. NOESY correlations from $H_3$-4 of Abu (δ 1.51) to H-2 of Thr-1 (δ 4.59) and $H_3$-4 of Thr-2 (δ 1.11) further support this configuration.

**Pyridinium dichromate (PDC) oxidation**

Rivulariapeptolide 1155 (Compound 2, 0.5 mg) was dissolved in $CH_2Cl_2$ (0.5 mL) and mixed with PDC (2.0mg). After stirring at rt for 5h, the reaction was quenched by shaking with 3 x 1 mL portions of $H_2O$, and the the $CH_2Cl_2$ phase was dried under $N_2$. The resulting oxidized material was analyzed by advanced Marfey's method and subsequent UHPLC-MS as described below.

**Advanced Marfey's method**

Rivulariapeptolide 1155 (Compound 2, 0.3 mg) and the residue of the PDC reaction were hydrolyzed with 6N HCl (0.3 mL) and heated at 110°C for 16 hrs in sealed tubes. The hydrolysates were concentrated to dryness under $N_2$, and each treated with a solution of 1-fluoro-2-4-dinitrophenyl-5-L-alanine amide (FDAA, 1 mg/mL) in acetone (100 µL) and a solution of 1 M $NaHCO_3$ (0.3 mL) in a sealed vial at 90 °C for 5 min. The reaction mixture was neutralized with 1N HCl (300 µL) and diluted with $CH_3CN$ (0.7 mL). The resulting solution was analyzed by LC-MS via a Vanquish UHPLC Sytsem couple to A Q-Exactive HF Mass Spectrometer (Thermo-Fisher). RP-UHPLC was performed on a Kinetex C18 column with 150 x 2 mm 1.8 um particle size, 100 A pore size. The gradient elution profile started at 5% Acetonitrile (ACN) /95% $H_2O$ (acidified with 0.1% formic acid (FA)) and was ramped to 30% ACN at 8 min following an increase to 99% ACN at 10 min. The column was the washed for three minutes at 99% ACN, and then re-equilibrated over 3 min at 5% CAN. The flow was set to 0.5 mL/min. Electrospray ionization (ESI) parameters were set to 50 arbitrary units (AU) sheath gas flow, auxiliary gas flow was set to 12 AU and sweep gas flow was set to 1 AU. Auxiliary gas temperature was set to 400 °C. The spray voltage was set to 3.5 kV and the inlet capillary was heated to 250 °C. S-lens level was set to 50 V applied.  MS scan range was set to 200-2000 *m/z* with a resolution at *m/z* 200 ($R_{m/z\ 200}$) of 120,000 with one micro-scan. The maximum ion injection time was set to 50 ms with automatic gain control (AGC) target of 3E6. Raw files where converted to .mzML format using MSconvert from the proteowizard software package. Extracted ion chromatograms (XICs) of the expected amino acid-FDAA conjugates where generated with the GNPS Dashboard with 5 ppm mass tolerance. Retention times and XICs of the amino acid-FDAA conjugates from the reaction mixtures were compared with the commercially available amino acid standards that were derivatized using identical methodology and allowed us to establish **2**.

**Docking and structure activity relationship studies reveal potential for therapeutic optimization (figure S35)**

Next, we performed docking studies to rationalize the bioactivity of the isolated compounds and to explore the potential for structure modifications that could yield lead structures for therapeutic interventions. The isolated compounds were docked by induced-fit, inside the binding pocket of alpha-chymotrypsin (PDBID 4Q2K) and all were found to have a similar binding mode (figure S35b, d, e). Crystal structures of Ahp-cyclodepsipeptides in complex with serine proteases by others[8,11] indicate that inhibition is based on a substrate-like binding mode in which distinct amino acid residues occupy the S- and S'-pockets, however, proteolytic cleavage does not occur, because the AHP moiety occupies an important part of the active site pocket. Furthermore, it has been demonstrated that the residue immediately following the Ahp unit (position 5, see Scheme S1) should bind in the S1 specificity-determining pocket[12,13]. Structure-activity relationships towards elastase inhibition revealed that the 2-amino-2-butenoic acid (Abu) moiety incorporated within the macrocycle, contributes to potent elastase inhibition[8,14].

This is in accordance with our experimental results where the replacement of the Abu (compound **2**) moiety for Leu (**1**) led to a fivefold decrease in potency towards elastase, but simultaneously to a more than threefold increase in potency towards chymotrypsin. The previously reported importance of a polar functionality in the side chain towards more potent elastase inhibition could not be corroborated by our data[11,14] .Masking, the secondary alcohol of the threonine in the side chain by an esterification with *N*-butyryl proline (**2**) instead of bearing the free polar hydroxyl group (**4**) increased the potency towards elastase. Binding to proteinase K, however, seems to be more favorable with less rigid and bulky side chain substituents as found in compounds **5** and **6**.

**Structural analysis**

We modeled the rivulariapeptolides peptides bound to α-chymotrypsin, based on the crystal structure (PDB ID 4GVU) of the macrocyclic peptide lyngbyastatin 7, which shares a chemical scaffold with the rivulariapeptolides, bound to elastase, which has 37% sequence identity with alpha-chymotrypsin and a similar three-dimensional structure. A crystal structure of alpha-chymotrypsin (PDB ID 4Q2K) was aligned with the elastase structure, using the Molecular Operating Environment (MOE, version 2019.01) software, transferred the lyngbyastatin 7 molecule to the chymotrypsin structure, and energy-minimized the resulting bimolecular complex. We then used ChemDraw integration with MOE to sketch each of the rivulariapeptidolides and energy-minimized each resulting complex. Minimization was performed using the Amber10:EHT force-field, no constraints were applied, and the convergence criterion was set to RMS of 0.1 kcal $mol^{-1}$ $Å^2$.

#### Supplementary Figures

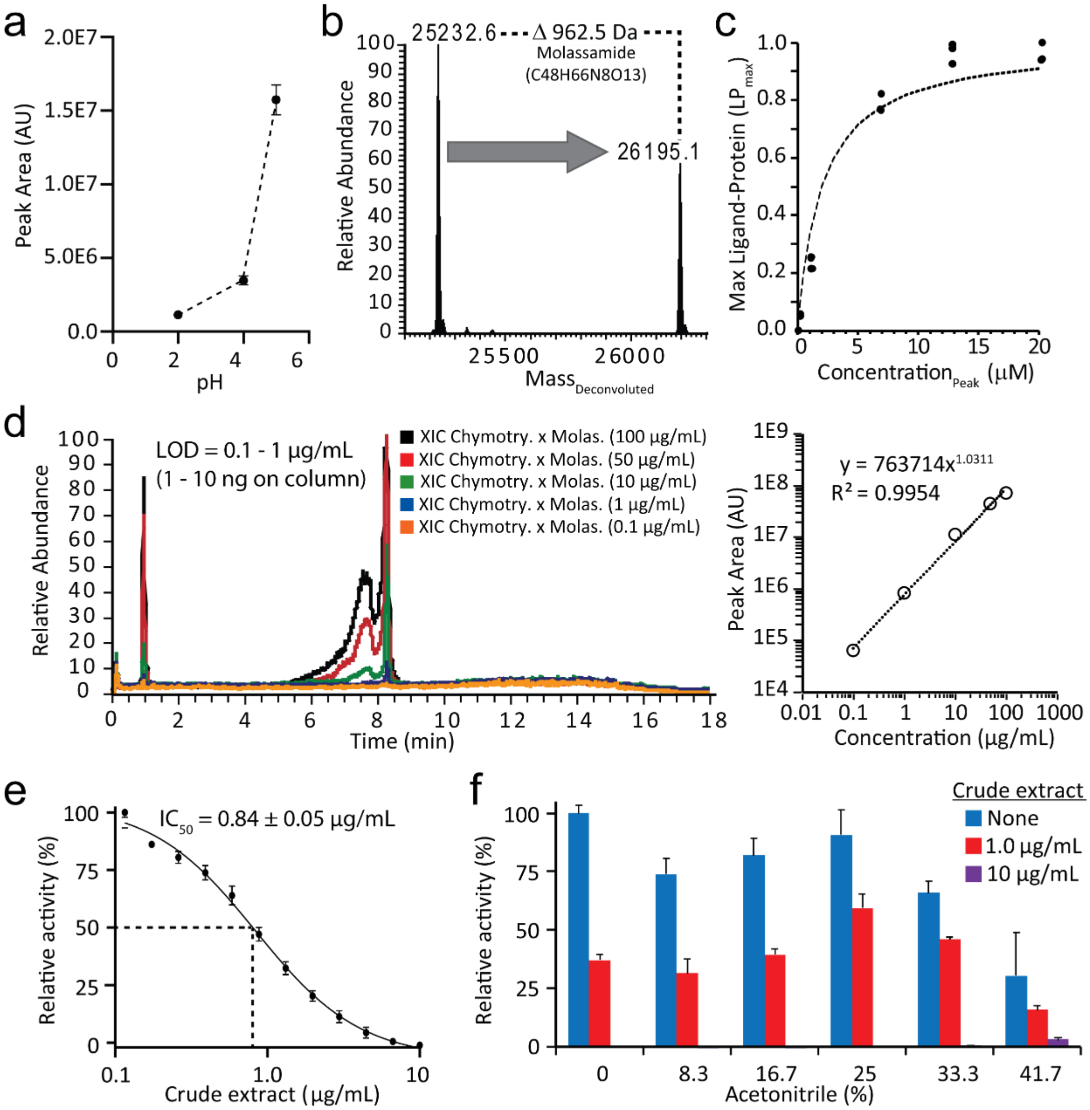


**Figure S1:** Optimization of native MS conditions a) Integrated peak area of the protein-ligand complex increases in a pH-dependent manner with increasing pH. (b) Molassamide was screened against chymotrypsin as a proof-of-concept experiment. A Δ*m/z* of 962.5 Da, the difference between unbound chymotrypsin and the chymotrypsin-molassamide complex, corresponds to the *m/z* of molassamide. (c) Concentration-dependent increase in chymotrypsin-molassamide complex, measured by native metabolomics. $Concentration_{Peak}$ (µM) refers to concentration of ligand. (d)The limit of detection (LOD) for the molassamide-chymotrypsin interaction was determined by generating a calibration curve for molassamide and injecting molassamide stock solutions of 0.1 – 100 µg/mL and chymotrypsin. The extracted ion chromatograms (XICs) for the chymotrypsin-molassamide complex are shown. (e) Concentration-dependent inhibition of chymotrypsin activity by 10 µg/mL of cyanobacterial crude extract in 10 mM ammonium acetate pH 4.5. (f) Relative chymotrypsin activity of cyanobacterial crude extract (1 µg/mL and 10 µg/mL) in native MS conditions (10 mM ammonium acetate pH 4.5 plus increasing concentrations of acetonitrile) compared with no addition of crude extract.

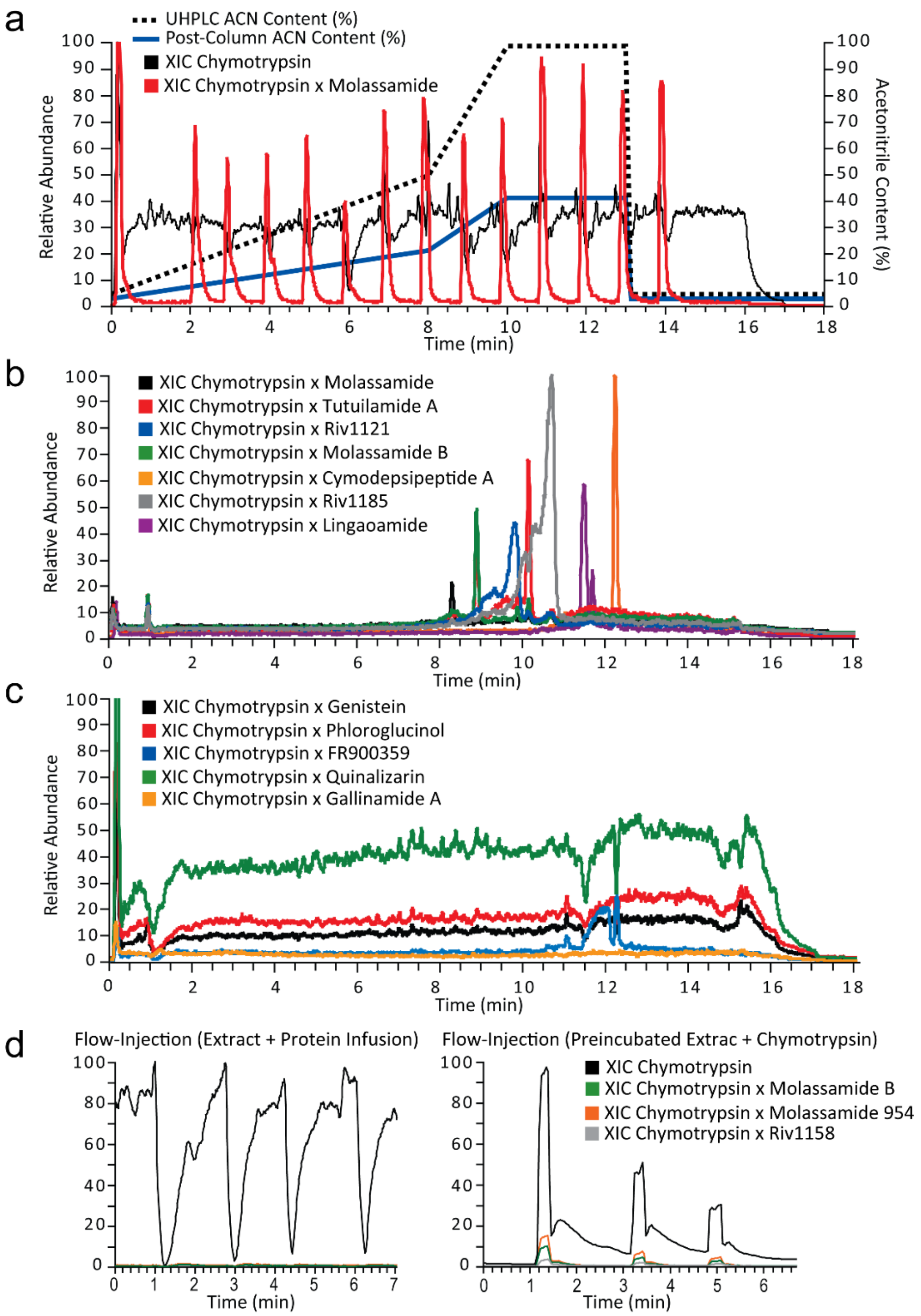


**Figure S2**: Selectivity tests and comparison of native metabolomics with direct infusion. (a) To test the variability of the changing acetonitrile concentration during the LC separation, we removed the UHPLC column and performed a series of flow injections over the full gradient. The XIC of the molassamide bound chymotrypsin reveals similar signal responses throughout the gradient (5-99% ACN on column) (b) Native MS run of positive controls versus alpha-chymotrypsin. Binding of all positive controls to chymotrypsin can be detected in the expected mass range by extracted ion chromatograms (XICs). (c) Native MS run of negative controls genistein, phloroglucinol, FR900359, quinalizarin, gallinamide A versus alpha-chymotrypsin. The drop in detected protein mass is clearly visible due to the co-elution of the small molecules, but no binding of the negative controls to chymotrypsin can be detected in the expected mass range.

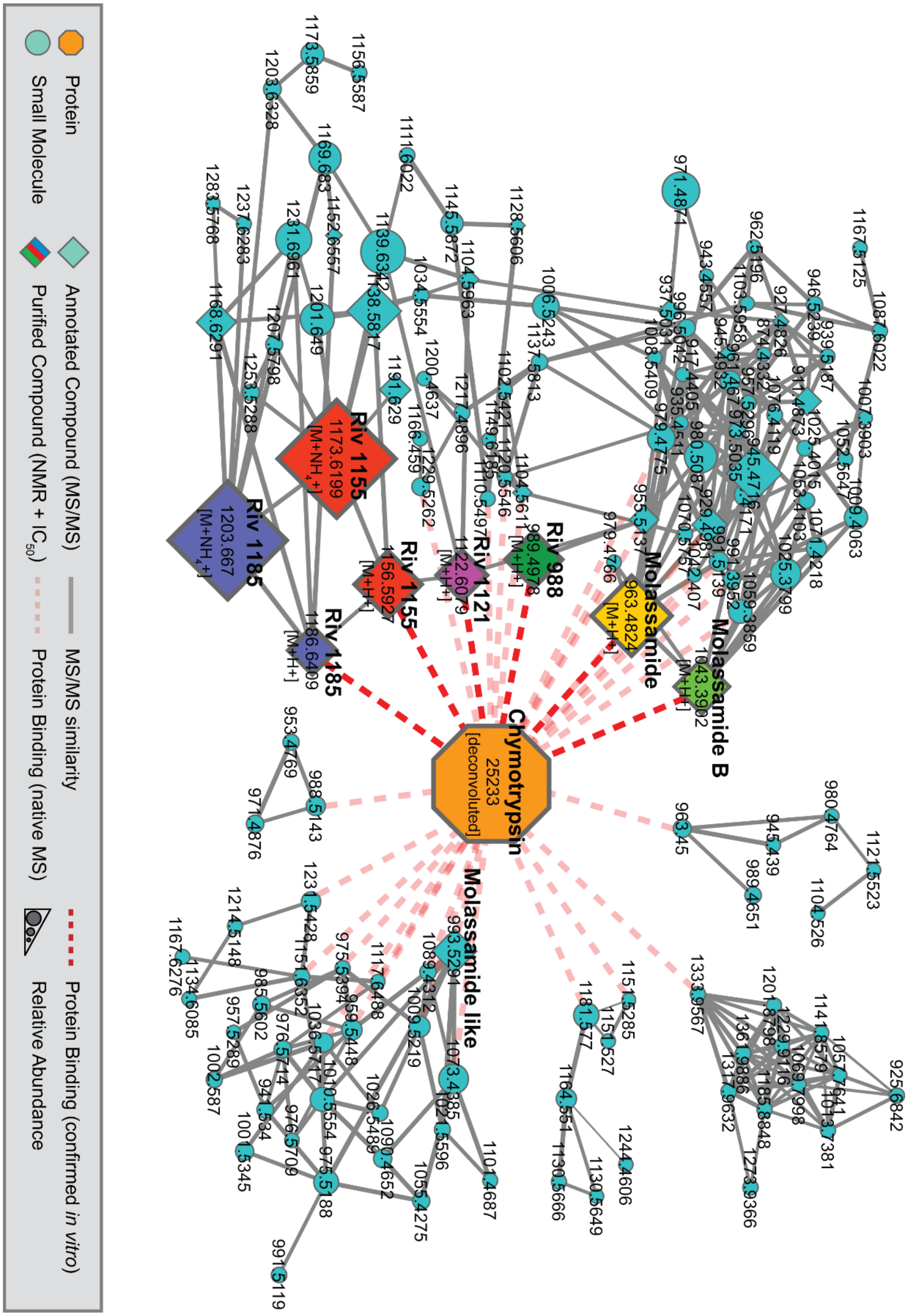


**Figure S3**: Correlation molecular network of chymotrypsin and putative small molecule inhibitors binders by native MS.

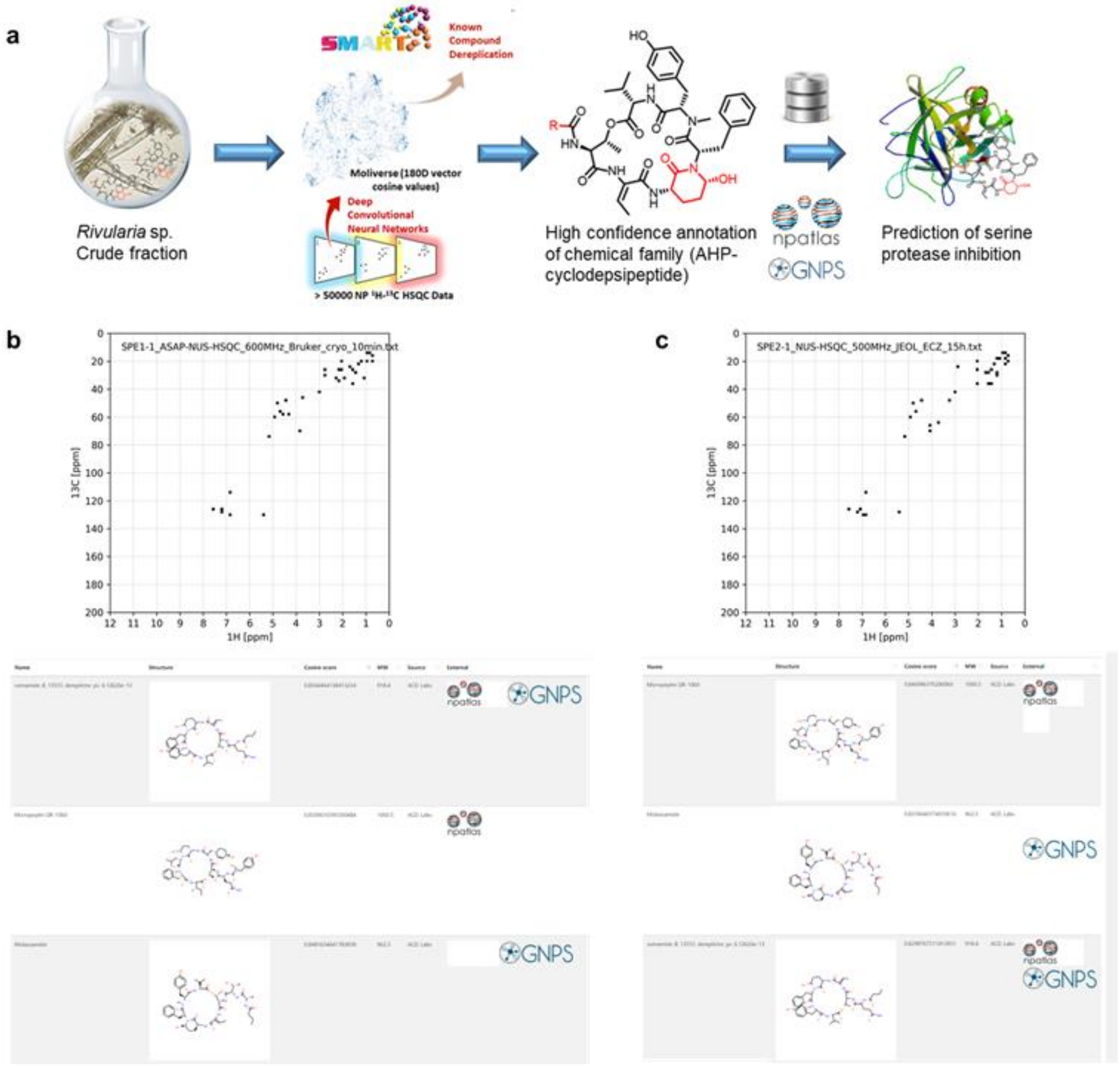


**Figure S4**: SMART 2.1 reproducibility test and mixture analysis (a) Workflow for SMART mixture analysis and bioactivity prediction. The crude extract was divided into two identical halves. One half was separated by solid phase extraction into four fractions of decreasing polarity (1-1, 1-2, 1-3, 1-4). The second half of the crude extract was fractionated in an identical fashion as described for fraction 1-1, yielding the equivalent SPE fraction 2-1. (b) Top 3 SMART 2.1 predicted compounds for SPE fraction 1-1. NMR acquisition was performed on 1 mg material in a Bruker 1.7 mm TCI MicroCryoProbe (599.10 MHz) to obtain the correlations of the major components of that fraction in 13 minutes. (c) Top 3 SMART 2.1 predicted compounds from SPE fraction 1-1. NMR acquisition was performed on 1 mg of fraction 2-1 on a JEOL 500 MHz ECZ instrument in 15 hours. For complete results refer to https://smart.ucsd.edu/resultclassic?task=4bf3d8c3-3a96-419e-a5f5-416e30acc56b, (fraction 1-1, Bruker 1.7 mm TCI MicroCryoProbe (599.10 MHz) or https://smart.ucsd.edu/resultclassic?task=a44d9275-dc58-49dd-9320-5c620fb89d3e (fraction 2-1, JEOL 500MHz ECZ instrument), respectively.

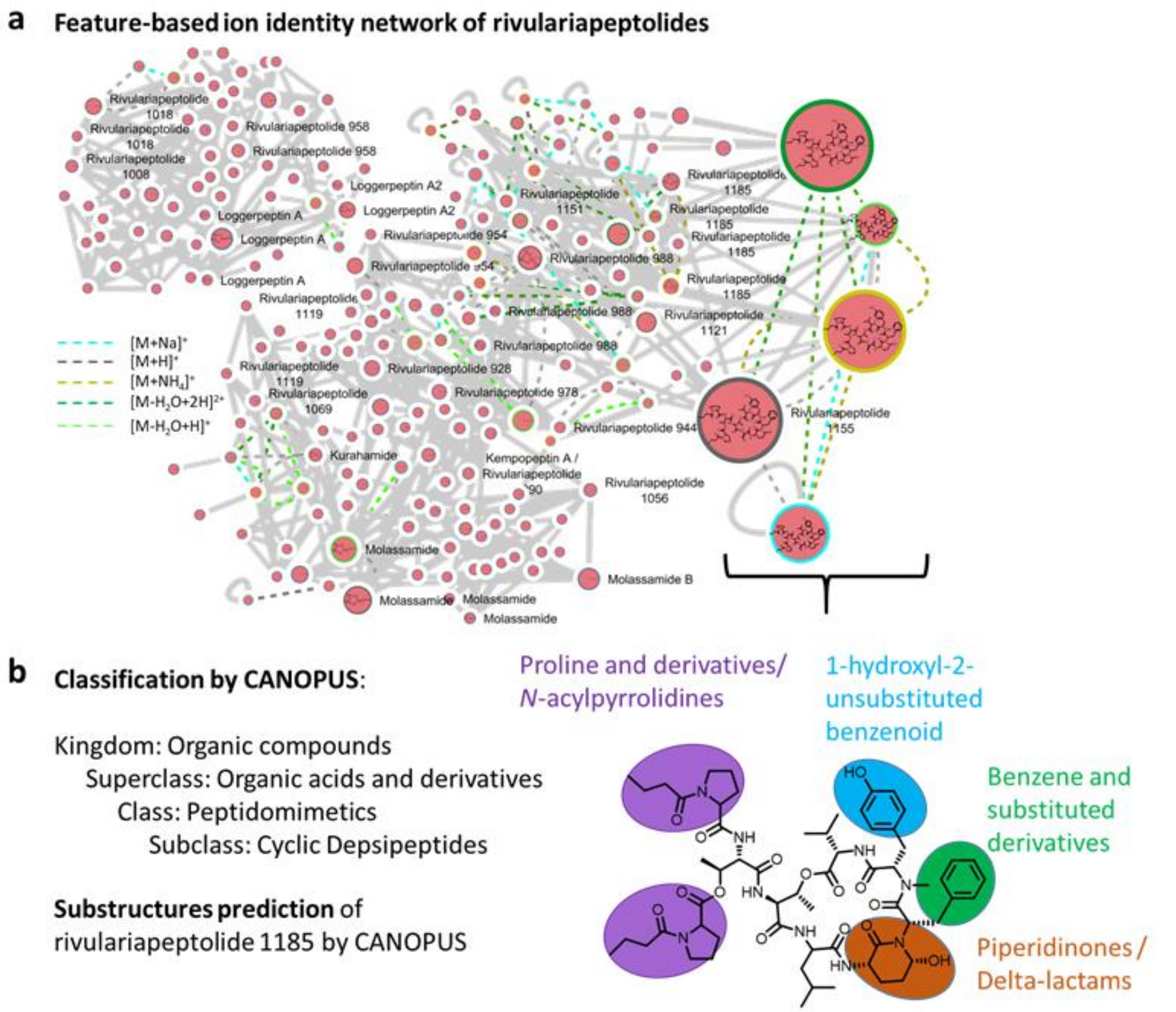


**Figure S5**: (a) Feature-based ion identity networking and (b) classification, substructure analysis of rivulariapeptolides and other Ahp-cyclodepsipeptides by CANOPUS.

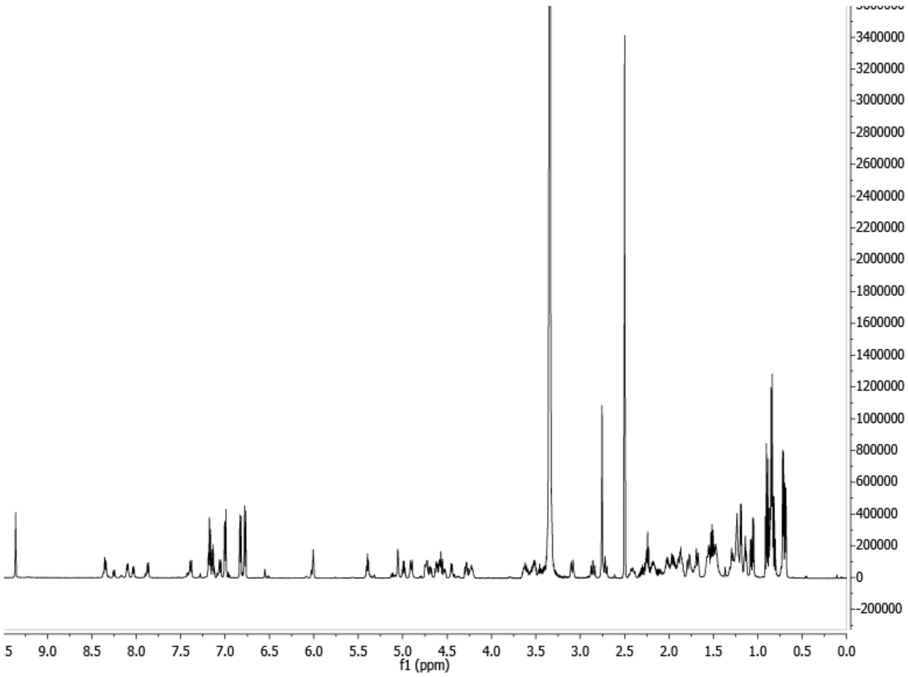


**Figure S6:** $^{1}$H NMR spectrum of rivulariapeptolide 1185 in DMSO-*d6*, 600 MHz.

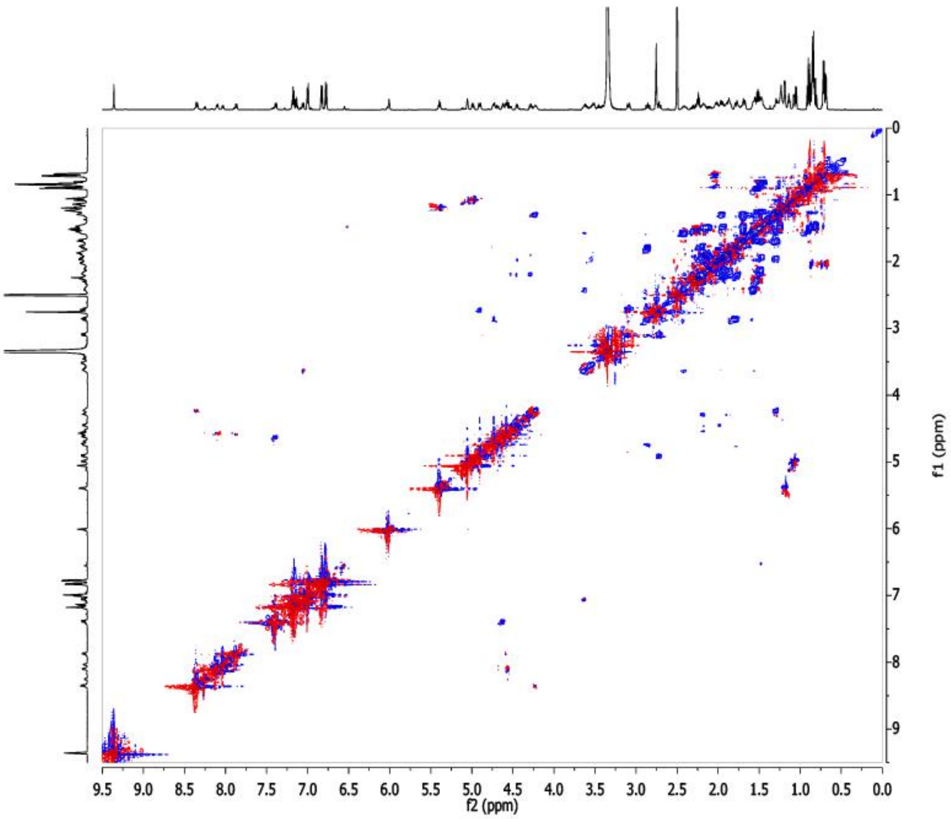


**Figure S7:** $^{1}$H-$^{1}$H COSY_CLIP spectrum of rivulariapeptolide 1185 in DMSO-*d6*, 600 MHz.

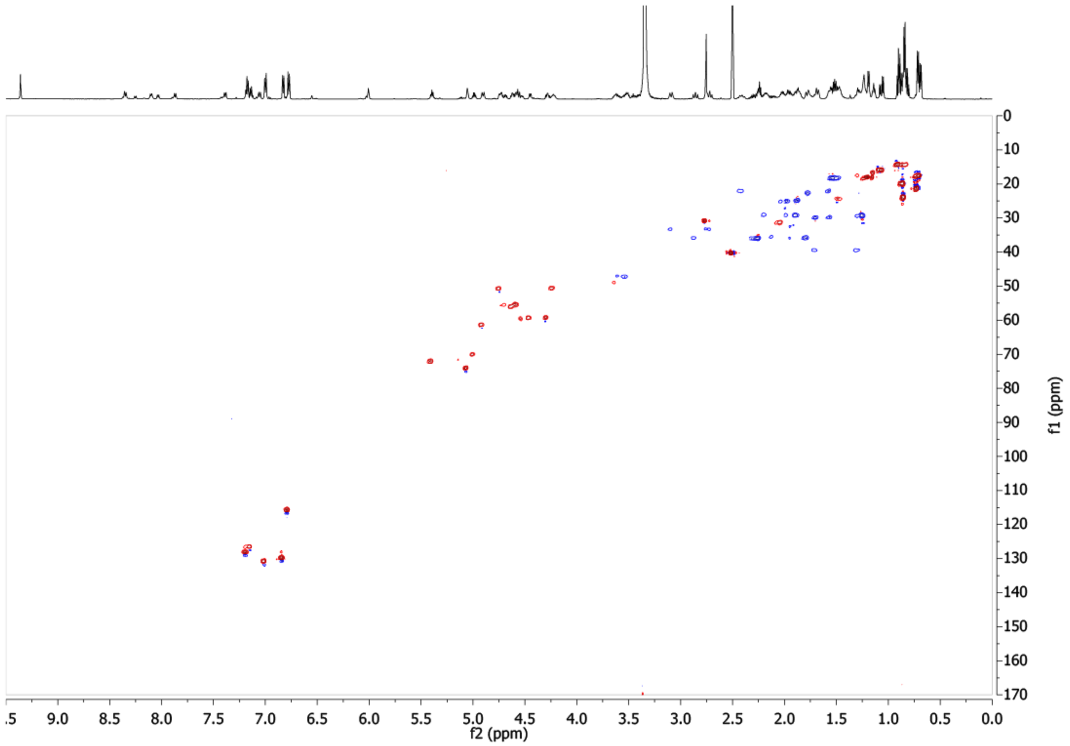


**Figure S8:** $^{1}$H-$^{13}$C HSQC spectrum of rivulariapeptolide 1185 in DMSO-*d6*, 600 MHz.

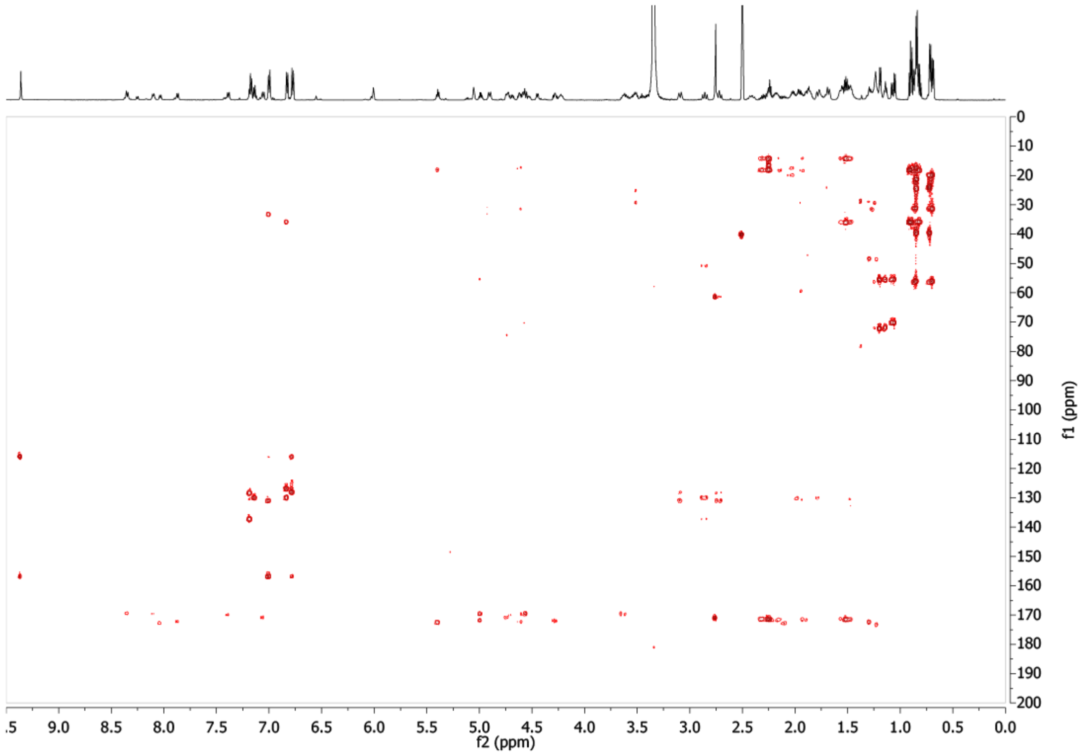


**Figure S9:** $^{1}$H-$^{13}$C HMBC spectrum of rivulariapeptolide 1185 in DMSO-*d6*, 600 MHz.

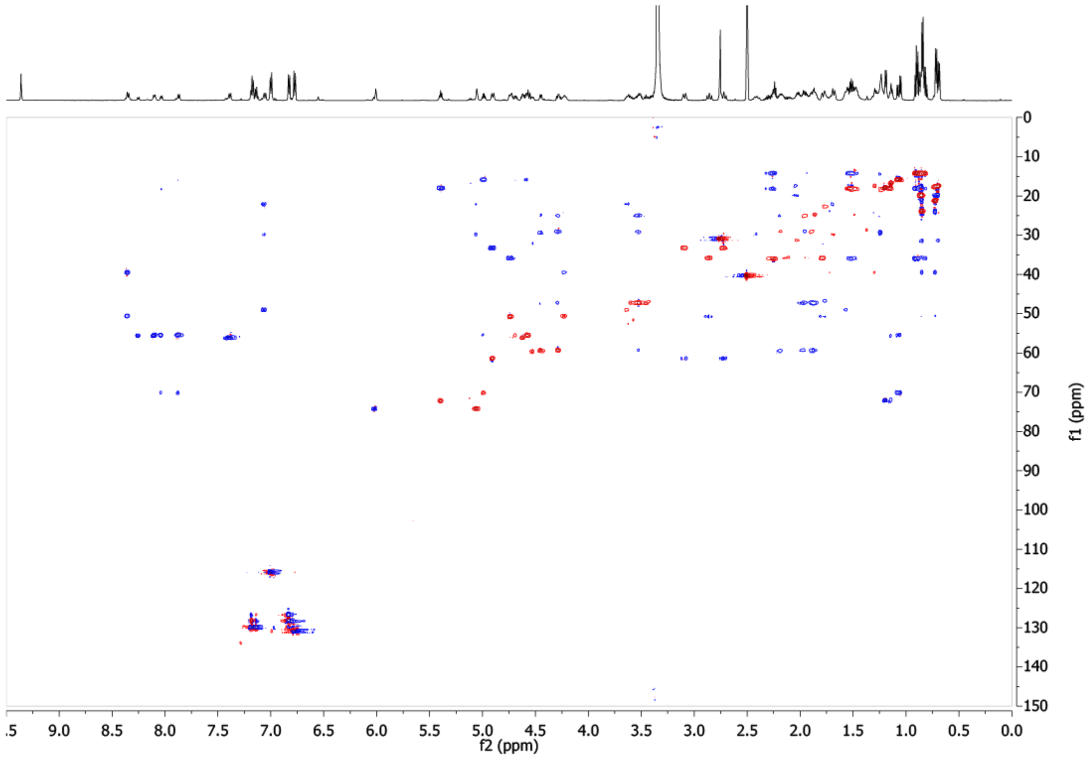


**Figure S10:** $^{1}H$-$^{13}C$ HSQC-TOCSY spectrum of rivulariapeptolide 1185 in DMSO-*d6*, 600 MHz.

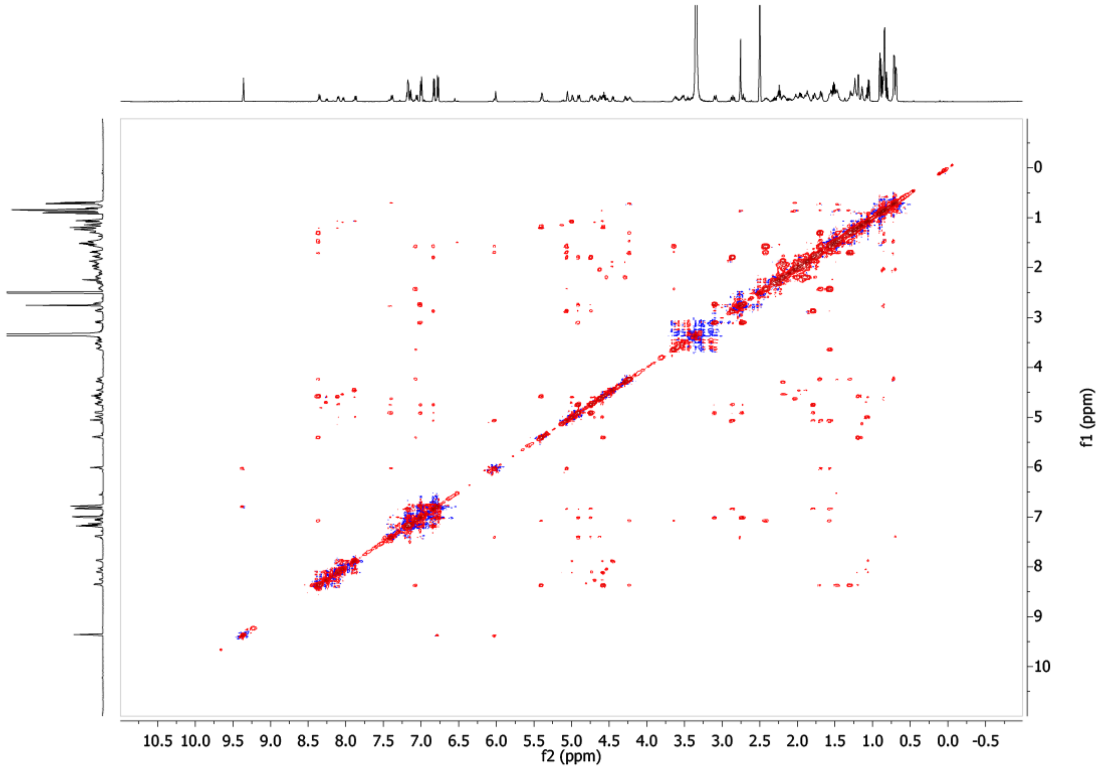


**Figure S11:** $^{1}H$-$^{1}H$ NOESY spectrum of rivulariapeptolide 1185 in DMSO-*d6*, 600 MHz.

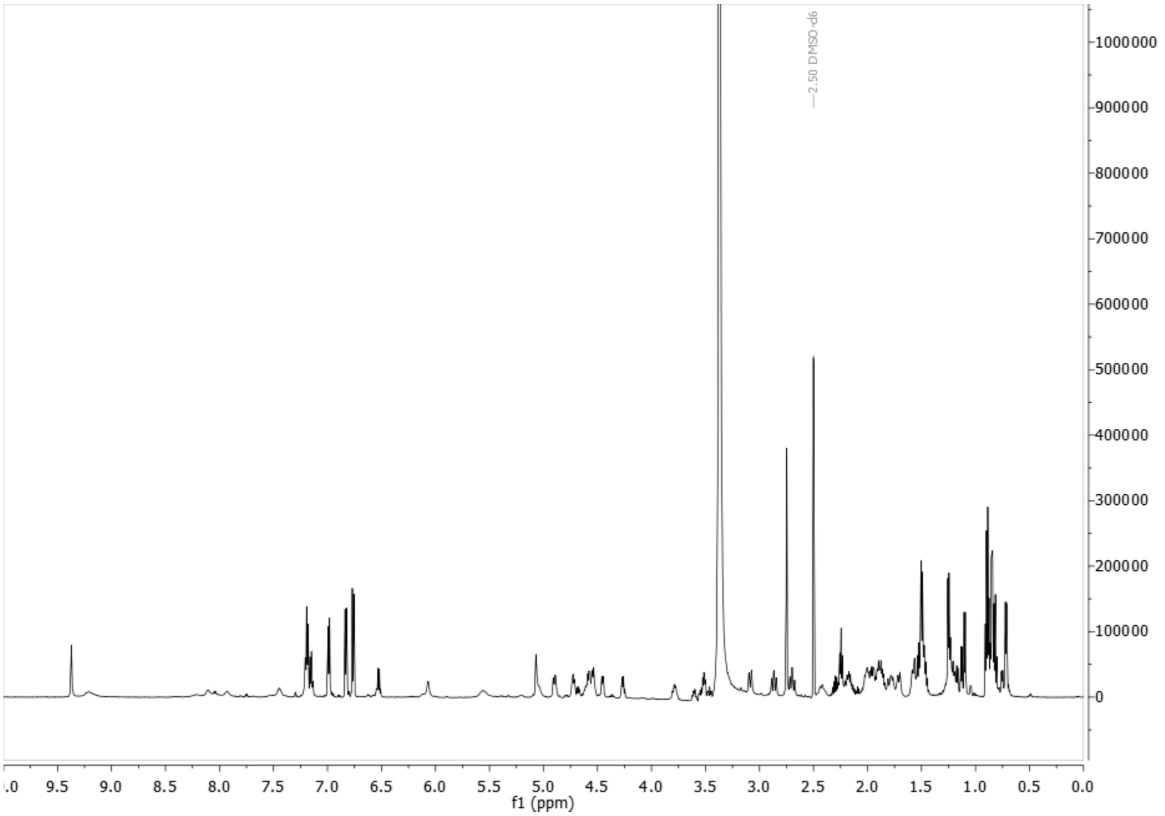


**Figure S12:** $^{1}$H NMR spectrum of rivulariapeptolide 1155 in DMSO-*d6*, 600 MHz.

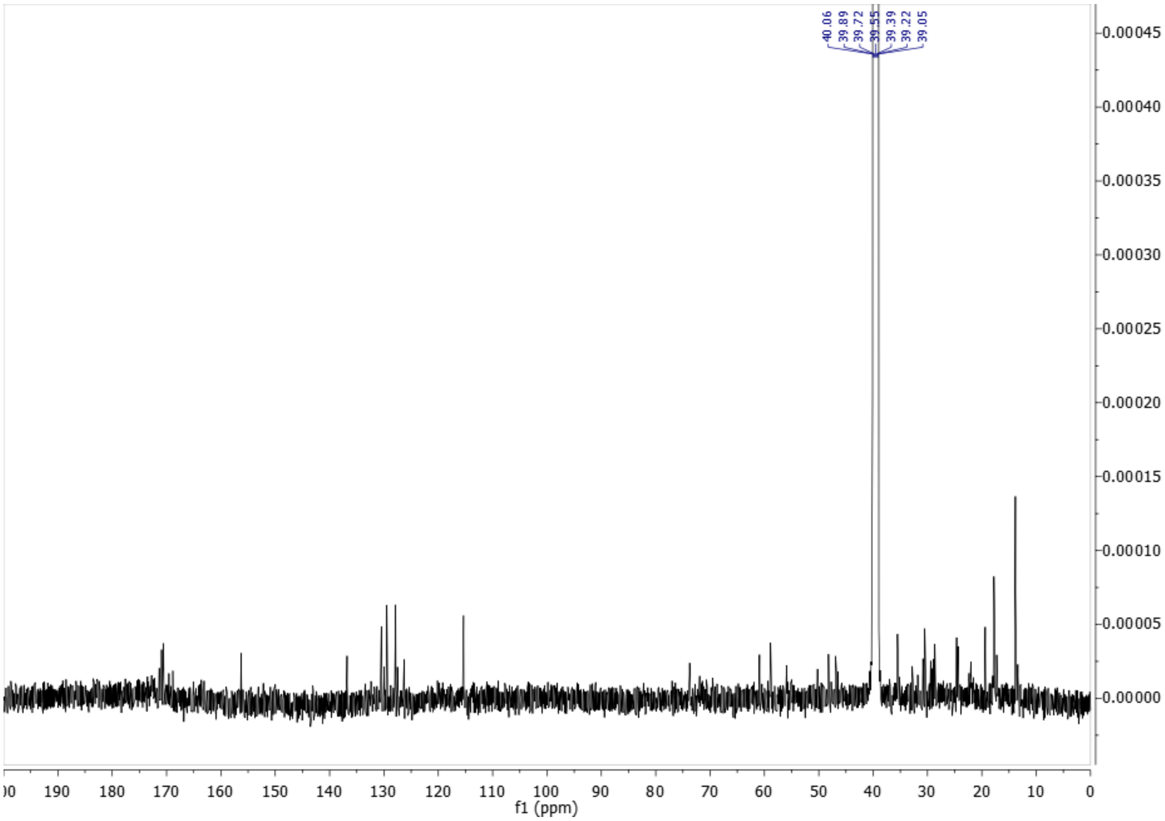


**Figure S13:** $^{13}$C NMR spectrum of rivulariapeptolide 1155 in DMSO-*d6*, 125 MHz.

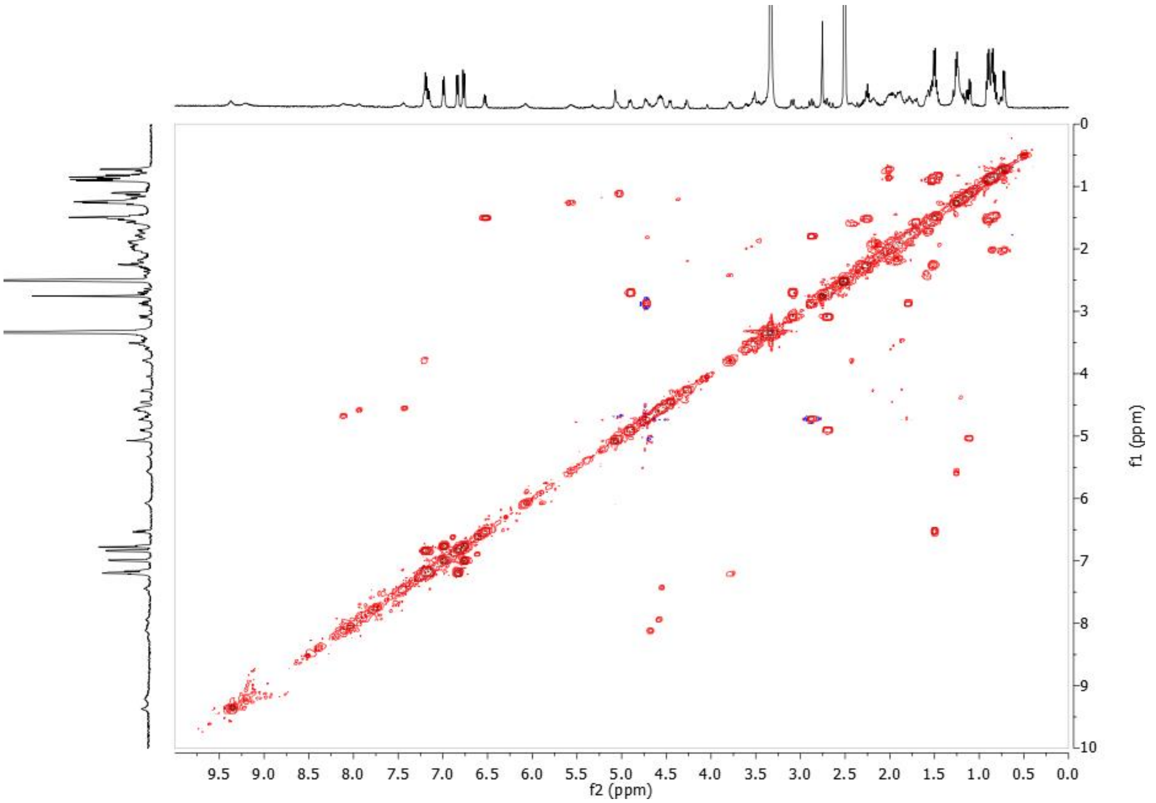

**Figure S14:** $^{1}H$-$^{1}H$ COSY spectrum of rivulariapeptolide 1155 in DMSO-*d6*, 500 MHz.

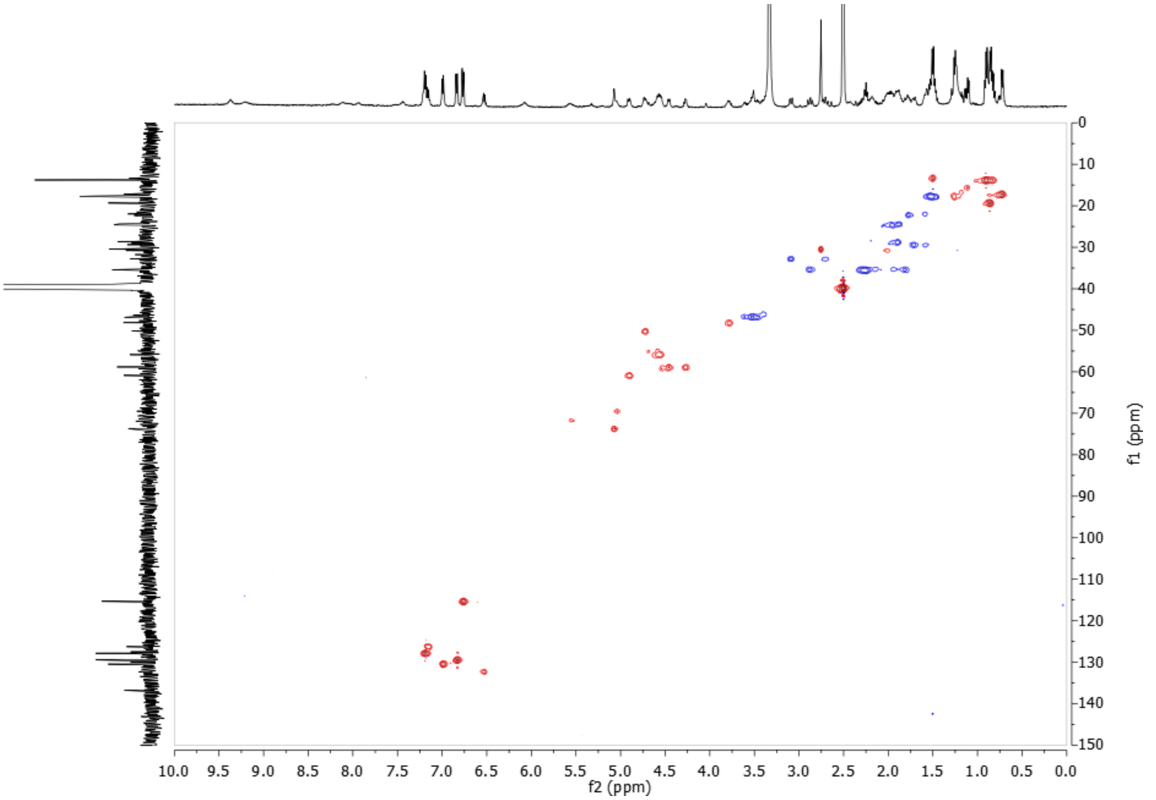

**Figure S15:** $^{1}H$-$^{13}C$ HSQC spectrum of rivulariapeptolide 1155 in DMSO-*d6*, 500 MHz.

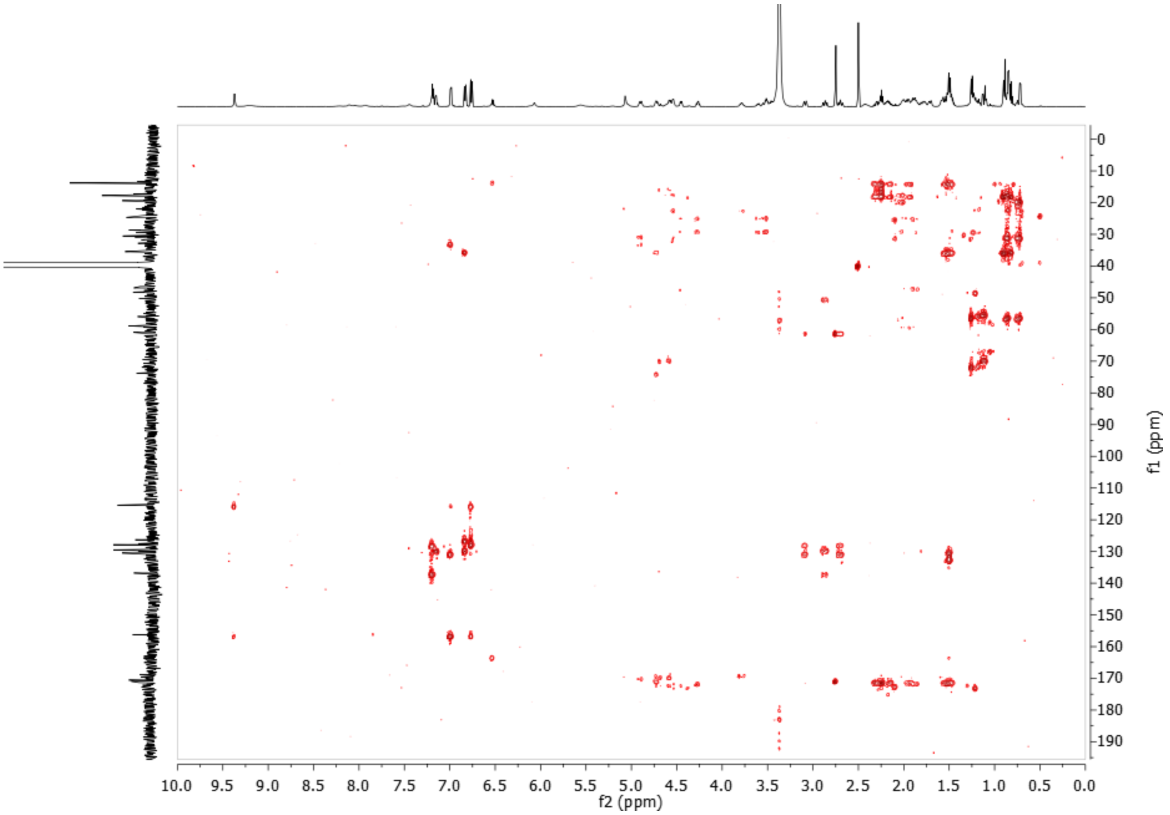


**Figure S16:** $^{1}$H-$^{13}$C HMBC spectrum of rivulariapeptolide 1155 in DMSO-*d6*, 600 MHz.

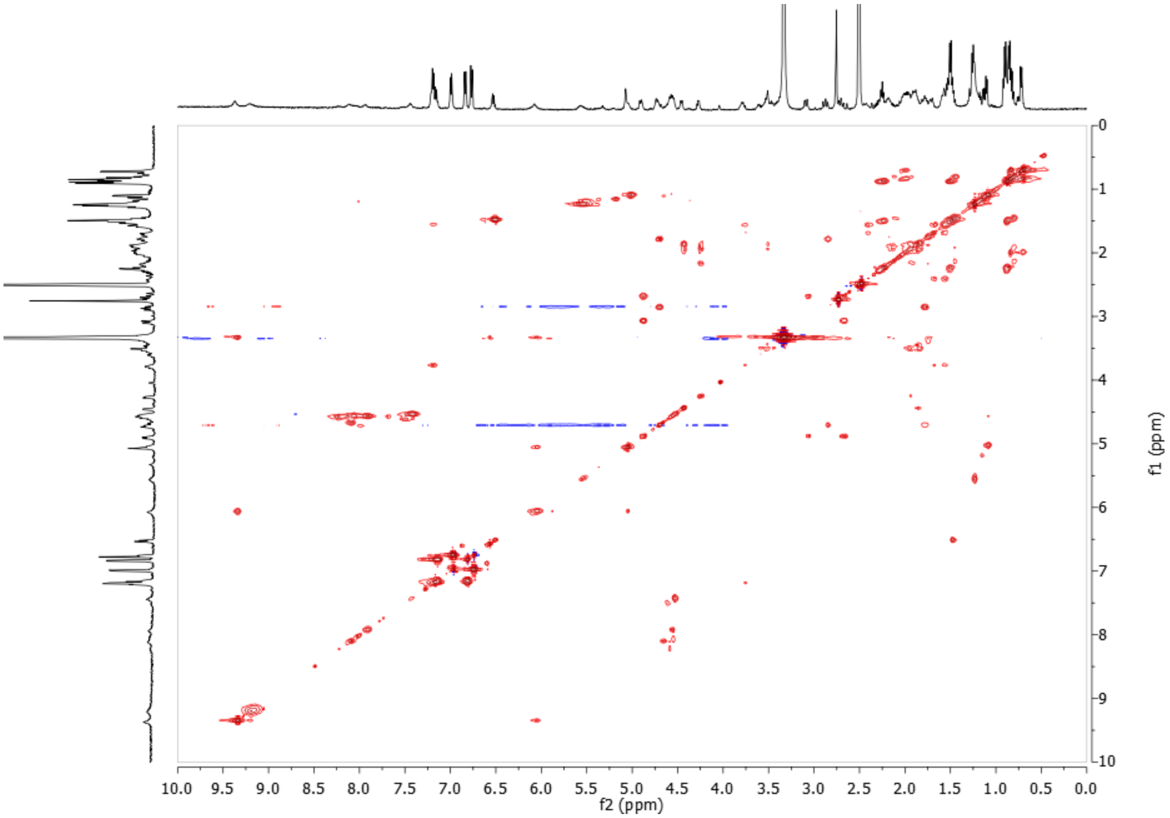


**Figure S17:** $^{1}$H-$^{1}$H TOCSY spectrum of rivulariapeptolide 1155 in DMSO-*d6*, 500 MHz.

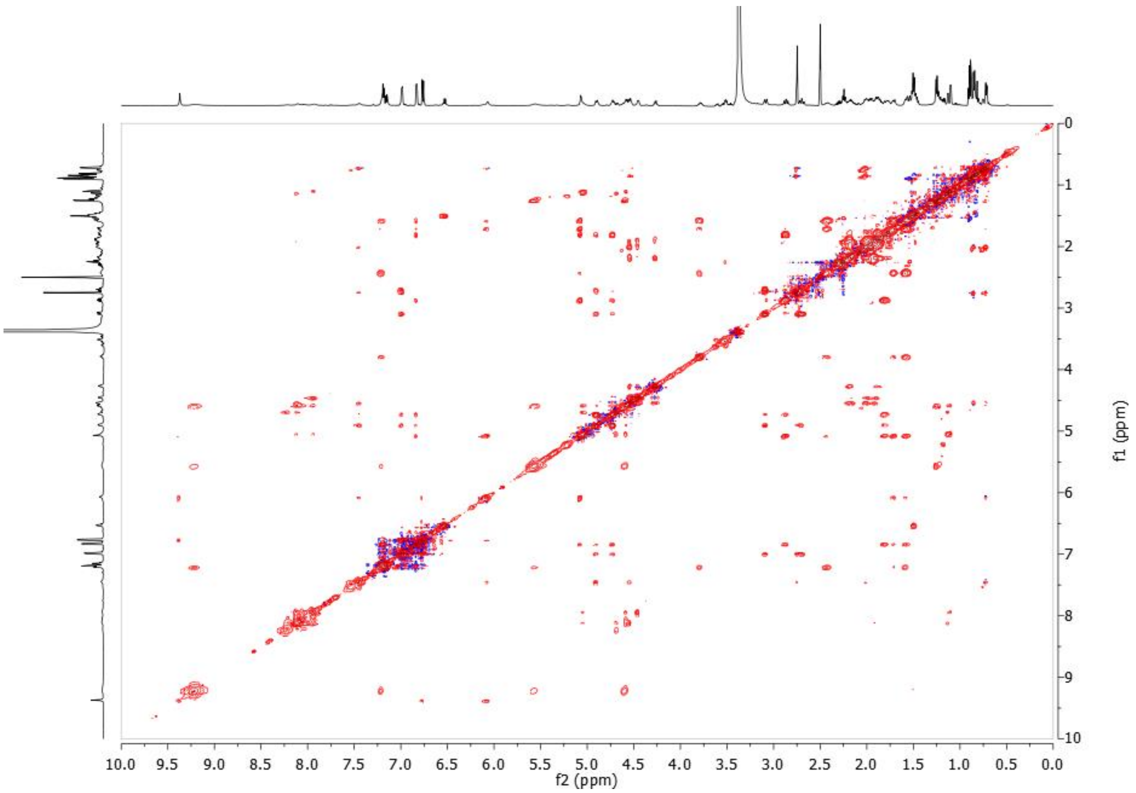


**Figure S18:** $^{1}$H-$^{1}$H NOESY spectrum of rivulariapeptolide 1155 in DMSO-*d6*, 600 MHz.

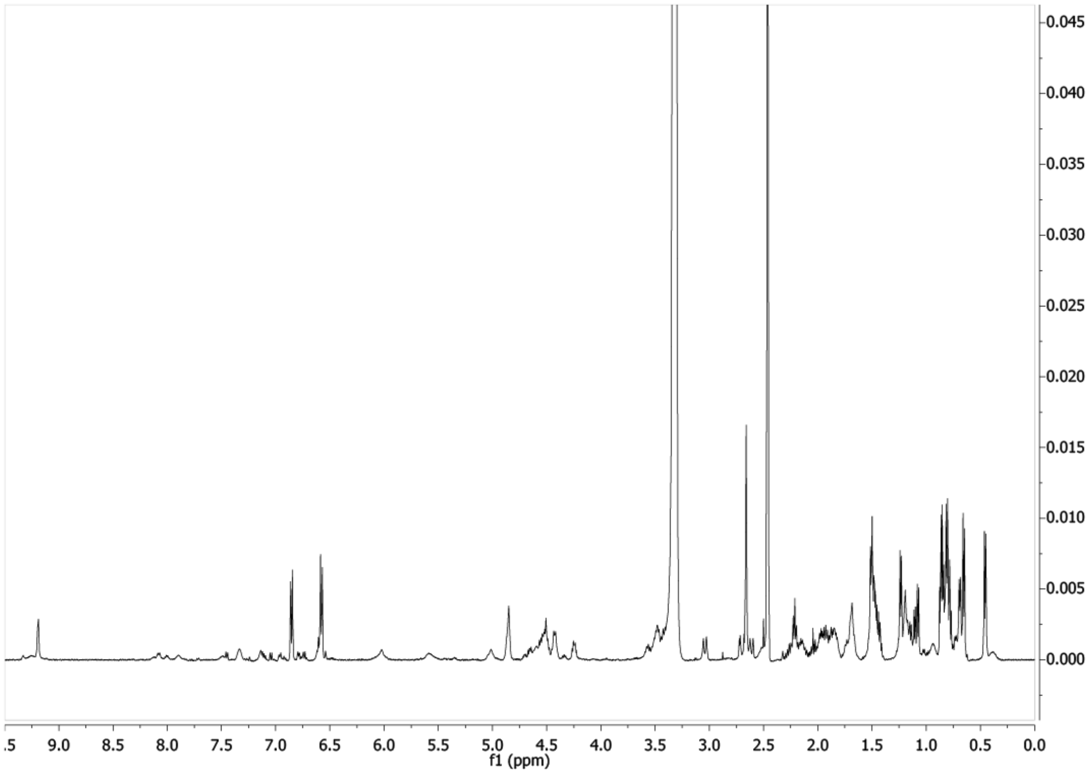


**Figure S19:** $^{1}$H NMR spectrum of rivulariapeptolide 1121 in DMSO-*d6*, 600 MHz.

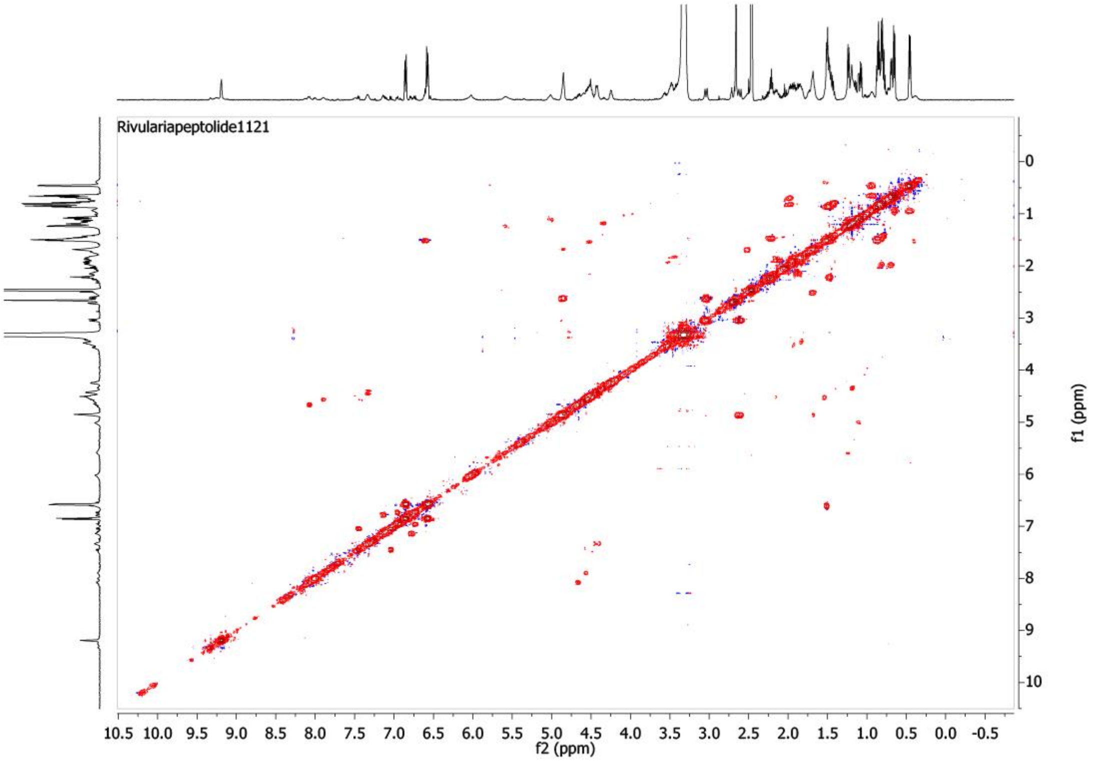


**Figure S20:** $^{1}H$-$^{1}H$ COSY spectrum of rivulariapeptolide 1121 in DMSO-*d6*, 600 MHz.

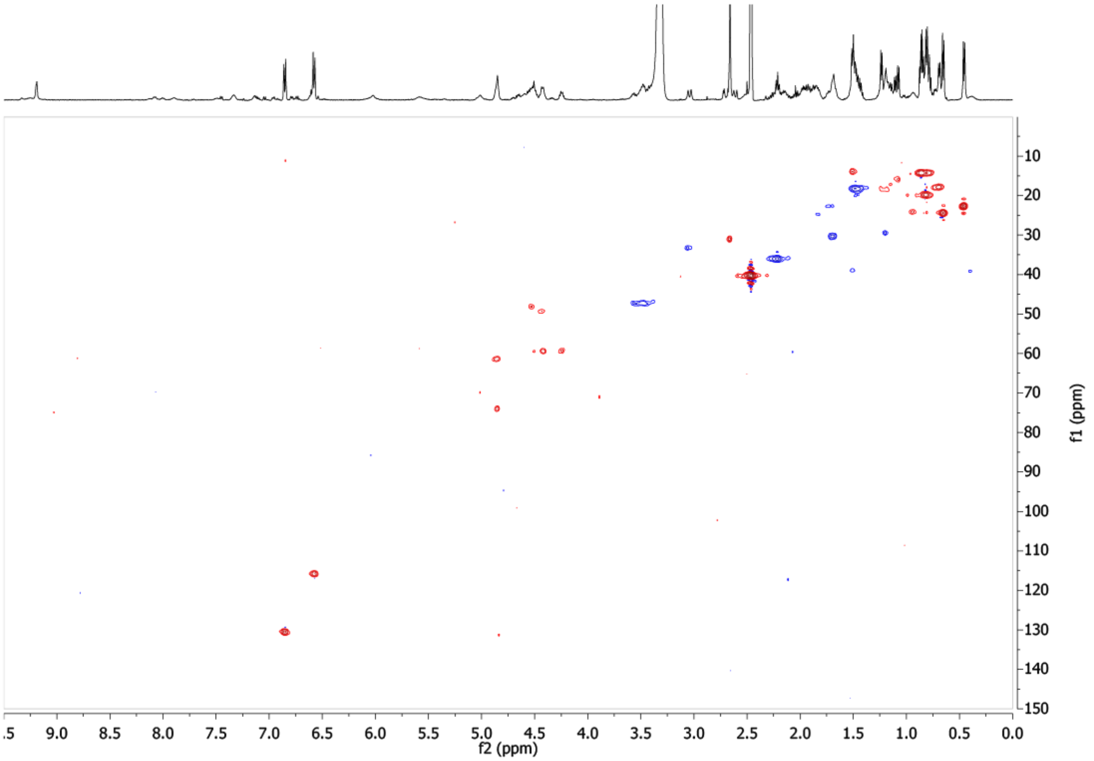


**Figure S21:** $^{1}H$-$^{13}C$ HSQC spectrum of rivulariapeptolide 1121 in DMSO-*d6*, 600 MHz.

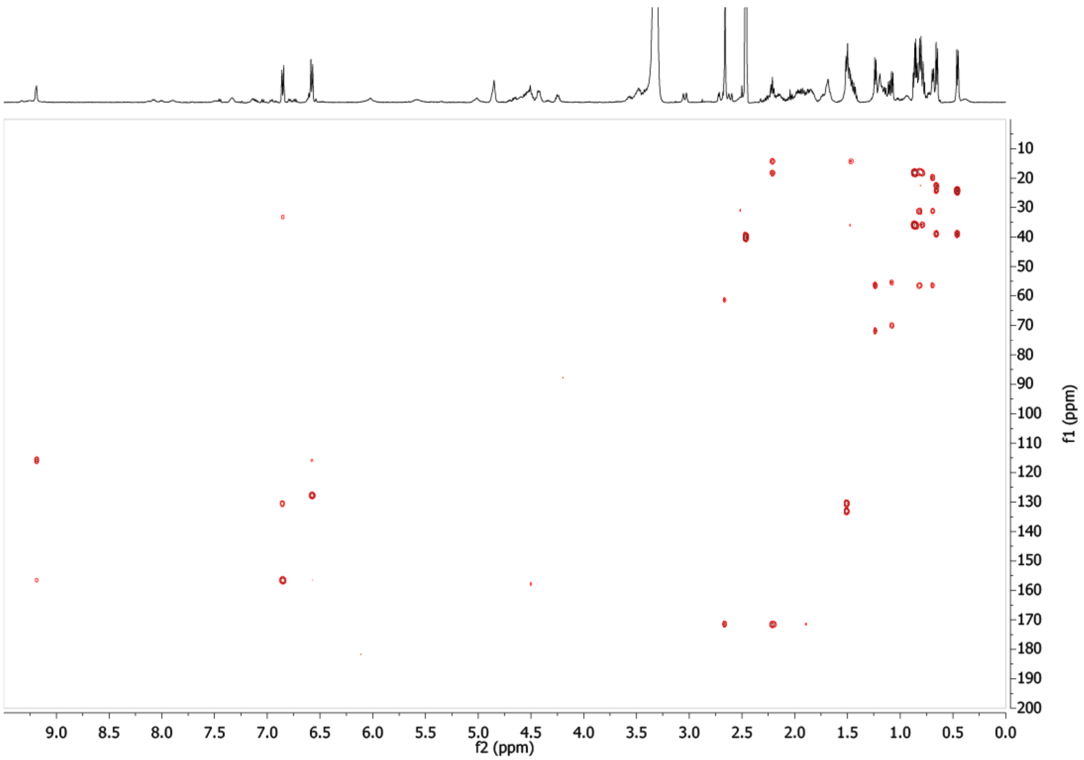


**Figure S22:** $^{1}H$-$^{13}C$ HMBC spectrum of rivulariapeptolide 1121 in DMSO-*d6*, 600 MHz.

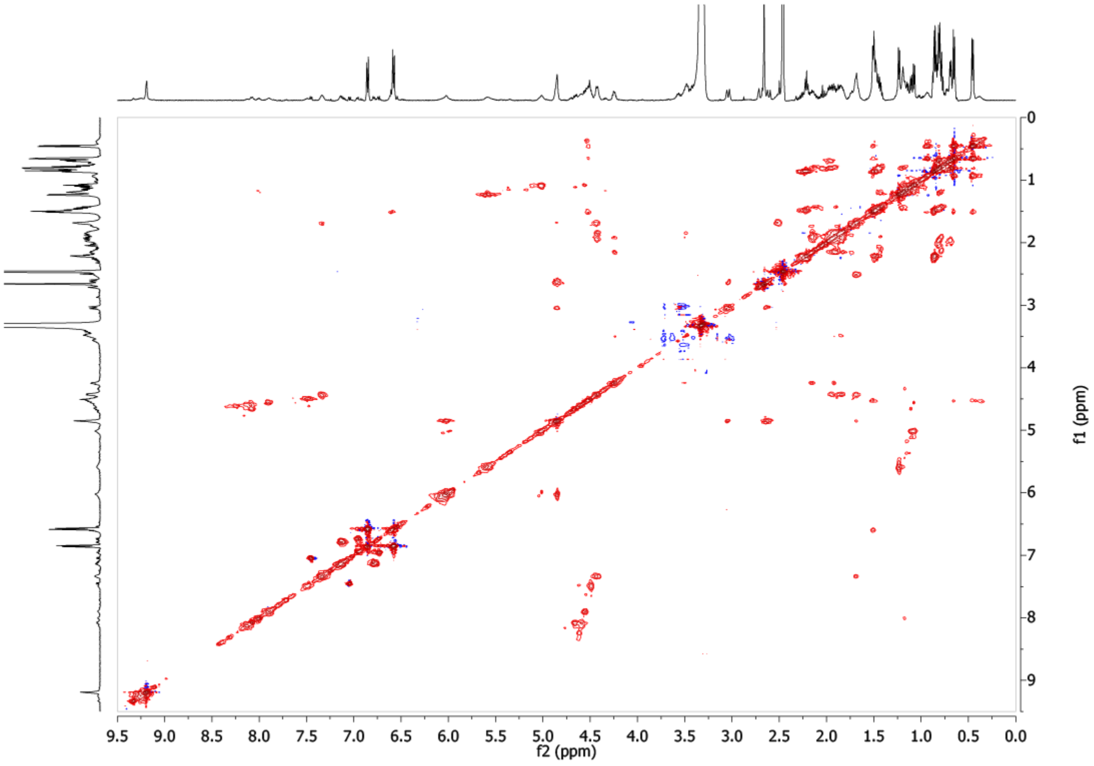


**Figure S23:** $^{1}H$-$^{1}H$ TOCSY spectrum of rivulariapeptolide 1121 in DMSO-*d6*, 600 MHz.

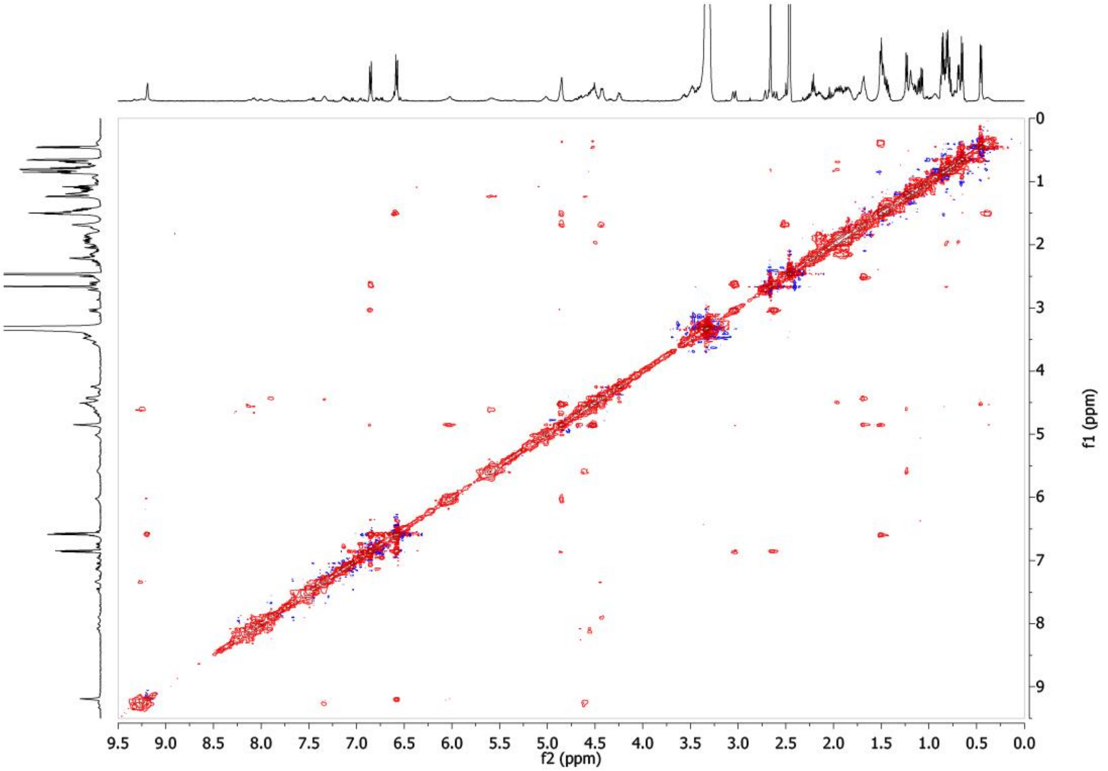


**Figure S24:** $^{1}$H-$^{1}$H NOESY spectrum of rivulariapeptolide 1121 in DMSO-*d6*, 600 MHz.

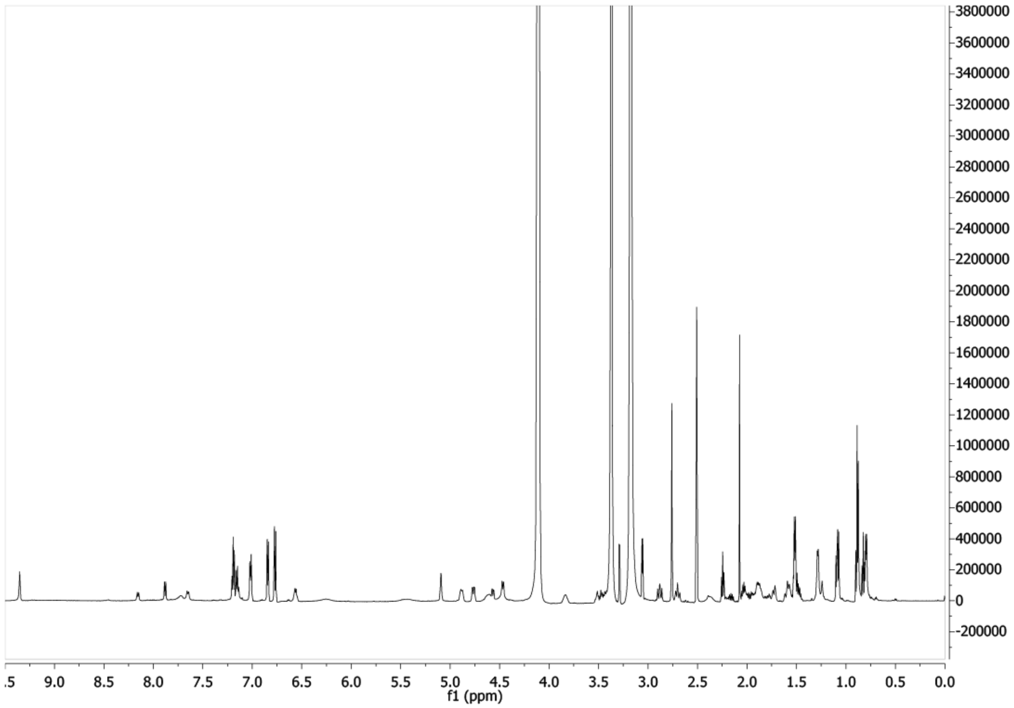


**Figure S25:** $^{1}$H NMR spectrum of rivulariapeptolide 988 in DMSO-*d6*, 600 MHz.

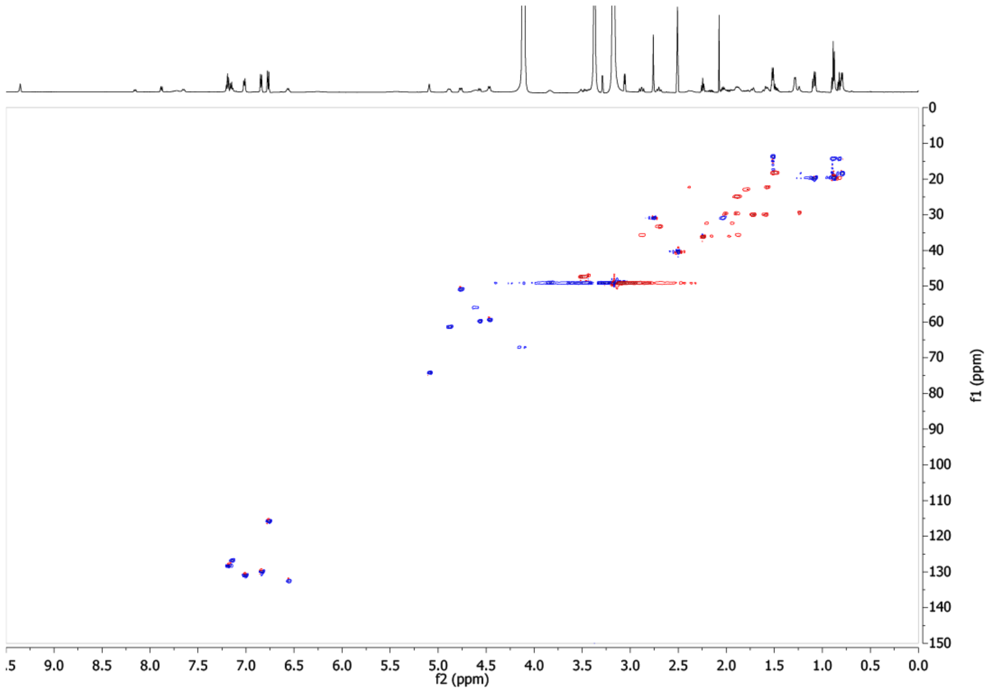


**Figure S26:** $^{1}$H-$^{13}$C HSQC spectrum of rivulariapeptolide 988 in DMSO-*d6*, 600 MHz.

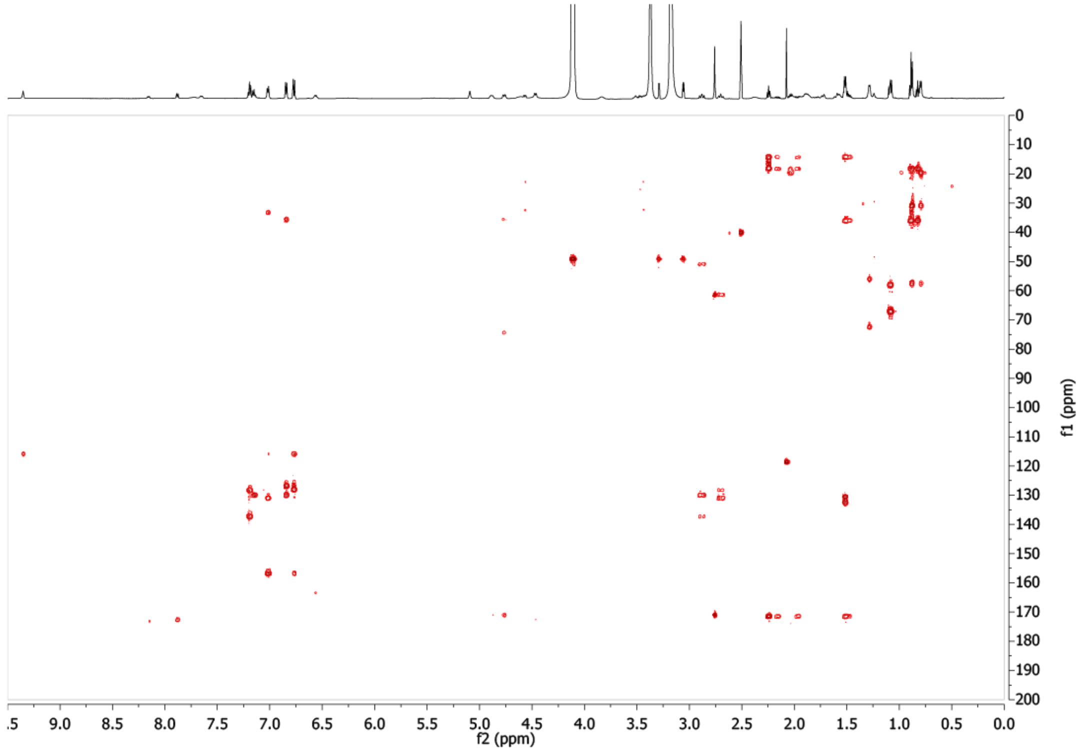


**Figure S27:** $^{1}$H-$^{13}$C HMBC spectrum of rivulariapeptolide 988 in DMSO-*d6*, 600 MHz.

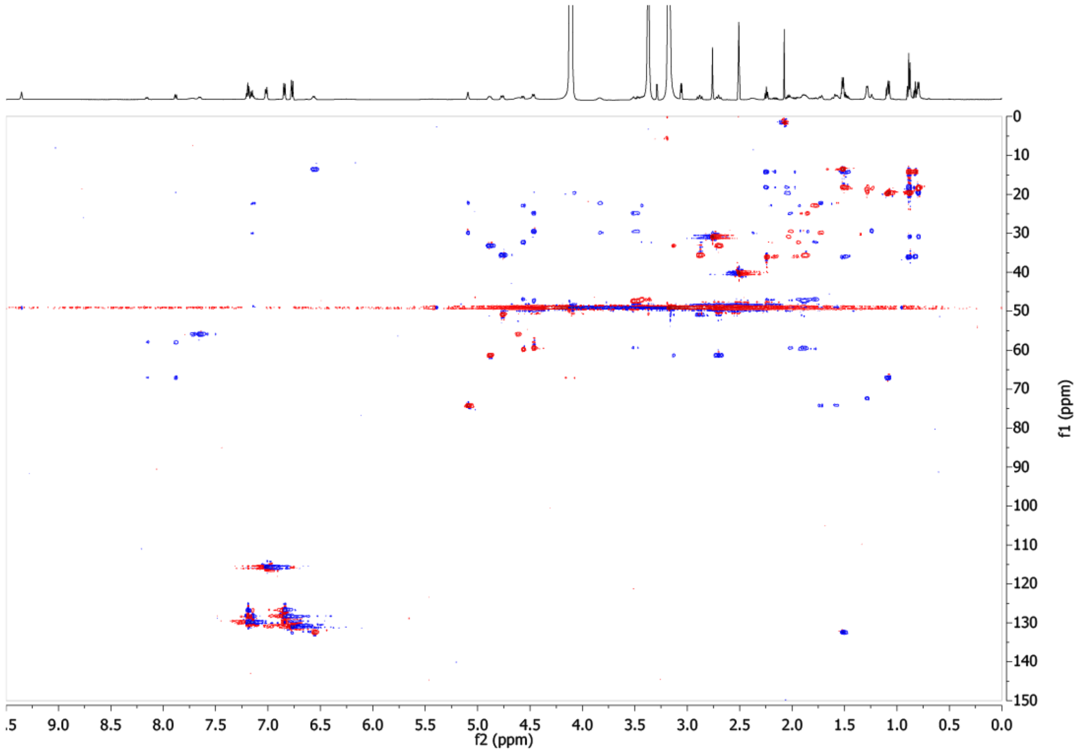


**Figure S28:** $^1$H-$^{13}$C HSQC-TOCSY spectrum of rivulariapeptolide 988 in DMSO-*d6*, 600 MHz.

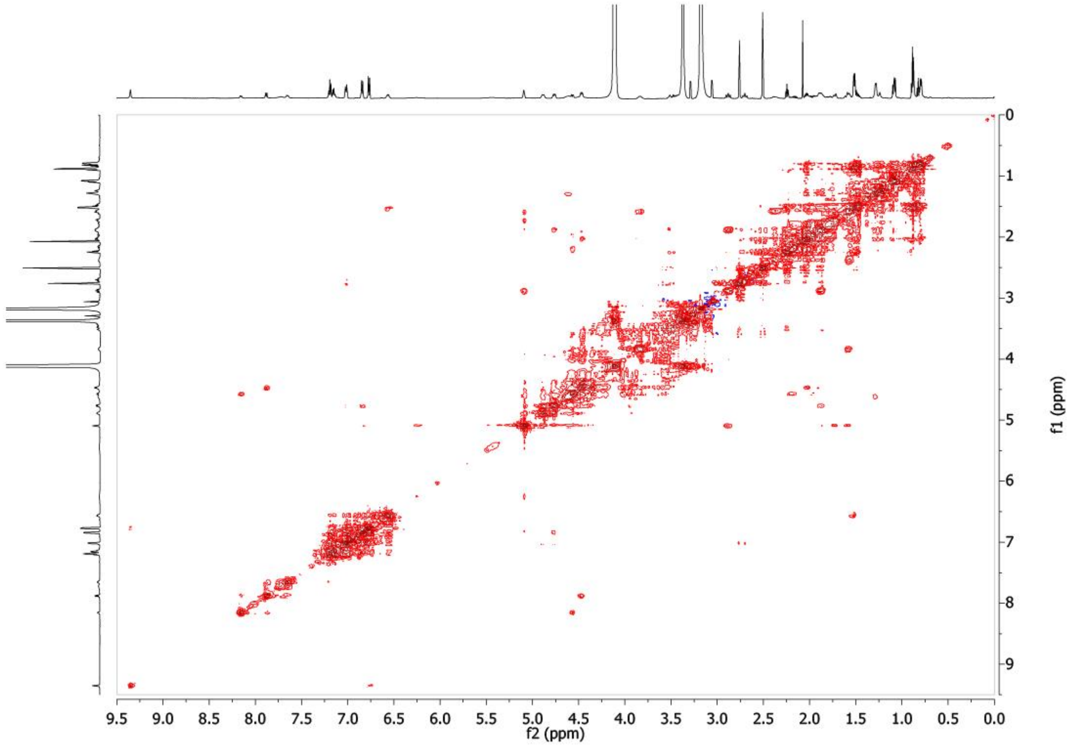


**Figure S29:** $^1$H-$^1$H ROESY spectrum of rivulariapeptolide 988 in DMSO-*d6*, 600 MHz.

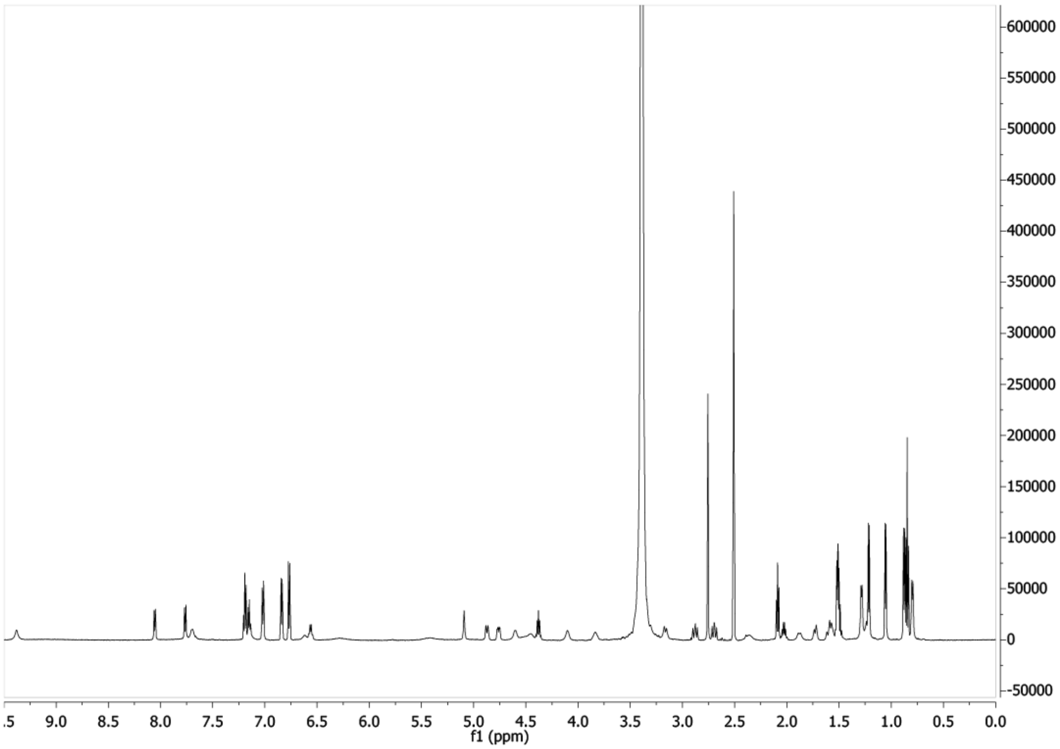


**Figure S30:** $^{1}$H NMR spectrum of molassamide (**5**) in DMSO-*d6*, 600 MHz.

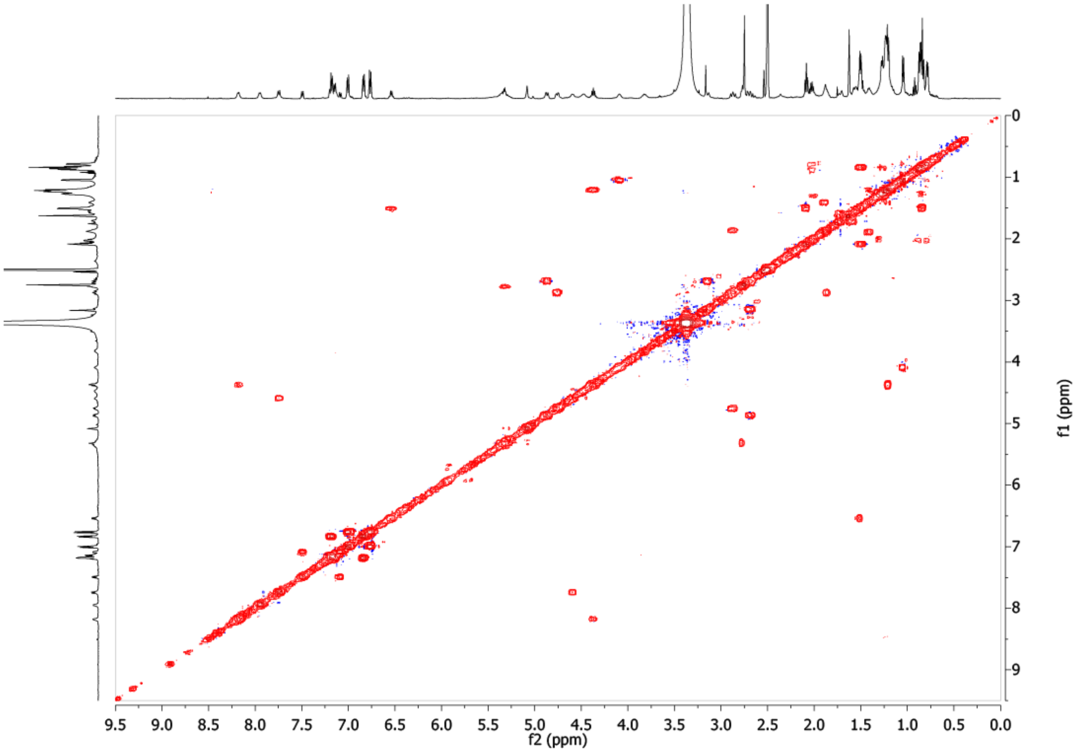


**Figure S31:** $^{1}$H-$^{1}$H COSY spectrum of molassamide in DMSO-*d6*, 600 MHz.

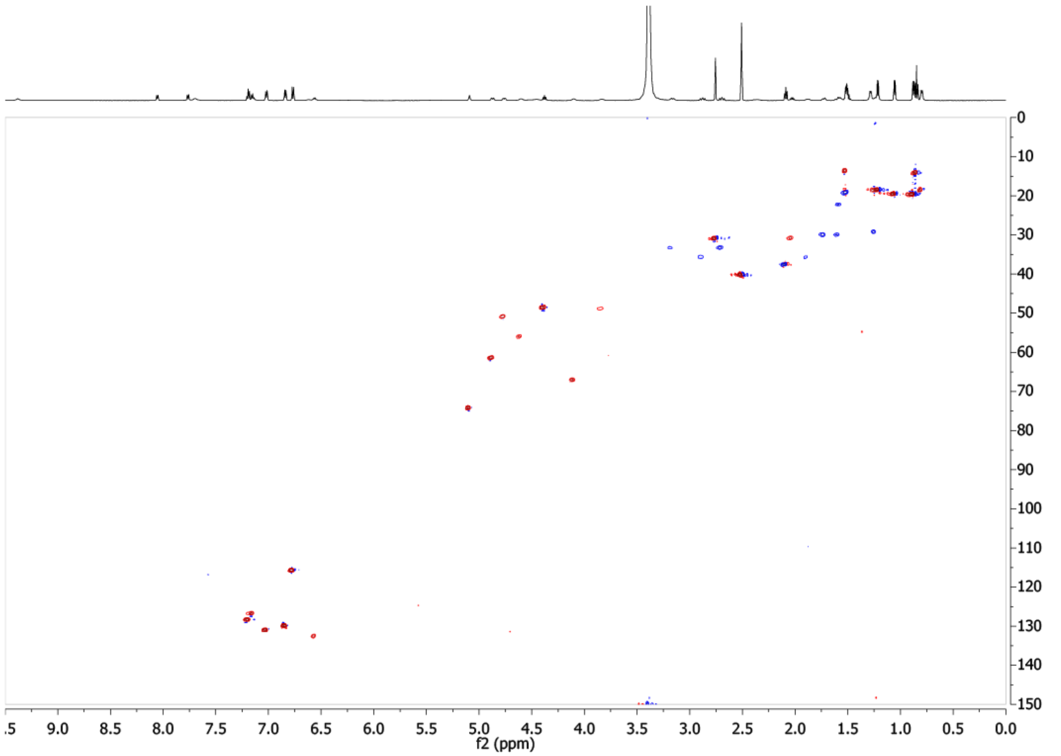


**Figure S32:** $^1$H-$^{13}$C HSQC spectrum of molassamide in DMSO-*d6*, 600 MHz.

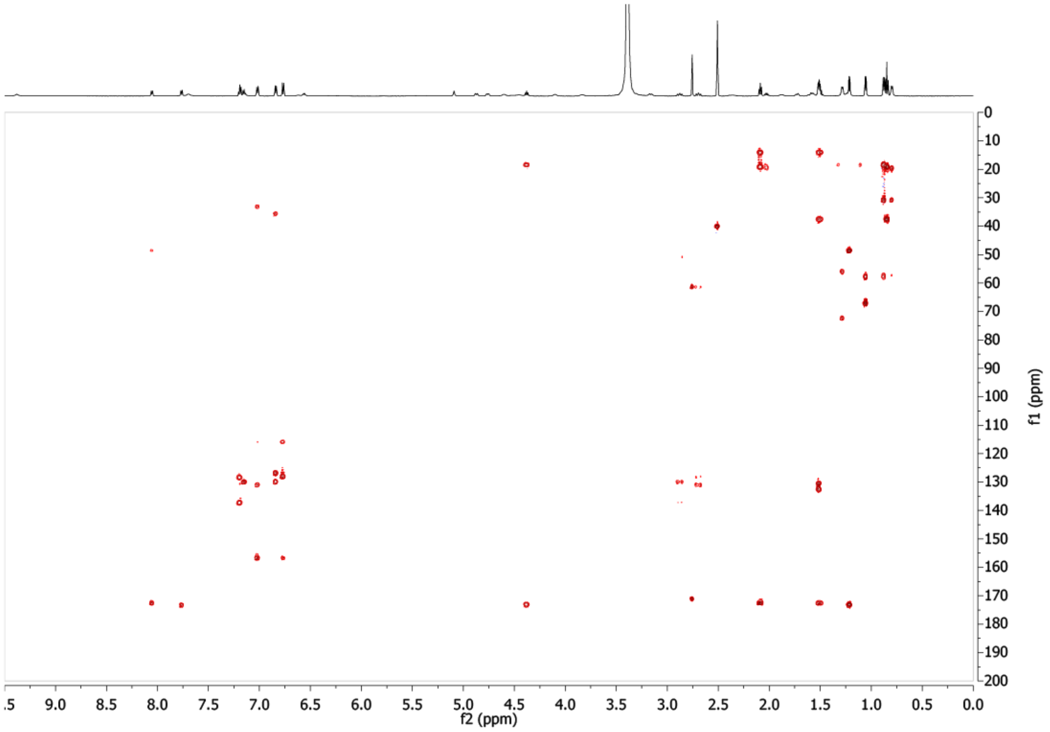


**Figure S33:** $^1$H-$^1$H HMBC spectrum of molassamide  in DMSO-*d6*, 600 MHz.

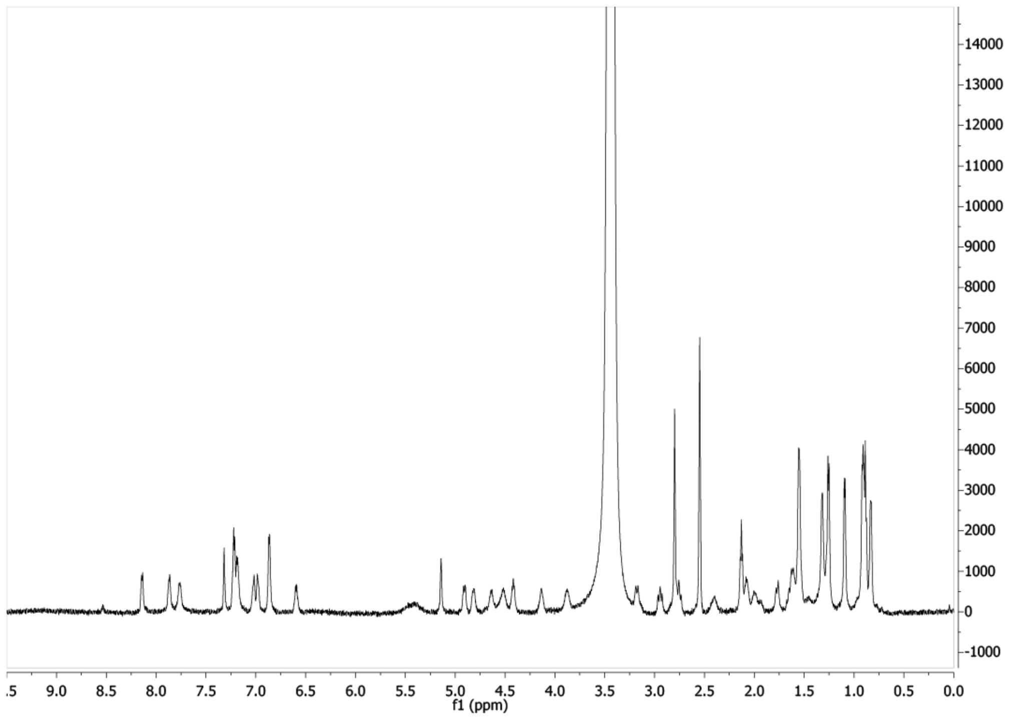


**Figure S34:** $^{1}$H NMR spectrum of molassamide B in DMSO-*d6*, 600 MHz.

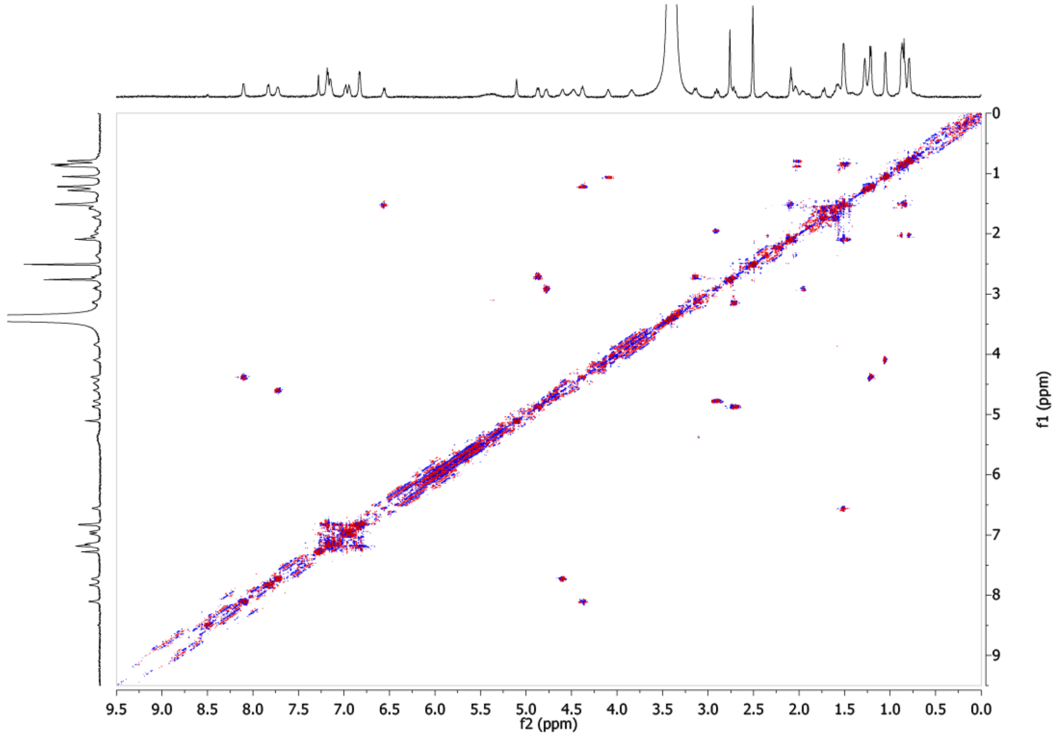


**Figure S35:** $^{1}$H-$^{1}$H COSY spectrum of molassamide B in DMSO-*d6*, 600 MHz.

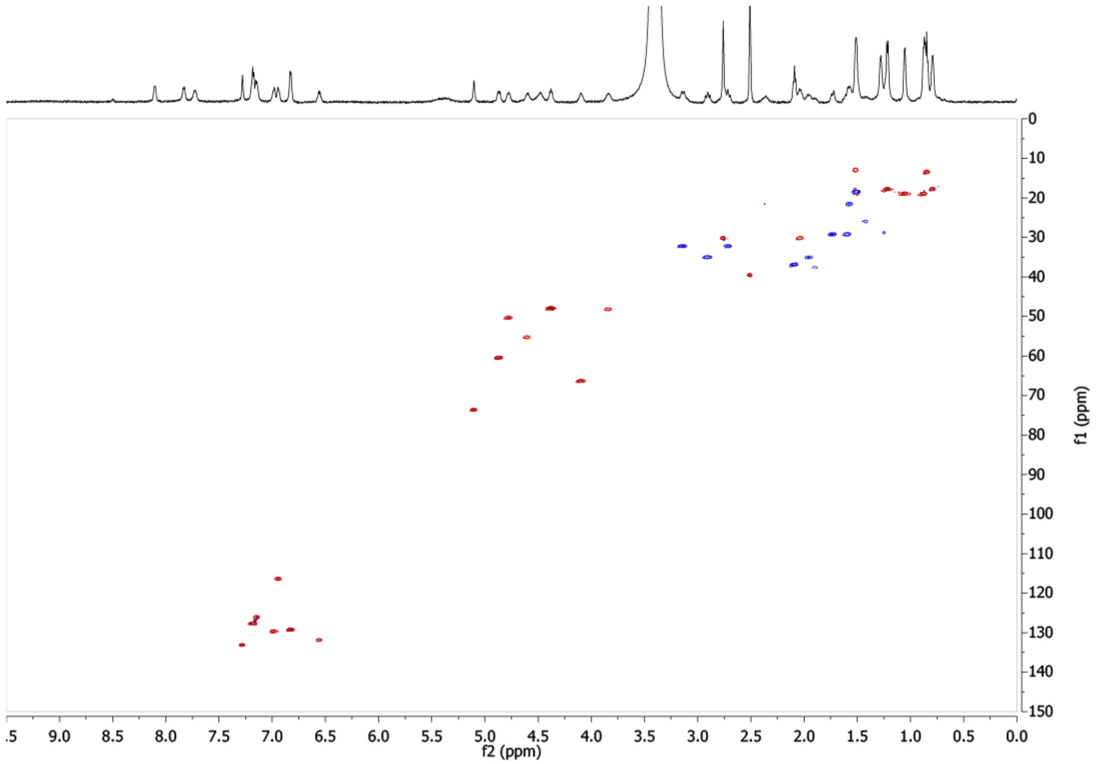


**Figure S36:** $^{1}H$-$^{13}C$ HSQC spectrum of molassamide B in DMSO-*d6*, 600 MHz.

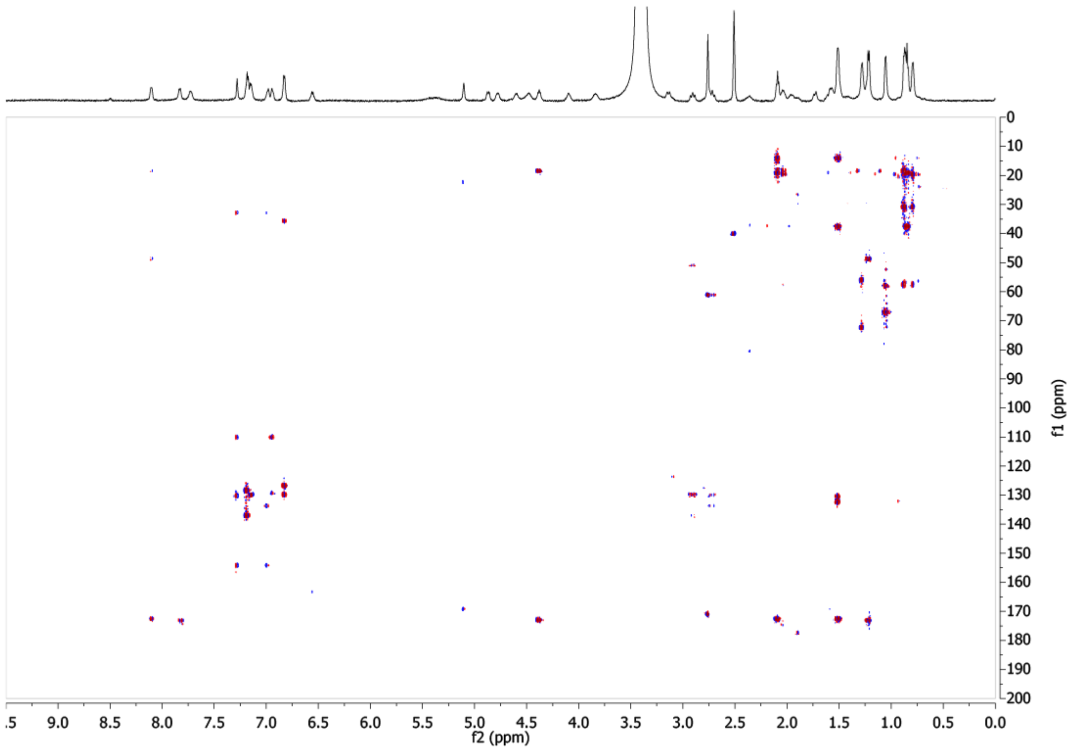


**Figure S37:** $^{1}H$-$^{13}C$ HMBC spectrum of molassamide B in DMSO-*d6*, 600 MHz.

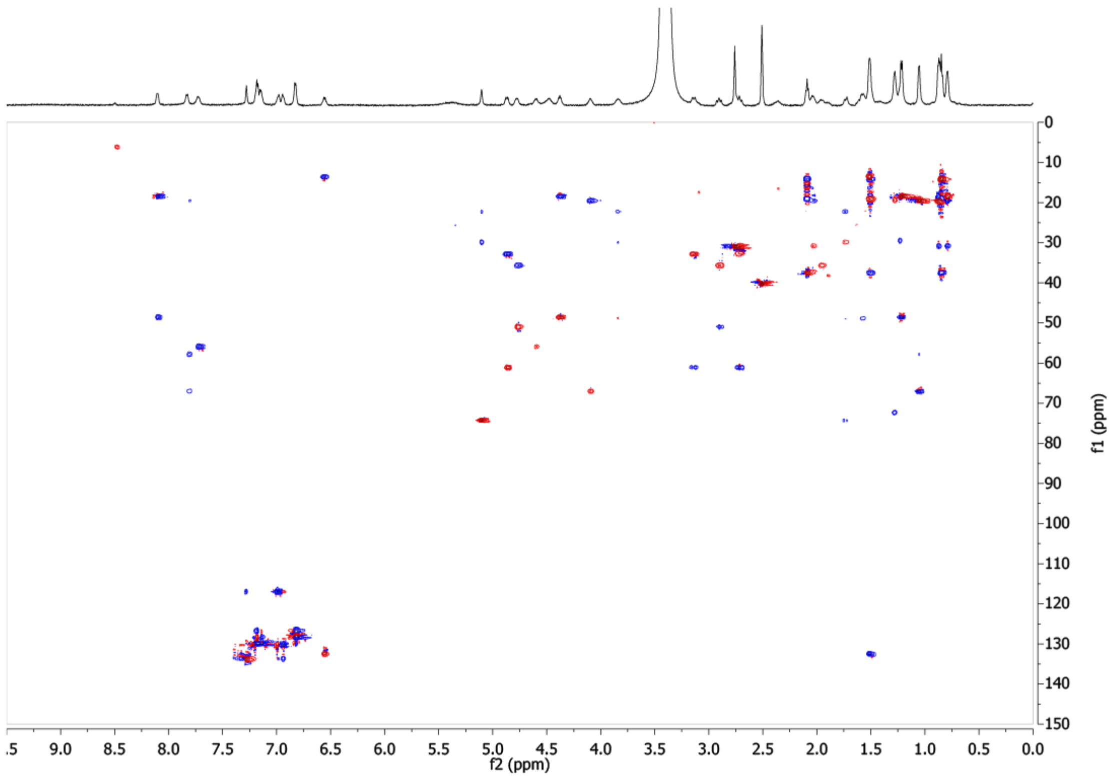


**Figure S38:** $^{1}H$-$^{13}C$ HSQC-TOCSY spectrum of molassamide B in DMSO-*d6*, 600 MHz.

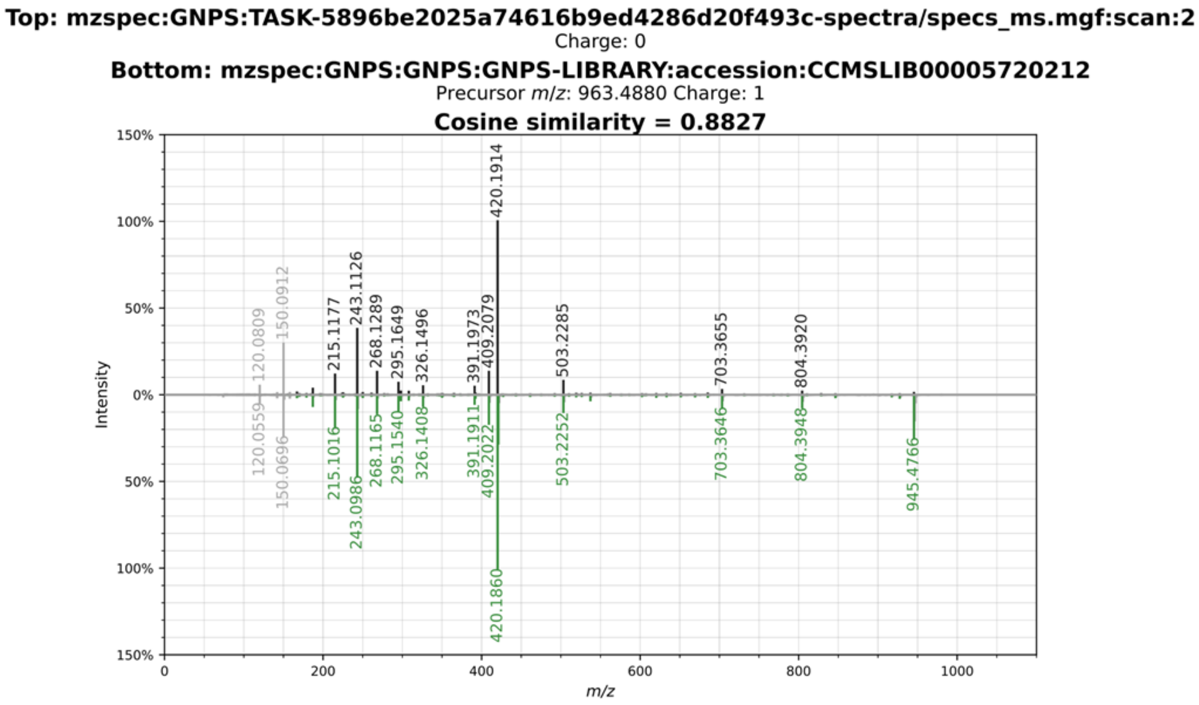


**Figure S39**: Mirror MS² plot of molassamide from this study (black) and GNPS library spectrum (green).

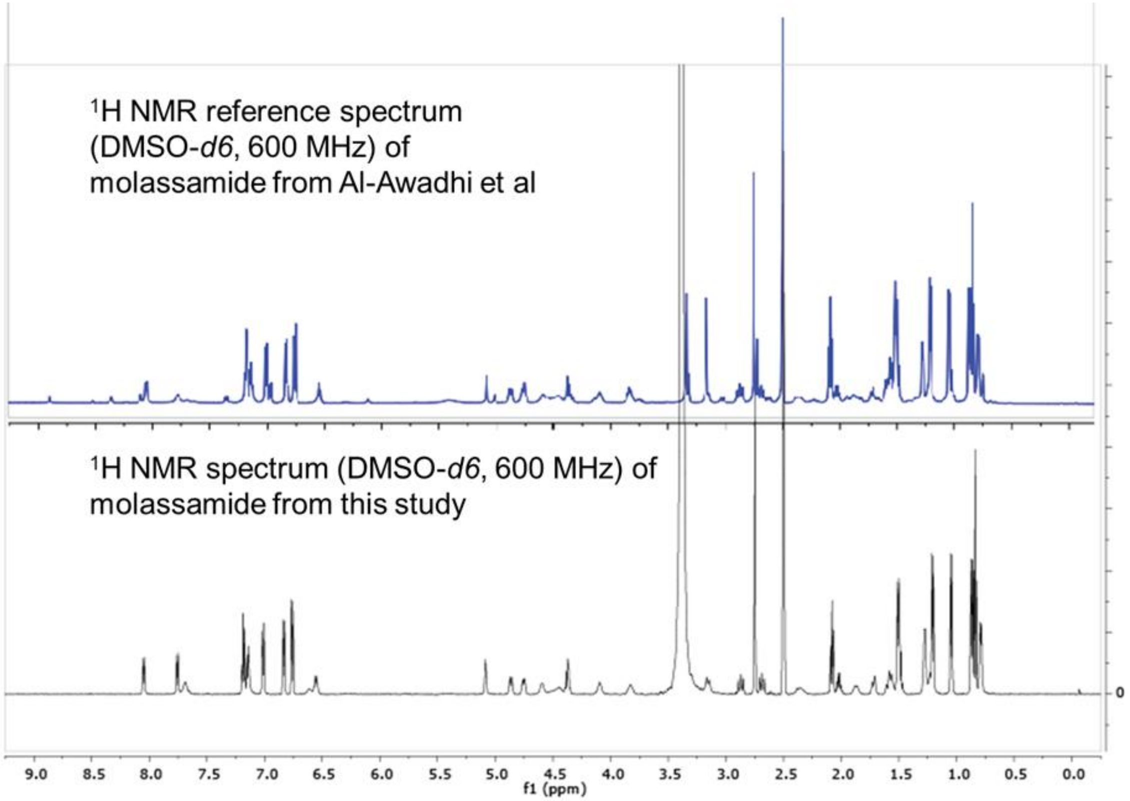


**Figure S40**: Comparison of 1H spectra of molassamide from this study and molassamide isolated by Al-Awadhi et al.[10]

**Rivulariapeptolide 1155**

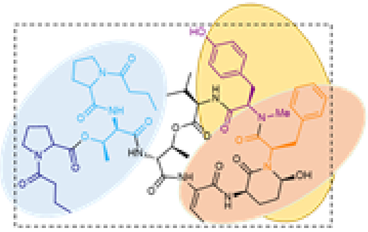


**Rivulariapeptolide 1185**

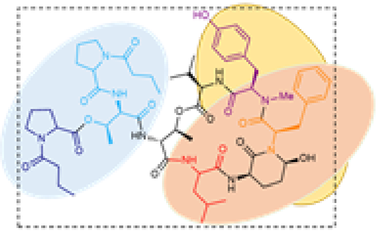


**Rivulariapeptolide 1121**

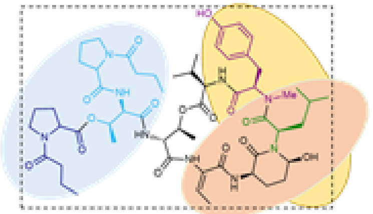


| AA code | Immonium ion (calcd) | Immonium ion (obsvd) |
|---|---|---|
| Phe | 120.0808 | 120.081 |
| N-Me-Tyr | 150.0913 | 150.0914 |
| N-Bu-Pro | 168.1019 | 168.1019 |
| N-Bu-Pro-CO | 140.1070 | 140.1070 |
| Leu | 86.0964 | n.d. |
| Ahp-Phe+CO-OH | 215.1179 | 215.1179 |
| Thr-N-Bu-Pro | 241.1547 | 241.1546 |
| Ahp-Phe-OH | 243.1128 | 243.1126 |
| Ahp-OH-Phe-N-Me-Tyr | 420.1918 | 420.1917 |
| (N-Bu-Pro)$_2$-Thr | 436.2442 | 436.2441 |
| Abu-Ahp-OH-Phe | 326.1499 | 326.15 |
| N-Me-Tyr-Phe-Ahp-Abu | 503.2289 | 503.23 |
| Abu-Thr-OH-Thr-(N-Bu-Pro)2 | 602.3184 | 602.3167 |
| Thr-Val-N-Me-Tyr-Phe-Ahp-Abu | 703.345 | 703.3461 |
| Abu-Thr-OH-Thr-((N-Bu-Pro)2)-Val | 719.3974 | 719.3947 |

| AA code | Immonium ion (calcd) | Immonium ion (obsvd) |
|---|---|---|
| Phe | 120.0808 | 120.0811 |
| N-Me-Tyr | 150.0913 | 150.0914 |
| N-Bu-Pro | 168.1019 | 168.1019 |
| N-Bu-Pro-CO | 140.1070 | 140.1070 |
| **Leu** | 86.0964 | **86.0971** |
| Ahp-Phe+CO-OH | 215.1179 | 215.1174 |
| Thr-N-Bu-Pro | 241.1547 | 241.1547 |
| Ahp-Phe-OH | 243.1128 | 243.1128 |
| Ahp-OH-Phe-N-Me-Tyr | 420.1918 | 420.1924 |
| (N-Bu-Pro)2-Thr | 436.2442 | 436.244 |
| **Leu**-Ahp-OH-Phe | 356.1969 | **356.1965** |
| **Leu**-Thr-OH-Thr-(N-Bu-Pro)2 | 632.3654 | **632.3650** |
| Thr-Val-N-Me-Tyr-Phe-Ahp-**Leu** | 733.3919 | **733.3907** |

| AA code | Immonium ion (calcd) | Immonium ion obsvd) |
|---|---|---|
| Phe | 120.0808 | **n.d.** |
| N-Me-Tyr | 150.0913 | 150.0914 |
| N-Bu-Pro | 168.1019 | 168.1018 |
| N-Bu-Pro-CO | 140.1070 | 140.1070 |
| **Leu** | 86.0964 | **86.0969** |
| Ahp+CO-OH-**Leu** | 181.1335 | **181.1335** |
| Thr-N-Bu-Pro | 241.1547 | 241.1545 |
| Ahp-OH-**Leu** | 209.1285 | **209.1284** |
| AHP-OH-**Leu**-N-Me-Tyr | 386.2074 | **386.2072** |
| (N-Bu-Pro)$_2$-Thr | 436.2442 | 436.244 |
| Abu-Ahp-OH-**Leu** | 292.1656 | **292.1647** |
| N-Me-Tyr-**Leu**-Ahp-Abu | 469.2445 | **469.2449** |
| Abu-Thr-OH-Thr-(N-Bu-Pro)2 | 602.3184 | 602.3181 |
| Thr-Val-N-Me-Tyr-**Leu**-Ahp-Abu | 669.3606 | 669.3585 |
| Abu-Thr-OH-Thr-((N-Bu-Pro)2)-Val | 719.3974 | 719.4006 |

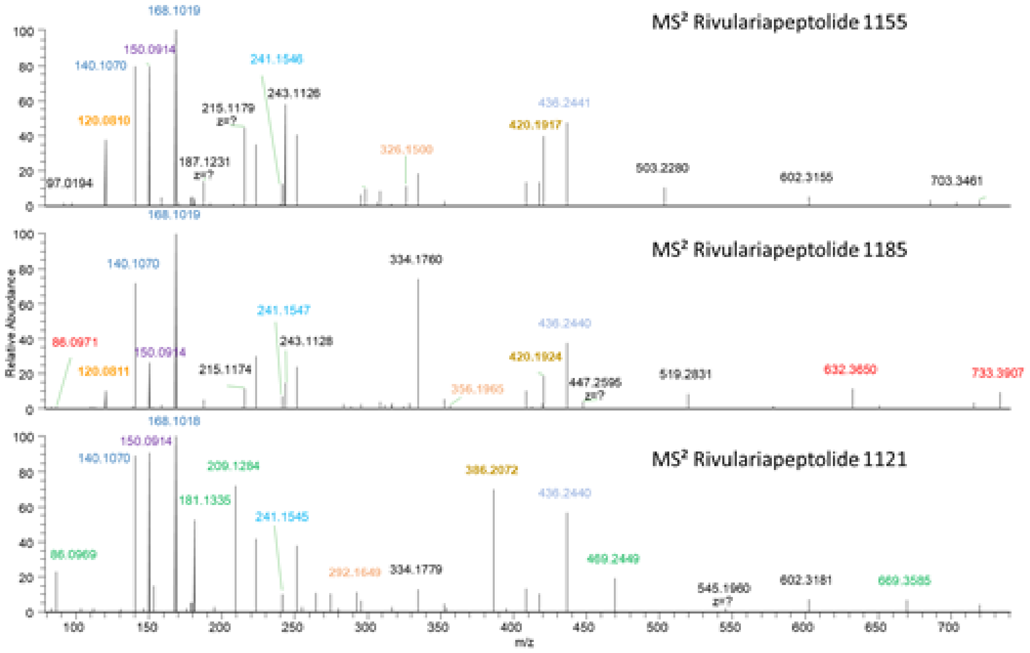


**Figure S41**: MS/MS annotations for rivulariapeptolides 1155, 1185, and 1121. Ions that helped to elucidate the amino acid identity and position for the new rivulariapeptolides 1185, and 1121 are highlighted in red and green, respectively. Rivulariapeptolide 1185 contains Leu instead of Abu in position 5 (refer to Scheme S1) and rivulariapeptolide 1121 contains Leu instead of Phe in position 3, compared with rivulariapeptolide 1155 n.d. = not detected.

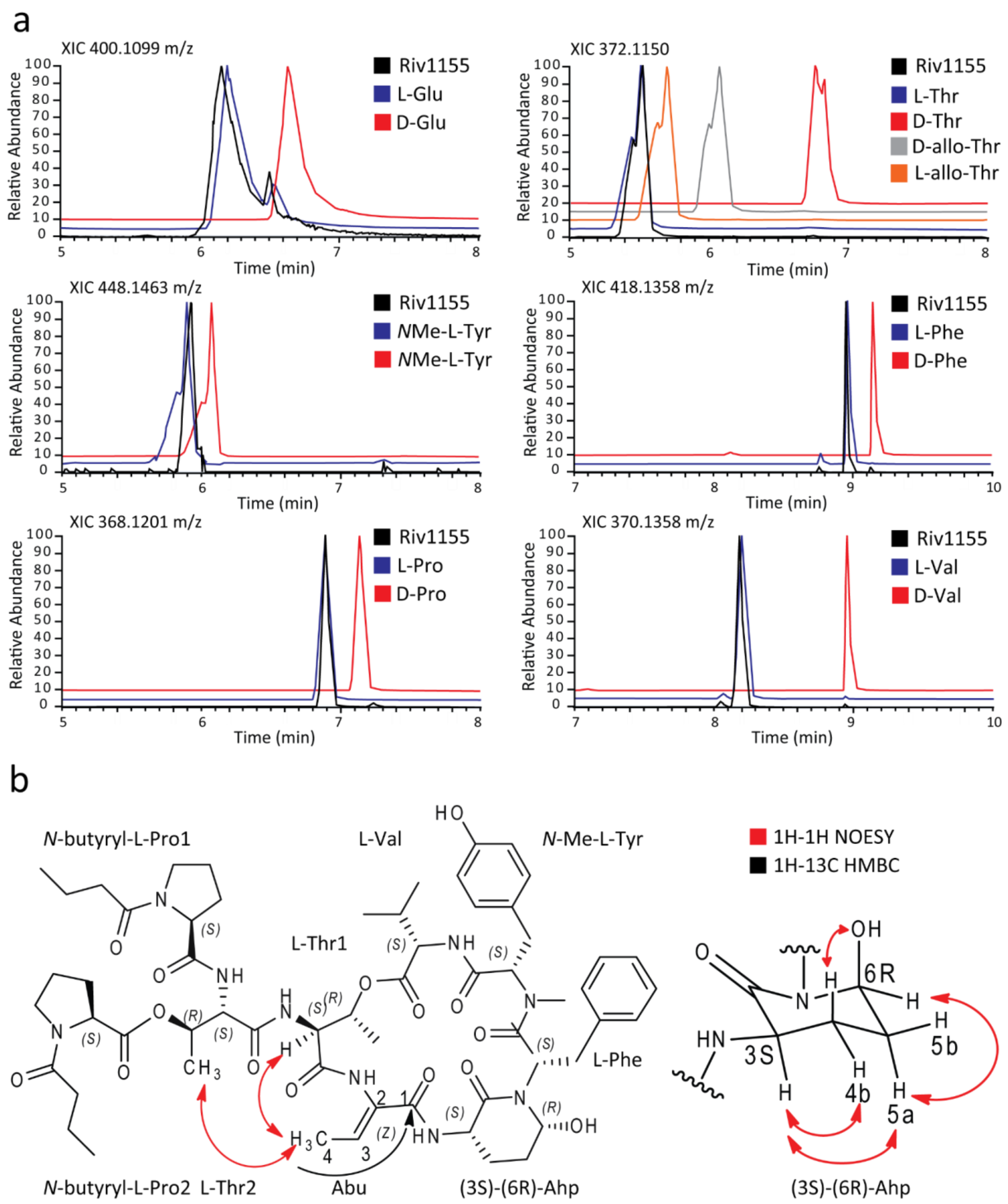


**Figure S42**: Determination of relative and absolute stereochemistry of rivulariapeptolides. (a) XICs of (PDC oxidation), total hydrolysis, and Marfey's derivatization analysis of rivulariapeptolide 1155 compared with in the same way derivatized amino acid standards. (b) Structure of rivulariapeptolide 1155 with assigned stereo configurations, including selected 2D 1H-1H-NOESY and 1H-13C-HMBC NMR correlations used to determine the relative configuration of C-3 and C-6 from the Ahp unit and the geometry of the double bond from the Abu moiety. Abbreviations for amino acids: L-Val = L-valine, *N*-Me-L-Tyr = *N*-methyl-L-tyrosine, L-Phe = L-phenylalanine, L-Ahp = (3S)-amino-(6R)-hydroxy-2-piperidone, Abu = 2-aminobut-2-enoic acid, L-Thr1 = L-threonine1, L-Thr2 = threonine2, *N*-Bu-L-Pro1 = *N*-butyryl-L-proline1, *N*-Bu-L-Pro2 = *N*-butyryl-L-proline2.

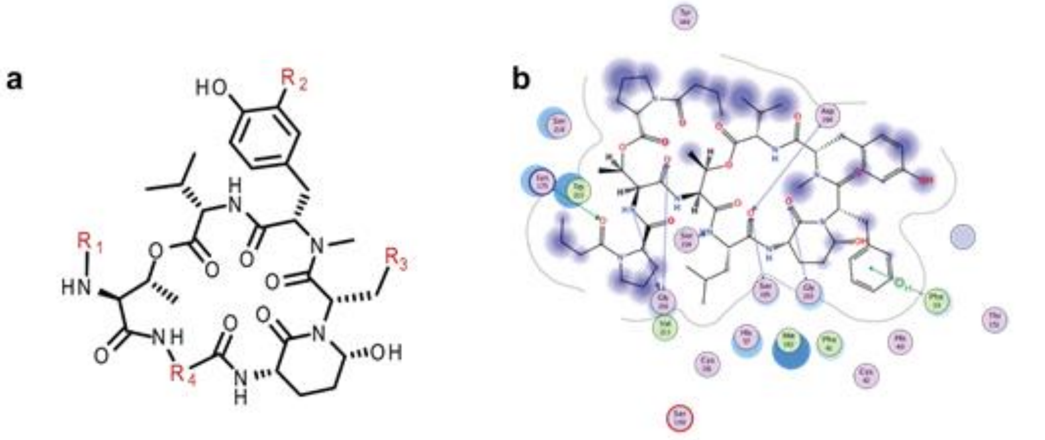


**c**

| Compound | $R_1$ | $R_2$ | $R_3$ | $R_4$ | $IC_{50}$ Chymo-trypsin [nM] | $IC_{50}$ Elastase [nM] | $IC_{50}$ Proteinase K [nM] |
|---|---|---|---|---|---|---|---|
| Rivulariapeptolide 1155 | (*N*-Ba-Pro)$_2$-Thr- | H | Ph | -C=CH-CH$_3$ | 28.2 ± 2.2 | 3.95 ± 0.4 | 39.8 ± 4.1 |
| Rivulariapeptolide 1185 | (*N*-Ba-Pro)$_2$-Thr- | H | Ph | -CH-CH$_2$-iPr | 8.50 ± 0.5 | 19.6 ± 1.3 | 40.5 ± 3.9 |
| Rivulariapeptolide 1121 | (*N*-Ba-Pro)$_2$-Thr- | H | iPr | -C=CH-CH$_3$ | 26.3 ± 3.1 | 10.9 ± 1.4 | 32.6 ± 3.5 |
| Rivulariapeptolide 988 | *N*-Ba-Pro-Thr- | H | Ph | -C=CH-CH$_3$ | 63.6 ± 2.6 | 13.4 ± 2.2 | 58.8 ± 5.6 |
| Molassamide | *N*-Ba-Ala-Thr- | H | Ph | -C=CH-CH$_3$ | 503.2 ± 33.6 | 31.3 ± 2.4 | 15.7 ± 0.6 |
| Molassamide B | *N*-Ba-Ala-Thr- | Br | Ph | -C=CH-CH$_3$ | 16.5 ± 1.0 | 9.1 ± 0.8 | 3.8 ± 0.4 |

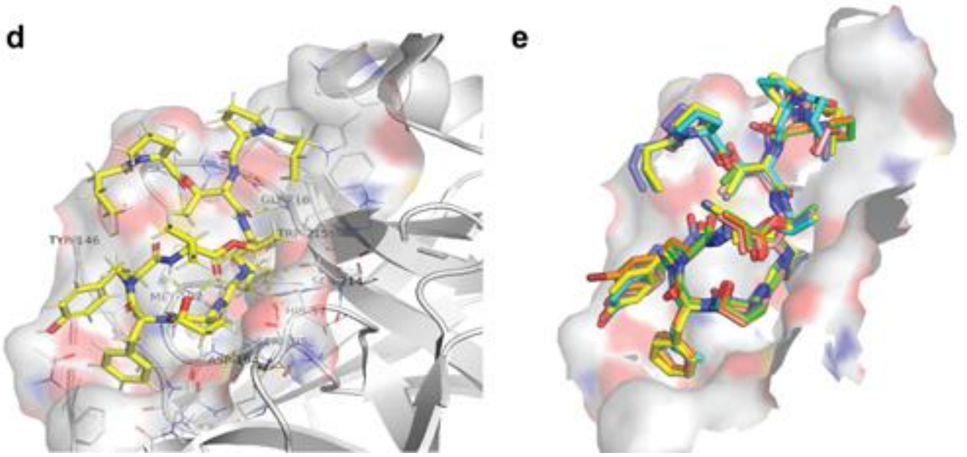


**Figure S43**: (a) Structures of the isolated rivulariapeptolides 1155 (**2**), 1185 (**1**), 1121 (**3**), 988 (**4**), molassamide (**5**), and molassamide B (**6**). (b) Proposed binding sites of **1** in the active site of alpha-chymotrypsin (pdb id 4Q2K) using MOE software. (c) Potency of isolated compounds for selected serine proteases following 40 min pre-incubations. Data are presented as the mean ± SD, n = 3. (d) Induced-fit docking of rivulariapeptolide 1185 inside the binding pocket of alpha-chymotrypsin (pdb id 4Q2K using MOE). The ligand is shown in sticks, the binding pocket as a surface, and the residues of the pocket are shown as lines and labeled. (d) All the isolated compounds compared to one another docked onto the surface of alpha-chymotrypsin (pdb id 4Q2K).

**Table S1**: NMR table for rivulariapeptolide 1185 (**1**) at 600 MHz ($^1$H), 150 MHz ($^{13}$C) in DMSO-*d6* and in MeOH-d4 at 500 MHz ($^1$H), 125 MHz ($^{13}$C), respectively.

| Unit | No. | $\delta_H$ (*J* in Hz)[a] | $\delta_C$[a] | TOCSY | HMBC | NOESY |
|---|---|---|---|---|---|---|
| **Val** | 1 | | 172.0 | | 3 (Thr-1) | |
| | 2 | 4.53, m | 58.8 | NH | 4,5 | 3,4,5, NH |
| | 3 | 2.03, m | 30.6 | 4,5 | 2,3,5 | 2,4,5, NH |
| | 4 | 0.85, d (6.7) | 19.2 | 3 | 2,3,4 | 2, 3 NMe (Tyr) |
| | 5 | 0.69, d (6.9) | 16.9 | 3 | | 2, 3 NMe (Tyr), NH |
| | NH | 8.03, d (7.9) | | | 1 | 2,3,4,5, NMe (Tyr), 2 (Tyr) |
| ***N*-Me-Tyr** | 1 | | 169.4 | | | |
| | 2 | 4.91, d (12.7) | 60.6 | 3b, 3a | 1,3b,N-Me | 1 (Phe), 3a,b (Phe), H(Val) |
| | 3a | 3.08, dd (14.0, 2.1) | 32.6 | 2, 3b | 2, 4, 5/9 | 2, 3b, 5/9 |
| | 3b | 2.71, dd (13.8, 11.8) | 32.6 | 2, 3a | 2, 4, 5/9 | 2,3a,5/9 |
| | 4 | | 127.5 | | | |
| | 5/9 | 7.00, d (8.1) | 130.0 | 6/8 | 7, 3 | 6/8, N-Me (weak), 3b |
| | 6/8 | 6.78, d (8.4) | 115.0 | 5/9 | 4, 7 (weak) | 5/9, 7-OH |
| | 7 | | 156.3 | | | |
| | 7-OH | 9.36, s | | 6,7,8 | 6,7,8 (Tyr) | 6 (AHP), 6/8 (Tyr) |
| | N-Me | 2.76 | 30.1 | | 2, 1(Phe) | 5/9, 4,5,NH (Val) |
| **Phe** | 1 | | 170.5 | | | |
| | 2 | 4.74, dd (11.9, 4.0 | 50.0 | 3a, (3b) | 3a,b, 1,6 (Ahp) | 3b, 3a, 5/9 2, 5/9 (NMe-Tyr) |
| | 3a | 2.86, dd (-13.6, 12.1) | 35.3 | 2, 3b | 2, 4, 5/9 | 2,3b, 5/9, 6 (AHP) |
| | 3b | 1.78, m | 35.3 | 2, 3a | 2, 4, 5/9 | 2,3b, 5/9, 6 (AHP) |
| | 4 | | 136.8 | | | |
| | 5/9 | 6.83, d (7.2) | 129.1 | 6/8, 7 | 3,5/9,7 | 2,3a,3b,6/8 (Phe) 5a,5b,6(Ahp) |
| | 6/8 | 7.18, t (7.2) | 127.4 | 5/9, 7 | 4, 6/8 | 5/9 (Phe)3,4, 5a/b (Ahp) |
| | 7 | 7.14, t (7.2 | 125.8 | 5/9, 6/8 | 5/9 | 5/9 |
| **Ahp** | 2 | | 169.0 | | | |
| | 3 | 3.63, m | 48.2 | 4a,4b, 5a,b NH | 2, 4a,4b | 4a,4b,5a, NH |
| | 4a | 2.41, m | 21.1 | 3, 4b, 5b | | 3, 4b,5a,5b NH |
| | 4b | 1.56, m | 21.1 | 3, 4a, 5a | | 3, 4a5a,5b NH |
| | 5a | 1.68, m | 29.2 | 5b, 6 | | 4a,4b, 5b6, NH |
| | 5b | 1.56, m | 29.2 | 5a,6 | | 5a,6, NH, 5/9 (Phe) |
| | 6 | 5.05, br s | 73.3 | 5a,5b,6-OH | 2 | 5a,5b,6-OH (AhP) 3a,b,5/9 (Phe) |
| | 6-OH | 6.01, br s | | 6 | | |
| | NH | 7.06 | | 3 | 4,5 | 3, 4a/4b, 2 (Leu) |
| **Leu** | 1 | | 171.8 | | | |
| | 2 | 4.23, m | 50.0 | 3a, 3b, 4 | | 3a, 3b, 5, NH (AHP), NH (Thr-1) |
| | 3a | 1.30 | 38.8 | 2, 4, NH | 1,4 | 4, NH |
| | 3b | 1.68 | 38.8 | 2, 4, NH | | |
| | 4 | 1.47 | 23.6 | | | |
| | 5 | 0.84 | 23.2 | 2, 3a, 3b, 5 | 1,2,3 | 3, NH |
| | 6 | 0.72, d (6.5) | 20.4 | | | |
| | NH | 8.35 | | 2, 3a, 3b | | 2,3 (Thr-1), NH (Ahp) |
| **Thr-1** | 1 | | 169.5 | | NH | |
| | 2 | 4.62, m | 55.4 | NH | | 3,4, NH (Thr-1), 3,4, NH(Thr-2), NH (Abu) |
| | 3 | 5.40, m | 71.4 | 4 | | 2,4, NH (Ahp), NH (Abu) |
| | 4 | 1.19, d (6.6) | 17.3 | 3 | 2,3 | 2,3, NH (Abu), NH (Thr-2) |
| | NH | 7.38 | | 2 | 1 | 2 (Pro-2), 3 (Thr-2), 4 (Pro-1), 4 (Pro-2) |
| **Thr-2** | 1 | | 169.1 | | | |
| | 2 | 4.58, m | 54.7 | 3,4,NH | | 3,4, NH |

| | | | | | | |
|---|---|---|---|---|---|---|
| | 3 | 4.99, m | 69.3 | 2,4, NH | | 2,4, 2 (Val), NH, NH (Thr-1) |
| | 4 | 1.06, d (6.5) | 15.2 | 2,3 | 2,3 | 3, NH, NH(Thr-1) |
| | NH | 7.87 | | 2,3 | | 2, 3, 4 (Thr-2), 2 (Pro-1), 3b (Pro-2) |
| **Pro-1** | 1 | | 171.7 | | | |
| | 2 | 4.45, m | 58.5 | 3a,3b, 4, 5a, 5b | | |
| | 3a | 1.97, m | 28.6 | 2,4 | | |
| | 3b | 1.89, m | 28.6 | 2,4 | | |
| | 4 | 1.88, m | 24.1 | 5a | | |
| | 5a | 3.52, m | 46.5 | 3,4 | 1 | 2,34 |
| | 5b | 3.46, m | 46.5 | 4 | 1,2,4 | |
| **Ba-1** | 1 | | 171.6 | | | |
| | 2 | 2.26, t (7.2) | 35.6 | | 1,3,4 | 5a (Pro-1) |
| | 3 | 1.52, m | 17.5 | | 1,2,4 | |
| | 4 | 0.91, m | 13.8 | | 2,3 | |
| **Pro-2** | 1 | | 170.6 | | | |
| | 2 | 4.28, m | 58.5 | 3a,3b, 4, 5af, 5b | 1(Ba-2) | |
| | 3a | 2.20, m | 28.2 | 2,4 | | |
| | 3b | 2.18, m | 28.2 | 2,4 | | |
| | 4 | 1.97, m | 24.2 | 5a | | |
| | 5a | 3.60, m | 46.4 | 4 | 2,3,4 | |
| | 5b | 2.43, m | 46.4 | 4 | 2,3,4 | |
| **Ba-2** | 1 | | 171.5 | | | |
| | 2 | 2.25, m | 35.1 | | 1,3,4 | 5a (Pro-2) |
| | 3 | 1.49, m | 17.5 | | 1,2,4 | |
| | 4 | 0.84, m | 13.7 | | 2,3 | |

**[**a] Assignments are based on extensive 1D and 2D NMR measurements ($^{1}$H, HMBC, HSQC, COSY, HSQC-TOCSY, NOESY). See also Figures S6-11.

**Table S2***:* NMR table for rivulariapeptolides 1155 (**2**), 1121 (**3**), and 988 (**4**) at 500 MHz and 600 MHz ($^{1}$H), respectively, 125 MHz and 150 MHz for ($^{13}$C) in DMSO-*d6*.

| **Unit** | **No.** | **2 $\delta_H$ [a]** | **2 $\delta_C$[a]** | **3 $\delta_H$ [b]** | **3 $\delta_C$[b]** | **4 $\delta_H$ [b]** | **4 $\delta_H$ [b]** |
|---|---|---|---|---|---|---|---|
| Val | 1 | | 172.4 | | 172.4 | | 172.4 |
| | 2 | 4.54, m | 55.9 | 4.53 | 55.9 | 4.54 | 55.9 |
| | 3 | 2.00, m | 30.8 | 2.03 | 30.4 | 2.03 | 30.1 |
| | 4 | 0.86, d | 19.4 | 0.85 | 19.2 | 0.88 | 18.8 |
| | 5 | 0.73, d | 17.3 | 0.73 | 17.2 | 0.79 | 17.6 |
| | NH | 7.43, br s | | 7.51 | | 7.43 | |
| *N*-Me-Tyr | 1 | | 169.7 | | 169.7 | | 170.4 |
| | 2 | 4.90, d (12.7) | 60.9 | 4.89 | 60.7 | 4.88 | 60.7 |
| | 3a | 3.09, dd (14.0, 2.1) | 32.8 | 3.10 | 32.7 | 3.12 | 32.6 |
| | 3b | 2.70, dd (13.8, 11.8) | 32.8 | 2.66 | 32.7 | 2.70 | 32.6 |
| | 4 | | 127.5 | | 127.1 | | 127.5 |
| | 5/9 | 6.99, d (8.1) | 130.5 | 6.89 | 129.9 | 7.01 | 130.3 |
| | 6/8 | 6.76, d (8.4) | 115.4 | 6.61 | 115.2 | 6.76 | 115.1 |
| | 7 | | 156.3 | | 155.9 | | 156.3 |
| | 7-OH | 9.36, s | | 9.23 | | 9.34 | |
| | N-Me | 2.76 | 30.5 | 2.70 | 30.4 | 2.75 | 30.5 |
| Phe / Leu | 1 | | 170.6 | | 173.6 | | 170.8 |
| | 2 | 4.73, m | 50.3 | 4.47 | 48.7 | 4.76 | 50.1 |
| | 3a | 2.88, dd (-13.6, 12.1) | 35.4 | 1.55 | 38.5 | 2.87 | 34.9 |
| | 3b | 1.81, m | 35.4 | 0.44 | 38.5 | 1.87 | 34.9 |
| | 4 | | 136.8 | 0.98 | 23.6 | | 136.7 |
| | 5/9 | 6.83, d (7.1) | 129.5 | 0.69 | 23.8 | 6.84 | 129.2 |

| | | | | | | | |
|---|---|---|---|---|---|---|---|
| | 6/8 | 7.19, t (7.2) | 127.9 | 0.50 | 22.1 | 7.19 | 127.6 |
| | 7 | 7.15, t (7.2 | 126.3 | | | 7.15 | 126.1 |
| | | | | NH, 6.59 | | | |
| Ahp | 2 | | 168.5 | | 168.5 | | 168.5 |
| | 3 | 3.78, m | 47.3 | 4.57 | 47.6 | 3.83 | 48.3 |
| | 4a | 2.42, m | 22.0 | 1.72 | 22.1 | 2.38 | 21.5 |
| | 4b | 1.58, m | 22.0 | 1.78 | 22.1 | 1.58 | 22.3 |
| | 5a | 1.71, m | 29.4 | 1.73 | 29.7 | 1.71 | 29.8 |
| | 5b | 1.58, m | 29.4 | 1.71, m | 29.7 | 1.58 | 29.8 |
| | 6 | 5.07, br s | 73.8 | 4.89 | 73.3 | 5.08 | 73.5 |
| | 6-OH | 6.07, br s | | 6.06, br s | | 6.24 | |
| | NH | 7.21 | | 7.21 | | 7.21 | |
| Abu | 1 | | 163.6 | | 163.6 | | 163.7 |
| | 2 | | 130.0 | | 129.8 | | 130.2 |
| | 3 | 6.53, q (7.2) | 132.3 | 6.63 | 132.4 | 6.56 | 131.8 |
| | 4 | 1.50, d (7.3) | 13.4 | 1.51 | 13.3 | 1.52 | 13.0 |
| | NH | 9.36 | | 9.36 | | 9.35 | |
| Thr-1 | 1 | | 169.5 | | 169.5 | | 169.5 |
| | 2 | 4.59, m | 55.9 | 4.63 | 55.9 | 4.61 | 55.2 |
| | 3 | 5.53, br s | 71.8 | 5.62 | 71.3 | 5.44 | 71.7 |
| | 4 | 1.26, d (6.5) | 17.8 | 1.29 | 17.6 | 1.26 | 18.1 |
| | NH | 7.93 | | 7.93 | | 7.88 | |
| Thr-2 | 1 | | 169.4 | | 169.4 | | 173.1 |
| | 2 | 4.69, m | 55.0 | 4.70 | 55.1 | 4.69 | 57.7 |
| | 3 | 5.04, br s | 69.6 | 5.05, br s | 69.2 | 5.04 | 66.4 |
| | 4 | 1.11, dd (8.2, 6.4) | 15.6 | 1.11 | 15.3 | 1.09 | 19.0 |
| | NH | 8.11 | | 8.11 | | 8.16 | |
| Pro-1 | 1 | | 171.9 | | 171.9 | | 172.6 |
| | 2 | 4.46, m | 59.0 | 4.46, m | 58.8 | 4.46 | 58.7 |
| | 3a | 1.97, m | 28.8 | 2.01 | 28.9 | 2.00 | 28.8 |
| | 3b | 1.89, m | 28.8 | 1.90 | 28.9 | 1.90 | 28.8 |
| | 4 | 1.88, m | 24.3 | 1.95 | 24.2 | 1.87 | 24.1 |
| | 5a | 3.52, m | 46.9 | 3.51 | 46.6 | 3.51 | 46.7 |
| | 5b | 3.45, m | 46.9 | 3.42 | 46.6 | 3.47 | 46.7 |
| Ba-1 | 1 | | 171.0 | | 170.7 | | 171.1 |
| | 2 | 2.26, t (7.4) | 35.6 | 2.25 | 35.4 | 2.24 | 35.4 |
| | 3 | 1.52, m | 17.8 | 1.52 | 17.6 | 1.51 | 17.4 |
| | 4 | 0.91, m | 13.8 | 0.90 | 13.6 | 0.88 | 13.5 |
| Pro-2 | 1 | | 170.6 | | 170.8 | | |
| | 2 | 4.27, m | 58.9 | 4.28, m | 58.7 | | |
| | 3a | 2.20, m | 28.6 | 2.21 | 28.4 | | |
| | 3b | 2.18, m | 28.6 | 2.19 | 28.4 | | |
| | 4 | 1.97, m | 24.6 | 1.9 | 24.2 | | |
| | 5a | 3.62, m | 46.7 | 3.60 | 46.7 | | |
| | 5b | 2.43, m | 46.7 | 3.37 | 46.7 | | |
| Ba-2 | 1 | | 171.4 | | 171.4 | | |
| | 2 | 2.25, m | 35.5 | 2.25 | 35.4 | | |
| | 3 | 1.49, m | 17.7 | 1.49 | 17.5 | | |
| | 4 | 0.84, m | 13.8 | 0.85 | 13.6 | | |

[a] For assignments of compound **2**, see Dührkop et al[5] **[**b] Assignments are based on extensive 1D and 2D NMR measurements ($^{1}$H, $^{13}$C, HMBC, HSQC, COSY, TOCSY, HSQC-TOCSY). See also Figures S12-S22.

**Table S3**: NMR table for Molassamide (**5**) and Molassamide B (**6**) at 500 MHz and 600 MHz ($^1$H), respectively, 125 MHz and 150 MHz ($^{13}$C) in DMSO-*d6.*

| Unit | No. | Molassamide $\delta_H$ | Molassamide $\delta_C$[a] | Molassamide B $\delta_H$ | Molassamide B $\delta_C$[a] |
|---|---|---|---|---|---|
| Val | 1 | | 172.9 | | 173.0 |
| | 2 | 4.48, br | 57.1 | 4.48 | 57.3 |
| | 3 | 2.02, m | 30.2 | 2.04 | 30.1 |
| | 4 | 0.79, d (6.6) | 17.9 | 0.79 | 17.7 |
| | 5 | 0.87, d (6.9) | 19.1 | 0.88 | 18.9 |
| | NH | 7.75, d (8.2) | | n.d. | |
| *N*-Me-(Br)-Tyr | 1 | | 169.7 | | 169.9 |
| | 2 | 4.87, d (11.3) | 60.8 | 4.87 | 60.6 |
| | 3a | 3.16, d (11.5) | 32.6 | 3.14 | 32.2 |
| | 3b | 2.69, dd () | 32.6 | 2.71 | 32.2 |
| | 4 | | 127.5 | | 133.6 |
| | 5 | 7.01, d (8.4) | 130.4 | 6.99 | 129.6 |
| | 6 | 6.76, d (8.4) | 115.2 | 6.94 | 116.3 |
| | 7 | | 156.3 | | 154.0 |
| | 7-OH | 4.45, br | | 4.47 | |
| | 8 | 6.76, d (8.4) | 115.2 | - | 110.0 |
| | 9 | 7.01, d (8.4) | 130.4 | 7.28 | 133.1 |
| | N-Me | 2.75 | 30.2 | 2.76 | 30.2 |
| Phe | 1 | | 170.6 | | 170.5 |
| | 2 | 4.76, dd (11.2, 4.0) | 50.3 | 4.78 | 50.3 |
| | 3a | 2.87, dd (14.0, 11.8) | 35.0 | 2.90 | 34.9 |
| | 3b | 1.87, d (11.5) | 35.0 | 1.95 | 34.9 |
| | 4 | | 136.8 | | 136.7 |
| | 5/9 | 6.83, d (7.3) | 129.3 | 6.83 | 129.2 |
| | 6/8 | 7.18, t (7.4) | 127.7 | 7.18 | 127.7 |
| | 7 | 7.15, t (7.1) | 126.1 | 7.15 | 126.1 |
| Ahp | 2 | | 168.5 | | 168.7 |
| | 3 | 3.83, br | 48.2 | 3.84 | 48.1 |
| | 4a | 2.37, m | 21.6 | 2.37 | 21.6 |
| | 4b | 1.56, m | 21.6 | 1.57 | 21.6 |
| | 5a | 1.72, m | 29.3 | 1.73 | 29.3 |
| | 5b | 1.58, m | 29.3 | 1.60 | 29.3 |
| | 6 | 5.08, br s | 73.7 | 5.10 | 73.4 |
| | 6-OH | 6.28, br s | | | |
| | NH | 7.13 | | n.d. | |
| Abu | 1 | | 163.6 | | 163.4 |
| | 2 | | 130.0 | | 130.2 |
| | 3 | 6.55, q (7.1) | 131.9 | 6.55 | 131.8 |
| | 4 | 1.51, d | 13.0 | 1.51 | 13.0 |
| | NH | 9.37, br s | | | |
| Thr-1 | 1 | | 169.5 | | |
| | 2 | 4.60, br | 55.4 | 4.60 | 55.2 |
| | 3 | 5.42, br s | 71.8 | 5.41 | 71.8 |
| | 4 | 1.27, d (6.1) | 17.8 | 1.27 | 18.2 |
| | NH | 7.69, br d (6.1) | | 7.72 | |
| Thr-2 | 1 | | 169.4 | | |
| | 2 | 4.39, br | 57.7 | 4.39 | 57.6 |
| | 3 | 4.09, br | 66.5 | 4.10 | 66.4 |
| | 4 | 1.05, d (6.4) | 18.9 | 1.05 | 18.8 |
| | NH | 7.81, br | | 7.83 | |
| Ala | 1 | | 173.0 | | 172.5 |
| | 2 | 4.37, m | 47.9 | 4.38 | 47.9 |
| | 3 | 1.21, d (7.1) | 17.9 | 1.22 | 17.8 |

| | | | | | |
|---|---|---|---|---|---|
| | NH | 8.05, d (7.5) | | 8.10 | |
| Ba | 1 | | 172.5 | | 172.0 |
| | 2 | 2.08, t (7.2) | 37.0 | 2.08 | 36.8 |
| | 3 | 1.51, m | 18.5 | 1.51 | 18.5 |
| | 4 | 0.84, t (7.3) | 13.5 | 0.85 | 13.4 |

[a] Assignments are based on extensive 1D and 2D NMR measurements ($^{1}H$, HMBC, HSQC, COSY, HSQC-TOCSY). See also Figures S23-S31, n.d. = not detected.

**Table S4:** Top 50 Chymotrypsin-Inhibitors among Ahp-cyclodepsipeptides.

| Rank | NAME(S) | AA1 | AA2 | AA3 | AA4 | AA5 | AA6 | AA7 | AA8 | AA9 | Chymotrypsin $IC_{50}$ [µM] |
|---|---|---|---|---|---|---|---|---|---|---|---|
| 1 | Micropeptin T-20 | | | Hpg | Thr | Phe | Ahp | Phe | MeTyr | Ile | 0.0025 |
| 2 | Rivulariapeptolide 1185 | Ba-Pro | Ba-Pro | Thr | Thr | Leu | Ahp | Phe | MeTyr | Val | 0.0132 |
| 3 | Molassamide B | Ba | Ala | Thr | Thr | Dhb | Ahp | Phe | MeTyr | Val | 0.0247 |
| 4 | Rivulariapeptolide 1121 | Ba-Pro | Ba-Pro | Thr | Thr | Dhb | Ahp | Leu | MeTyr | Val | 0.0355 |
| 5 | Rivulariapeptolide 1155 | Ba-Pro | Ba-Pro | Thr | Thr | Dhb | Ahp | Phe | MeTyr | Val | 0.0418 |
| 6 | Rivulariapeptolide 988 | | Ba-Pro | Thr | Thr | Dhb | Ahp | Phe | MeTyr | Val | 0.0955 |
| 7 | Crocapeptin A1 | | Pa | Gln | Thr | Leu | Ahp | Phe | MeTyr | Val | 0.1 |
| 8 | Crocapeptin A2 | | iBa | Gln | Thr | Leu | Ahp | Phe | MeTyr | Val | 0.1 |
| 9 | Crocapeptin A3 | | Pea | Gln | Thr | Leu | Ahp | Phe | MeTyr | Val | 0.1 |
| 10 | Bouillomide A | Ba | Ala | Val | Thr | Dhb | Ahp | Phe | MeTyr | Val | 0.17 |
| 11 | Crocapeptin B | | iBa | Gln | Thr | Leu | Ahp | Phe | MeTyr | Ile | 0.2 |
| 12 | Micropeptin KB1046 | | Hpla | Gln | Thr | ThTyr | Ahp | Val | MePhe | Ile | 0.22 |
| 13 | Symplostatin 10 | | Msg | Ile | Thr | Dhb | Ahp | Phe | MeTyr | Ile | 0.222 |
| 14 | Molassamide (others) | Ba | Ala | Thr | Thr | Dhb | Ahp | Phe | MeTyr | Val | 0.234 |
| 15 | Loggerpeptin A | Ba | Ala | Thr | Thr | Leu | Ahp | Phe | Dmy | Val | 0.24 |
| 16 | Cyanopeptolin CP1018 | | Oa | Asp | Thr | Arg | Ahp | Phe | MePhe | Val | 0.24 |
| 17 | Cyanopeptolin CP992 | | Ba | Asp | Thr | Arg | Ahp | Phe | MeHty | Val | 0.24 |
| 18 | Cyanopeptolin CP1027 | | Ha | Asp | Thr | Tyr | Ahp | Phe | MeHty | Val | 0.26 |
| 19 | Cyanopeptolin CP978 | | Ba | Asp | Thr | Arg | Ahp | Phe | MeTyr | Val | 0.26 |
| 20 | Micropeptin KT1042 | | Hpla | Gln | Thr | Tyr | Ahp | Ile | MePhe | Val | 0.26 |
| 21 | Symplostatin 8 | | Msg | Val | Thr | Dhb | Ahp | Phe | MeTyr | Ile | 0.268 |
| 22 | Lyngbyastatin 4 | Hsg | Ala | Hty | Thr | Dhb | Ahp | Phe | MeTyr | Val | 0.3 |
| 23 | Symplostatin 6 | | Msg | Val | Thr | Dhb | Ahp | Phe | MePhe | Val | 0.322 |
| 24 | Symplostatin 9 | | Msg | Val | Thr | Dhb | Ahp | Phe | MeTyr | Val | 0.324 |
| 25 | Symplocamide A | | Ba | Gln | Thr | Cit | Ahp | Ile | Dmy(3-Br) | Val | 0.38 |
| 26 | Micropeptin 88-A | | | Glu | Thr | ThTyr | Ahp | Val | MePhe | Ile | 0.4 |

| | | | | | | | | | | | |
|---|---|---|---|---|---|---|---|---|---|---|---|
| 27 | Symplostatin 7 | | Msg | Ile | Thr | Dhb | Ahp | Phe | MePhe | Ile | 0.515 |
| 28 | Tutuilamide C | | MeCmb | Val | Thr | Dhb | Ahp | Phe | MeTyr | Val | 0.542 |
| 29 | Tutuilamide B | | Cmb | Val | Thr | Dhb | Ahp | Phe | MeTyr | Val | 0.577 |
| 30 | Micropeptin KB1048 | | Ha | Asp(Me) | Thr | Arg | Amp | Ile | MeTyr(3-Cl) | Ile | 0.63 |
| 31 | Microcystin 996 | | Ba | Gln | Thr | Hty | Ahp | Phe | MePhe | Val | 0.64 |
| 32 | Planktopeptin BL1125 | Gla | Hty | Gln | Thr | Leu | Ahp | Thr | Dmy | Ile | 0.8 |
| 33 | Molassamide (this study) | Ba | Ala | Thr | Thr | Dhb | Ahp | Phe | MeTyr | Val | 0.862 |
| 34 | Micropeptin KB991 | | Hpla | Gln | Thr | Leu | Amp | Ile | MeTyr(3-Cl) | Val | 0.87 |
| 35 | Cyanopeptolin 963A | | Ha | Asp | Thr | Tyr | Ahp | Leu | MePhe | Val | 0.9 |
| 36 | Largamide D oxazolidine | Gla-Ahppa | Ala | Val | Thr | Leu | Ahp[a] | allo-Thr | MeTyr(3-Br) | Val | 0.928 |
| 37 | Tutuilamide A | | Cmb | Ile | Thr | Dhb | Ahp | Phe | MeTyr | Val | 1 |
| 38 | Micropeptin 103 | Ha | Gly | Thr | Thr | Gln | Ahp | Phe | MeTrp | Val | 1 |
| 39 | Micropeptin E | | Ha | Glu | Thr | Tyr | Ahp | Leu | MeTyr | Val | 1 |
| 40 | Dinghupepeptin B | | Mba | Gln | Thr | NHeGln | Amp | Phe | MeTyr | Ala | 1.1 |
| 41 | Micropeptin C | | Ha | Glu | Thr | Tyr | Ahp | Phe | MeTyr | Val | 1.1 |
| 42 | Micropeptin LH1021 | | Ha | Thr | Thr | Gln | Ahp | Phe | MeTyr | Val | 1.1 |
| 43 | Micropeptin D | | Ha | Glu | Thr | Tyr | Ahp | Leu | MeTyr | Val | 1.2 |
| 44 | Micropeptin 88-Y | Ac | Tyr | Glue | Thr | Tyr | Ahp | Val | MePhe | Ile | 1.3 |
| 45 | Micropeptin MM836 | | | Gla | Thr | Leu | Ahp | Phe | MePhe | Ile | 1.4 |
| 46 | Jizanpeptin C | | Msg | Val | Thr | Lys | Ahp | allo-Ile | Dmy(3-Br) | Ile | 1.4 |
| 47 | Micropeptin F | | Oa | Glu | Thr | Tyr | Ahp | Leu | MeTyr | Val | 1.5 |
| 48 | Nostopeptin B | | Ac | Gln | 2S,3R,4R-Hmp | Leu | Ahp | Ile | MeTyr | Ile | 1.6 |
| 49 | Micropeptin DR1056 | | Hpla | Gln | Thr | Tyr | Ahp | Leu | MePhe | Ile | 1.6 |
| 50 | Micropeptin MM850 | | | Gla | Thr | Leu | Amp | Phe | MePhe | Ile | 1.7 |

Abbreviations for amino acids/residues (AA) found in Ahp-cyclodepsipetides: Ac: acetic acid; Ahppa: 2-amino-5-(4'-methoxyphenyl)pentanoic acid; Ba: butyric acid; Dmy: *N,O*-dimethyltyrosine; Gla: Glyceric acid; Ha: hexanoic acid; Hpla: 2-Hydroxy-3-(4-hydroxyphenyl)propanoic acid; Hpg: 3'-*O*-phosphate (R)-glyceric acid; Hsg: 3-O-sulfated (R)-glyceric acid; Hty: Homotyrosine; iBa: iso-Butyric acid; Mba: methyl-2-butenoic acid; MePhe:(3',4'-OH): 3',4'-Hydroxy-*N*-methylphenylalanine; Msg: 2-*O*-methyl-3-*O*-sulfate-(R)-glyceric acid; Oa: Octanoic acid; Pa: Propionic acid; Pea:2-pentanic acid; ThTyr: Tetrahydrotyrosine**.**
[a]OH-Group is linked to the Thr-subsite (AA 7 ) and forms an oxazolidine.
